# Rapid evolution can select for fitness tradeoffs in fluctuating environments

**DOI:** 10.64898/2026.09.01.748641

**Authors:** John McEnany, Benjamin H. Good

**Author notes:** Correspondence should be addressed to: B.H.G.

## Abstract

In fluctuating environments, the fates of new mutations depend on the fitness tradeoffs they experience across multiple environmental conditions. Selection on these tradeoffs is well-understood when mutations compete in isolation, but much less is known in the empirically relevant case where multiple mutations compete at the same time. Here, we develop a theory to predict how rapidly adapting populations select on fitness tradeoffs at many linked genetic loci. We derive analytical expressions showing how the fixation probabilities of these mutations depend on their underlying fitness tradeoffs, the timescales of environmental variation, and the future mutations they produce over time. We find that in large populations, competition between linked mutations can strongly favor mutations that carry larger fitness tradeoffs, even when they have a lower geometric mean fitness. We show that this “specialist advantage” arises because transient benefits enhance the rate of producing future mutations – an effect that continues to compound over time even when local conditions shift. We also demonstrate that successful lineages acquire well-timed mutations that appear to anticipate future environments, which can boost the rate of adaptation. These results show how large populations balance competing demands in temporally fluctuating environments, and provide a baseline for interpreting fitness tradeoffs in many natural and experimental settings.

## Introduction

Time-varying environments create a dilemma for natural selection: mutations that are beneficial in one environment often carry costs or tradeoffs in other settings (1). These tradeoffs can arise in diverse contexts (1–7), from seasonal variations in temperature (2) and resources (3), to the evolution of antibiotic resistance (4, 5). They can also arise from more subtle environmental shifts that preserve the overall direction of selection (7). Experimental advances have made it possible to map these fitness tradeoffs with increasing scale and precision, yielding joint fitness measurements for thousands of genetic variants across dozens of environmental conditions (4, 7–15). Yet despite this progress, it remains difficult to predict how evolution chooses between mutations with different tradeoffs when the environment varies over time.

There are two limiting cases where the effects of fluctuating environments are particularly well understood. When the environment changes more rapidly than the timescale of natural selection, the environmental fluctuations self-average, and the long-term dynamics are primarily determined by the time-averaged geometric mean fitness (16–21). Conversely, when the environment changes more slowly than the time it takes for new mutations to reach fixation, their fates will be determined by a constant local environment (20, 21), and any tradeoffs will accumulate nearly neutrally (5). However, there is also a third regime that lies between these two extreme limits, where the effects of time-varying environments become significantly more complex (21). This intermediate regime emerges whenever local environmental perturbations can generate large shifts in mutation frequencies, but the time to fixation remains long enough that mutational tradeoffs are eventually exposed. Empirical estimates suggest that this “strong tradeoff” regime is particularly relevant for microbes and other unicellular organisms, due to their fast generation times and large population sizes. In these rapidly evolving populations, even subtle shifts in the environment can generate local large fitness differences that act over days and weeks (7, 10, 11, 22), yet the size of the population is large enough that mutations take a long time to fix (23).

Previous work has explored these dynamics when mutations compete in isolation (21). However, in many empirical contexts, from laboratory evolution experiments (24–27) to natural populations of viruses (28–30), bacteria (11, 31–34) and certain cancers (35, 36), the supply of new mutations is large enough that multiple mutations will often arise and compete with each other at the same time. The competition between these linked mutations (“clonal interference;” (37, 38)) ensures that the fates of new mutations will not only depend on their own fitness tradeoffs, but also on the tradeoffs of the mutations that they happen to be linked to. This allows for qualitatively new phenomena that are absent from existing models of selection in time-varying environments. For example, mutant lineages can be rescued from unfavorable environments by acquiring further mutations that alter their net tradeoff profiles. Conversely, large populations can harbor “pre-adapted” lineages that are able to expand when the right conditions arise. These arguments suggest that the effects of linked selection may play a critical role in shaping which combinations of tradeoffs will eventually fix.

Here, we address this scenario by developing a mathematical model of rapid adaptation in a temporally varying environment. We focus on a simple and well-studied case, where the environment periodically fluctuates between a pair of environmental states. Within this setting, we develop an analytical approach to predict how the fixation probabilities of new mutations depend on the mutational landscape of their competitors, as well as the statistics of environmental change. We show that adaptive mutations with larger fitness tradeoffs have an advantage in founding further mutations, which can dramatically enhance their fixation probability even when they have a lower geometric mean fitness. We also show that when fitness tradeoffs are common, successful lineages tend to acquire mutations that (by chance) “anticipate” upcoming environmental changes, accelerating the adaptation of the population as a whole. These results suggest that large populations can exhibit unique signatures of adaptation in temporally fluctuating environments, where average fitness alone does not necessarily predict evolutionary success.

## Results

### Rapid evolution favors mutations with environmental tradeoffs

To study how environmental tradeoffs influence adaptation in time-varying environments, we focus on a simple setting where the environment periodically fluctuates between a pair of environmental states (Fig. 1A). Within this context, we consider a large, well-mixed population of *N* haploid individuals, which can acquire new mutations at a large number of linked genetic loci. The effect of each mutation can then be described by its incremental (log) fitness effect in each of the two conditions (*s*_1_ and *s*_2_). The collection of all such mutations defines a corresponding joint distribution of fitness effects (JDFE, Fig. 1B), which plays an analogous role to the classical DFE in a multi-environment setting. Generalist mutations that are independent of the environment will fall on the diagonal of this distribution (*s*_1_ = *s*_2_), while environment-sensitive mutations will generally have an additional off-diagonal component Δ*s* ≡ (*s*_1_ − *s*_2_)*/*2.

**Figure 1.**
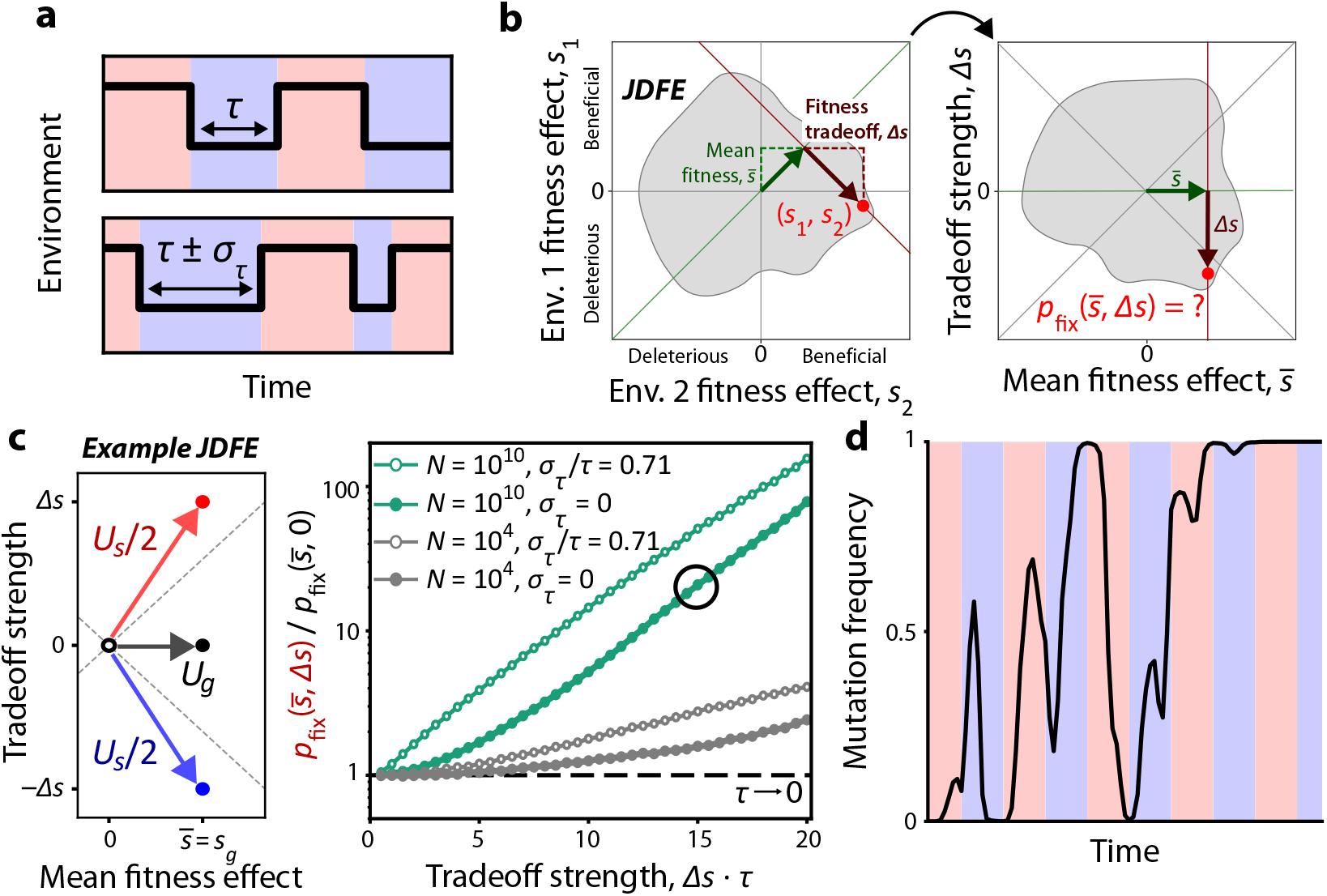
Mutations with tradeoffs are favored in a rapidly adapting population. **(a)** We consider a two-state environment where each state lasts *τ* generations. The switching can either be perfectly periodic (top) or have stochasticity *σ*_*τ*_ (bottom). **(b)** The mutational landscape in a two-state environment can be described by a two-dimensional joint distribution of fitness effects (JDFE). We reparametrize these fitness effects into geometric mean fitness 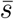 and tradeoff strength Δ*s*, then aim to calculate the fixation probability as a function of these parameters. **(c)** Left: An example coarse-grained JDFE where mutations are grouped into generalist and specialist types that arise at rates *U*_*g*_ and *U*_*s*_, respectively. Generalist mutations have the same benefit *s*_*g*_ in each environment. Specialist mutations have a geometric mean fitness effect 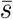 (which in this case, is equal to *s*_*g*_) and an additional tradeoff Δ*s*. Right: We simulated evolution under this JDFE, and measured the specialist advantage 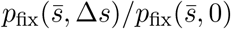 as a function of tradeoff strength Δ*s*. Results are shown for the rapid adaptation regime (*N* = 10^10^) and the successive mutations regime (*N* = 10^4^), at different values of switching stochasticity *σ*_*τ*_. Dashed line shows the *τ* → 0 prediction where the fixation probability depends only on 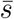. Simulations were run with 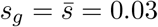, *U*_*g*_ = *U*_*s*_ = 3 · 10^−6^, and *τ* = 100. **(d)** Sample trajectory of a successful specialist mutation (favoring the red environment) for the parameters corresponding to the circled point in (c).

Previous work has shown that when the environment fluctuates sufficiently rapidly, the fates of each mutation will be determined by their geometric mean fitness effects,

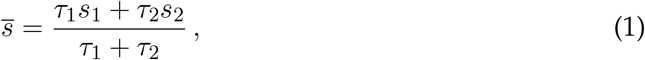

where *τ*_1_ and *τ*_2_ denote the average time spent in each of the two conditions (Fig. 1B). This suggests a useful reparameterization of the JDFE for a given evolution environment, in which each mutation is characterized by its mean fitness 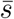 and its corresponding tradeoff Δ*s*. Generalist mutations will lie on the x-axis of this distribution (Δ*s* = 0), while environment-sensitive mutations will fall either above or below this line (Δ*s* ≠ 0). In the main text, we will generally focus on the symmetric case where *τ* ≡ *τ*_1_ = *τ*_2_, but this simplification is not critical for our results (SI 2.1.5).

Following previous work (21), we will quantify the emergent selection pressures on these mutations by considering their long-term fixation probabilities 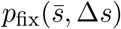; this provides a natural measure of the strength of selection when many different mutations are available (39, 40). A central goal of this study will be to determine how these fixation probabilities depend on the underlying fitness tradeoffs and the timescales of environmental variation.

To address this question, we begin by considering a coarse-grained version of the twodimensional JDFE in Fig. 1B, which contains three different types of mutations (Fig. 1C). Generalist mutations occur at a total rate *U*_*g*_, and provide an average fitness benefit that is independent of the environment (*s*_*g*_ ≡ *s*_1_ = *s*_2_). We also consider a second class of environment-dependent “specialist” mutations; these occur at a total rate *U*_*s*_, and confer an average benefit 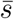 as well as a corresponding tradeoff ±Δ*s* (Fig. 1C). For simplicity, we work in the infinite sites limit and neglect the effects of epistasis, so that the JDFE remains constant as the population adapts. These assumptions define a minimal model for studying the effects of fitness tradeoffs on a fluctuating fitness “seascape.”

Figure 1C shows the measured fixation probabilities in simulations of this model (SI 4.1) for several different population sizes and switching conditions. In previous single-locus models (21), the fixation probabilities of beneficial generalists often exceed their specialist counterparts, until the latter start to have an appreciable chance of fixing in a single environment. In contrast, the simulations in Fig. 1C show that the opposite behavior occurs in large populations where multiple mutations compete at the same time (*NU*_tot_ ≫ 1, teal curves). In these settings, the fixation probability of an environment-sensitive mutation can be ~10-100x larger than a corresponding generalist mutation with the same geometric mean fitness. These advantages are largest for strong fitness tradeoffs (Δ*s* · *τ* ≫ 1) and vanish when Δ*s* · *τ* ≪ 1. When environmental switching is stochastic, the benefits of specialism grow even larger (Fig. 1C).

This behavior is counterintuitive under existing theories that emphasize the (realized) time-averaged fitness. As the population size becomes larger, new mutations take longer to reach fixation, and the examples in Fig. 1C experience multiple environmental cycles before they are able to fix. During this time, successful mutations spend a roughly equal time in both favorable and unfavorable environments, and their trajectories fluctuate strongly due to environmental change and further evolution in the population (Fig. 1D). But rather than averaging out (as in the single-locus case), these transient fluctuations seem to provide a strong systematic advantage to mutations with strong tradeoffs. How does this “specialist advantage” emerge when multiple mutations compete for fixation?

### Modeling rare mutations with tradeoffs in a perfectly periodic environment

To understand the origin of the specialist advantage in Fig. 1, it is useful to begin by considering a simpler case, where specialist mutations are rare (*U*_*s*_ → 0) but generalist mutations are still common (*NU*_*g*_ ≫ 1). In this case, we can consider the fates of individual specialist mutations as they compete against a diverse generalist background (Fig. 2A). Crucially, both the specialist strain and the background population will continue to acquire generalist mutations, but their environmental tradeoffs will remain fixed as the environment-sensitive mutant competes for fixation.

**Figure 2.**
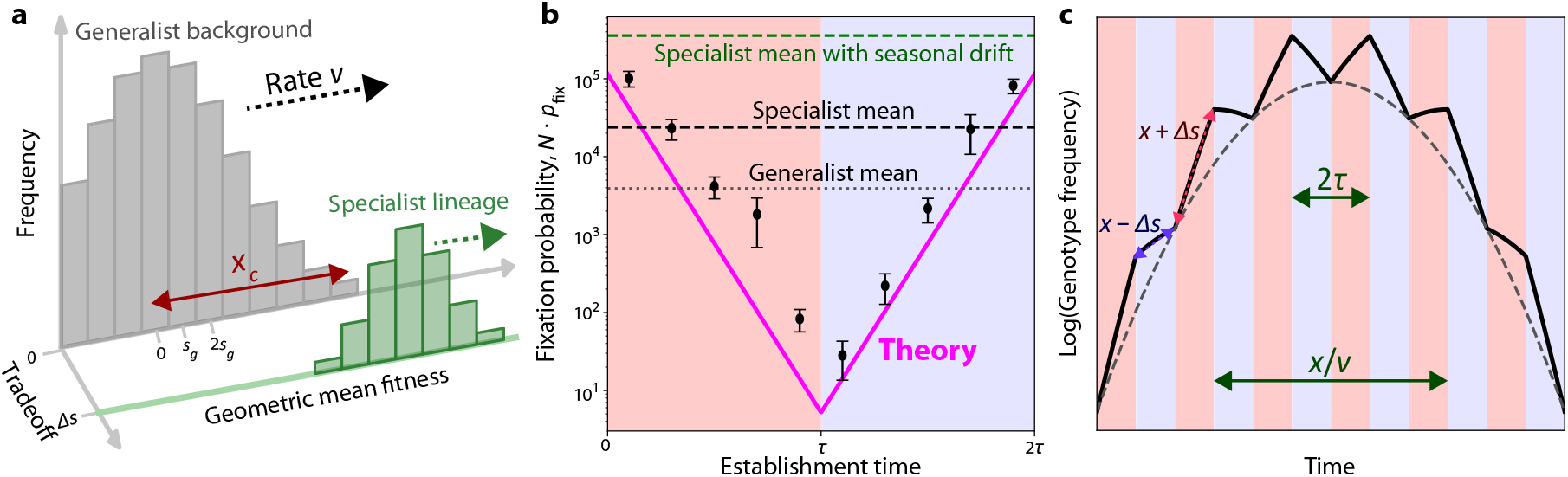
Traveling fitness waves with rare environmental tradeoffs. **(a)** Schematic of the two-dimensional fitness distribution of a population with rare specialist mutations. The background population forms a traveling wave (gray) insensitive to environmental change, increasing its fitness at rate *v* by acquiring generalist mutations with fitness effect *s*_*g*_. A small specialist lineage (green) descends from a mutation with an environmental tradeoff Δ*s*, and can acquire further generalist mutations that advance its geometric mean fitness. **(b)** The fixation probability of a rare specialist mutation as a function of the time when it arises, within a beneficial (red) or deleterious (blue) environment. Dashed line shows the specialist fixation probability averaged over time while dotted line shows the fixation probability of a generalist mutation. The green line shows the fixation probability of the same specialist when the environment has a modest amount of seasonal drift, *σ*_*τ*_ */τ* = 0.35. Pink line is the theoretical prediction from Eq. (9). Error bars show SEM (see SI 4.2). Simulations were run with *N* = 10^10^, 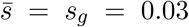, Δ*s* = 0.1, *U*_*g*_ = 3 · 10^−6^, and *τ* = 100. **(c)** A schematic of the expected trajectory of a specialist genotype that arises with geometric mean fitness *x*, in an environment which fluctuates with timescale *τ* (shading). Over timescales ~ *x/v*, this genotype will be outcompeted by fitter ones, which may be this genotype’s descendants or those of another strain in the background population. The dashed line shows the trajectory of an environment-insensitive genotype with the same geometric mean fitness.

Simulations confirm that as before, rare environment-sensitive mutations have a marked fixation advantage over generalists. Intriguingly, this advantage is strongly dependent on when the specialist mutation arises: the fixation probability is maximized for mutations which establish near the start of a beneficial epoch, and decreases exponentially for later times (Fig. 2B). The time-averaged fixation probability is dominated by mutations that establish near the optimal time, at the peak of the two-sided exponential. In order to account for this time dependence, we will need a method to analyze competition between strains.

A useful approach is to construct a fitness distribution by sorting individuals into fitness classes, based on their beneficial mutation count. In the absence of the specialist strain, past work (41) has shown this fitness distribution forms a “traveling wave” which advances at the rate of adaptation 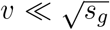 (Fig. 2A). Evolution in this regime is driven by mutations in the high-fitness individuals at the “nose” of the wave, whose fitness is *x*_*c*_ ≫ *s*_*g*_ higher than the population mean. These results apply for a broad range of parameters applicable to large microbial communities, 1 ≪ *NU*_*g*_ ≪ *Ns*_*g*_ (SI 1.1). Importantly, these traveling wave results will still hold in the presence of a low-frequency specialist strain, allowing us to express *v* and *x*_*c*_ in terms of other evolutionary parameters (SI 1.1). Henceforth, we will make the additional assumption that Δ*s* ≪ *x*_*c*_, so the tradeoff strength is not so strong that it overpowers all other fitness variation in the population. This assumption does not limit the relative size of 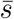 and Δ*s*, so specialist mutations can still have strong tradeoffs relative to their average benefit.

If a specialist mutation arises on a high-fitness background close to the nose of the traveling wave, it will begin rising in frequency deterministically as long as it escapes initial genetic drift. The instantaneous growth rate of this genotype depends on its relative fitness, which changes whenever the environmental state does. In addition, this relative fitness will experience an overall decline at rate *v*, caused by adaptation in the rest of the population. When the focal genotype’s fitness is surpassed by the bulk population’s, it will begin falling in frequency, forming a parabolic log frequency trajectory punctuated by environmental change (Fig. 2C). Thus, the specialist (or indeed, any) mutation can only survive if its offspring produce their own beneficial mutations, keeping up with adaptation in the rest of the population even as each individual genotype in the lineage goes extinct. The specialist advantage, and its dependence on establishment time, can be explained through this process of founding future mutations.

#### Rolling fitness of a lineage

To study the dynamics of how an environment-sensitive lineage founds new beneficial mutations, we will extend the successive establishment picture analyzed in Ref. (42) to the case of fluctuating environments. We consider a new specialist mutation which establishes at time *t* on a high-fitness genetic background, such that its relative geometric mean fitness is initially *x* ~ *x*_*c*_. The specialist genotype will rise in frequency deterministically, eventually producing a new generalist mutation that survives initial genetic drift. Crucially, the descendants of this mutation will effectively act like a new specialist lineage which establishes at shifted relative average fitness *x*′ and time *t*′, given by

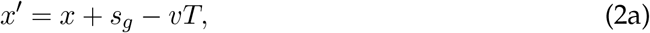

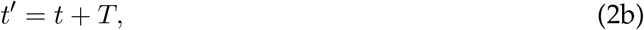

where *T* represents the number of generations between the establishment of the initial lineage (at average fitness *x*) and its offspring (at *x*′). Previous work (42) demonstrates that the fixation probability of the initial lineage can be approximated by solving the recursive “rolling fitness” relations in Eq. (2) over a large number of founding steps – predicting whether the descendants of a lineage will tend to increase their fitness until they reach fixation, or be outcompeted by the bulk population and go extinct.

All the complexities of growth in a changing environment have been absorbed into *T*, the waiting time for the next mutation. While *T* is a random variable, it is sufficient for our purposes to consider the *typical* time it takes the lineage to establish its first mutation, which we will also denote as *T* for convenience. It is most convenient to write the solution for *T* under the additional assumption that *τ* ≫ *s*_*g*_*/v*, in which case *t* and *t*′ will usually be in the same environmental state given by the indicator function *I*(*t*) = ±1. In this case, *T* takes the form

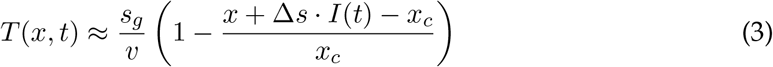

to leading order in (*x − x*_*c*_)*/x*_*c*_ and Δ*s/x*_*c*_ (SI 2.1.1). Eq. (3) indicates that mutations in a beneficial environment with *I*(*t*) = +1 rise in frequency faster and found beneficial mutations sooner (Fig. 2C). A similar effect is achieved if the mutation starts on a higher-fitness background with larger *x*. Meanwhile, Eq. (2) shows that a mutation which arises sooner (smaller *T*) lands at a higher rolling fitness *x*′ because the rest of the population has had less time to adapt. Crucially, these effects compound over successive foundings, such that small differences in initial fitness increase over time:

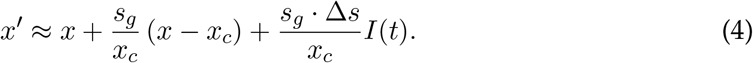

The recursive solution to Eq. (4) is particularly simple for generalist lineages with Δ*s* = 0, where mutations that start above the “break-even fitness” *x*_*c*_ will tend to fix deterministically, while those below *x*_*c*_ will be likely to go extinct (42). Specialist lineages experience a similar compounding effect, but are also sensitive to the environmental state *I*(*t*) during each founding step. Thus, their rolling fitness after *n* successive foundings can be written as a weighted sum of the environmental states they experienced during each step:

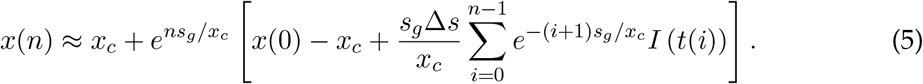

The influence of the environment in early steps soon after the specialist arises (smaller *i*) has more time to compound over future foundings, so those terms have the most weight in the sum (SI 2.1.1). Many cycles after the specialist mutation was founded (*n* ≫ *x*_*c*_*/s*_*g*_), we can exploit the periodic structure of *I*(*t*) to simplify the sum in Eq. (5) further:

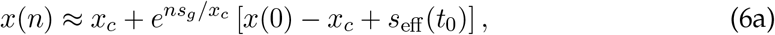

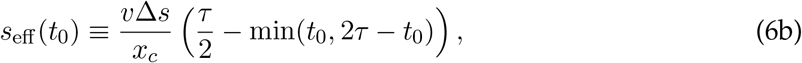

where *t*_0_ is defined such that the environment is beneficial between *t*_0_ = 0 and *t*_0_ = *τ*, and deleterious from *t*_0_ = *τ* to 2*τ*. This relation can also be shown to hold in the faster-cycling *τ* ≪ *s*_*g*_*/v* regime, where individual foundings span multiple environmental epochs (SI 2.1.1).

Eq. (6a) indicates that the long-term fixation probability of a specialist mutation 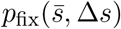 coincides with that of a generalist mutation with shifted fitness effect 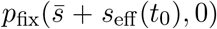. The shift *s*_eff_ (*t*_0_) can therefore be interpreted as an effective fitness boost experienced by lineages with environmental tradeoffs, depending on the time *t*_0_ they arise. This shift is positive if *t*_0_ is near the start of a beneficial epoch, and negative if *t*_0_ is near the start of a deleterious epoch, switching sign halfway through each environmental state. This behavior is consistent with the observed time dependence in Fig. 2A and the weighted sum in Eq. (5), favoring mutations which experience a full beneficial epoch immediately after they arise. Thus, even after many cycles, the time the lineage initially arose strongly influences its fate.

A specialist mutation which arises at the optimal time will have *s*_eff_ = Δ*s* · *vτ/*2*x*_*c*_, much smaller than the direct benefit Δ*s* it would experience in a permanent favorable environment. However, this boost will still have a large impact on the mutation’s fixation probability, which we discuss next.

#### Fixation probability of a general mutation

The overall fixation probability can be calculated by marginalizing over the fitness of the possible genetic backgrounds the mutation could arise on:

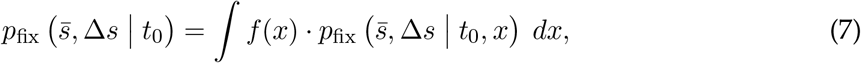

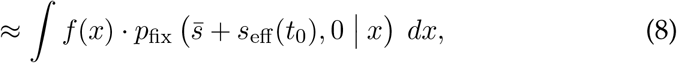

where 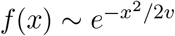 is the background fitness distribution in Eq. (S4). Past work (39) has shown that the integral in Eq. (8) is dominated by backgrounds where 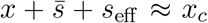, because fixation is strongly suppressed at lower fitnesses, but higher-fitness backgrounds are much rarer in the population. Therefore, a well-timed specialist mutation (with larger *s*_eff_) can reach fixation on slightly lower-fitness backgrounds (smaller *x*; SI 2.1.4). Because these lower-fitness backgrounds are exponentially more frequent in the population, the overall fixation probability has a significant dependence on establishment time:

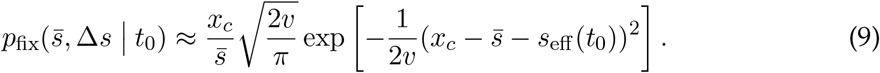

Since rare environment-sensitive mutations arise uniformly over time, integrating over *t*_0_ will yield the overall fixation probability. This average fixation probability is dominated by mutants that arise near the start of a beneficial environment, yielding

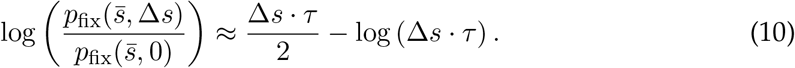

Thus, environment-sensitive mutations have an exponentially enhanced fixation probability scaling with Δ*s* · *τ* (Fig. 3A), systematically outcompeting generalist mutations with the same geometric mean fitness. This result holds when 1*/*Δ*s* ≪ *τ* ≪ *x*_*c*_*/v*, corresponding to the strong-tradeoffs regime where the environment changes many times before the mutant reaches fixation. A similar result applies in the case of asymmetric environments with *τ*_1_≠ *τ*_2_, where the harmonic mean of *τ*_1_ and *τ*_2_ becomes the relevant timescale (SI 2.1.5).

**Figure 3.**
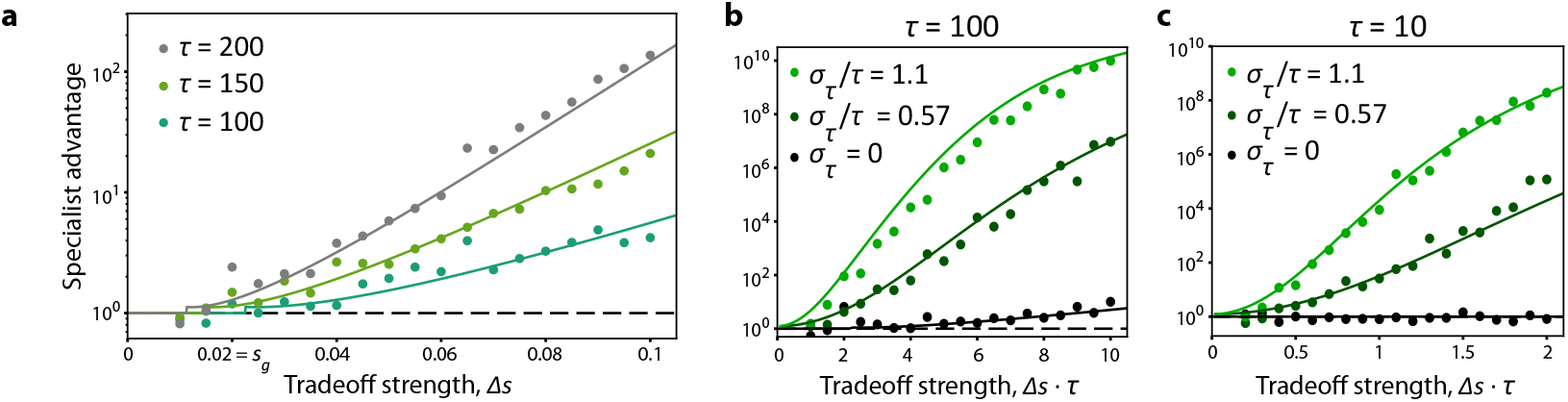
Fixation probabilities of rare mutations with environmental tradeoffs. **(a)** Fixation advantage of rare environment-sensitive specialists, 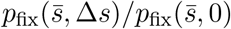, in a perfectly periodic environment (*σ*_*τ*_ = 0) for different environmental fluctuation times *τ* and tradeoff strengths Δ*s*. Curves show theory prediction from Eq. (S31) and Eq. (S32), and unity for Δ*s* · *τ <* 2.25 (where the specialist advantage is negligible). **(b)** The specialist advantage in a stochastic environment with *τ* = 100, for different values of *σ*_*τ*_ and Δ*s*. Curves show theory prediction obtained from numerically integrating Eq. (S46) under the normal approximation in Eq. (S47) with no cutoff *c*. **(c)** Same as (b), but in a rapidly fluctuating environment with *τ* = 10. Simulations were run with *N* = 10^20^, 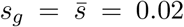 and *U*_*g*_ = 3 · 10^−9^.

### Effects of environmental stochasticity on rare lineages with tradeoffs

Environmental fluctuations are in general not perfectly periodic. We can account for this by modeling the amount of time the environment stays in one state as a random variable with mean *τ* and variance 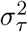, e.g. 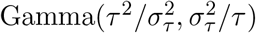, which we use in simulations. The ratio *σ*_*τ*_ */τ* sets the strength of environmental unpredictability or seasonal drift: *σ*_*τ*_ */τ* = 1 represents a memoryless environment which has an equal chance of switching at any time, while *σ*_*τ*_ */τ* ≪ 1 corresponds to a mostly periodic environment with small amounts of stochasticity.

Strikingly, even a modest amount of environmental stochasticity, *σ*_*τ*_ */τ* ≪ 1, increases the fixation probability of rare specialist mutations by an order of magnitude (Fig. 2B). To explain this, we can extend our analysis of successive mutations to account for seasonal drift. A lineage which spends longer in a beneficial environment will grow and found new mutations faster, so the rolling fitness will change stochastically each environmental cycle based on whether the beneficial or deleterious period lasted longer. When the stochasticity is not too strong, we can express its effect in terms of the time differences *δτ* between successive epochs (SI 2.2.1). This results in a modified version of Eq. (6a), where an environment-sensitive lineage experiences an effective fitness shift 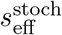 based on the realized stochasticity ***δτ***:

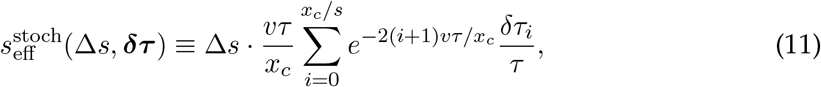

valid in the slow-cycling *τ* ≫ *s*_*g*_*/v* regime (see SI 2.2.1 for the more general case). This is a weighted sum of the environmental stochasticity, with more weight placed on earlier cycles (smaller *i*) – similar to the weighted sum over environments in Eq. (5), where a bias toward beneficial environments soon after the lineage establishes acts like a long-term fitness benefit. Note that in principle, the true *s*_eff_ in a stochastic environment should be some combination of the deterministic shift in Eq. (6b) and the stochastic shift in Eq. (11); here, we focus on the regime where the stochastic part dominates.

The overall fixation probability now requires marginalizing over genetic backgrounds and environmental stochasticity,

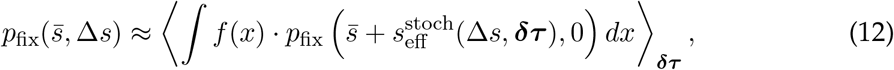

where the interior integral will once again be dominated by genetic backgrounds near 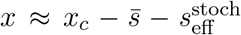. To average over environmental stochasticity, it is reasonable to invoke the central limit theorem if the underlying distribution of *δτ* is not too wide-tailed, approximating 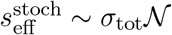 with *N* a standard normal random variable and

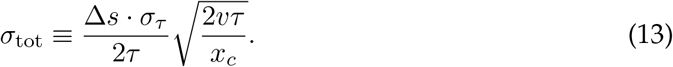

Numerically solving Eq. (12) under this normal approximation recapitulates the extreme specialist advantage observed in simulations (Fig. 3B-C), where specialist mutations can even be 10^10^ more times likely to fix as generalists. This is because lucky sequences of environmental fluctuations give environment-sensitive lineages on lower-fitness backgrounds a viable path to fixation, significantly widening the range of lineages that can fix. For modest environmental fluctuations 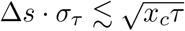, the specialist advantage is

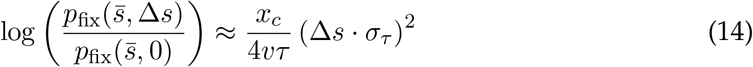

to leading order. Because the *x*_*c*_*/vτ* prefactor is large, even weak environmental stochasticity (Δ*s* · *σ*_*τ*_ ≪ 1) can significantly influence fixation probabilities.

The expectation in Eq. (12) can be dominated by rare environmental fluctuations strongly biased toward beneficial environments, which may not be observed in practice (SI 2.2.1). However, these rare events are not critical for our qualitative results, because even typical realizations of environmental stochasticity continue to provide a large advantage to mutations with tradeoffs (Fig. S1). Another possibility is that the distribution of *δτ* is wide-tailed enough for the normal approximation to break down, in which case the fixation probability can be dominated by long beneficial epochs where the mutation appears to provide an unconditional benefit of 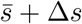 (Fig. S2; SI 2.2.2). More generally, Eq. (12) still applies for more complicated underlying distributions of ***δτ***, such as correlated epoch durations (43). These details do not change the overall picture that small levels of environmental stochasticity can provide huge benefits for a rare lineage sensitive to environmental change.

Note that the effects of stochasticity discussed in this section and Figure 3B-C are much larger than those observed in Figure 1C. The reason for this difference is our assumption that specialist mutations are rare: in this case, only one lineage at a time can benefit from environmental stochasticity, drastically increasing its fixation probability. When specialist mutations are widely available, this will no longer be the case, which we discuss later.

### Frequent mutations with tradeoffs in a periodic environment

Generally, mutations with substantially different fitness effects between environments are not always rare relative to generalist mutations. When specialist mutations are common (*NU*_*s*_ ~ *NU*_*g*_ ≫ 1), then the dynamics of the entire population will be reshaped by environmental change, so the one-dimensional “traveling wave” results we employed in the rare-specialists regime will no longer apply. By extending our earlier calculations to the frequent-specialists regime, we will confirm that tradeoffs continue to be favored even when they are common, and demonstrate their impact on collective properties like the population-wide rate of adaptation. As before, we will start with periodic environments (*σ*_*τ*_ = 0) and then discuss stochastically varying ones.

When mutations with different environmental fitness effects are widely available, the population as a whole will have a fully two-dimensional fitness distribution which is responsive to environmental change. Likewise, the descendants of a focal lineage will follow many different “mutational paths” depending on which types of mutations they acquire at each time. Despite this complexity, we can make progress by realizing that *successful* lineages (those that eventually reach fixation) are likely to follow an optimized mutational path which depends on environmental change. In other words, selection favors the organisms that happened to acquire the right mutations at the right times, even though each individual mutation arises randomly. To uncover the impact of frequent specialist mutations, we will derive this optimal mutational path and self-consistently relate it to the entire fitness distribution of the population.

#### Treatment of frequent tradeoffs

In order to describe competition between a focal lineage and the rest of the population, we must make an ansatz for the dynamics of the bulk population in two-dimensional fitness space. We choose to parametrize fitness in terms of geometric mean fitness *X*_∥_ and environmental “specialism” *X*_⊥_, defined such that a mutation with fitness effect 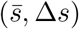 shifts *X*_∥_ by 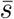 and *X*_⊥_ by Δ*s*. We similarly define relative fitnesses 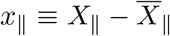 and 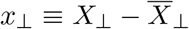, where bars represent the population mean. In the rare-specialists case we studied earlier, the bulk population was insensitive to the environment 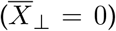 and had an average fitness which increased at rate *v*, such that 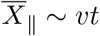. For the frequent-specialists regime, we make a similar linear ansatz:

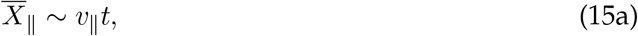

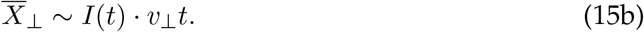

where *I*(*t*) = ±1 once again represents the current environmental state. Under this ansatz, the population zigzags in two-dimensional fitness space, increasing its long-term average fitness at the rate of adaptation *v*_∥_, but also cyclically specializing to the current environment at the “rate of specialization” *v*_⊥_ (Fig. 4A). The structure of this linear ansatz is in good agreement with simulations (Fig. S3A), but both rates *v*_∥_ and *v*_⊥_ must be determined self-consistently from the behavior of individual lineages.

**Figure 4.**
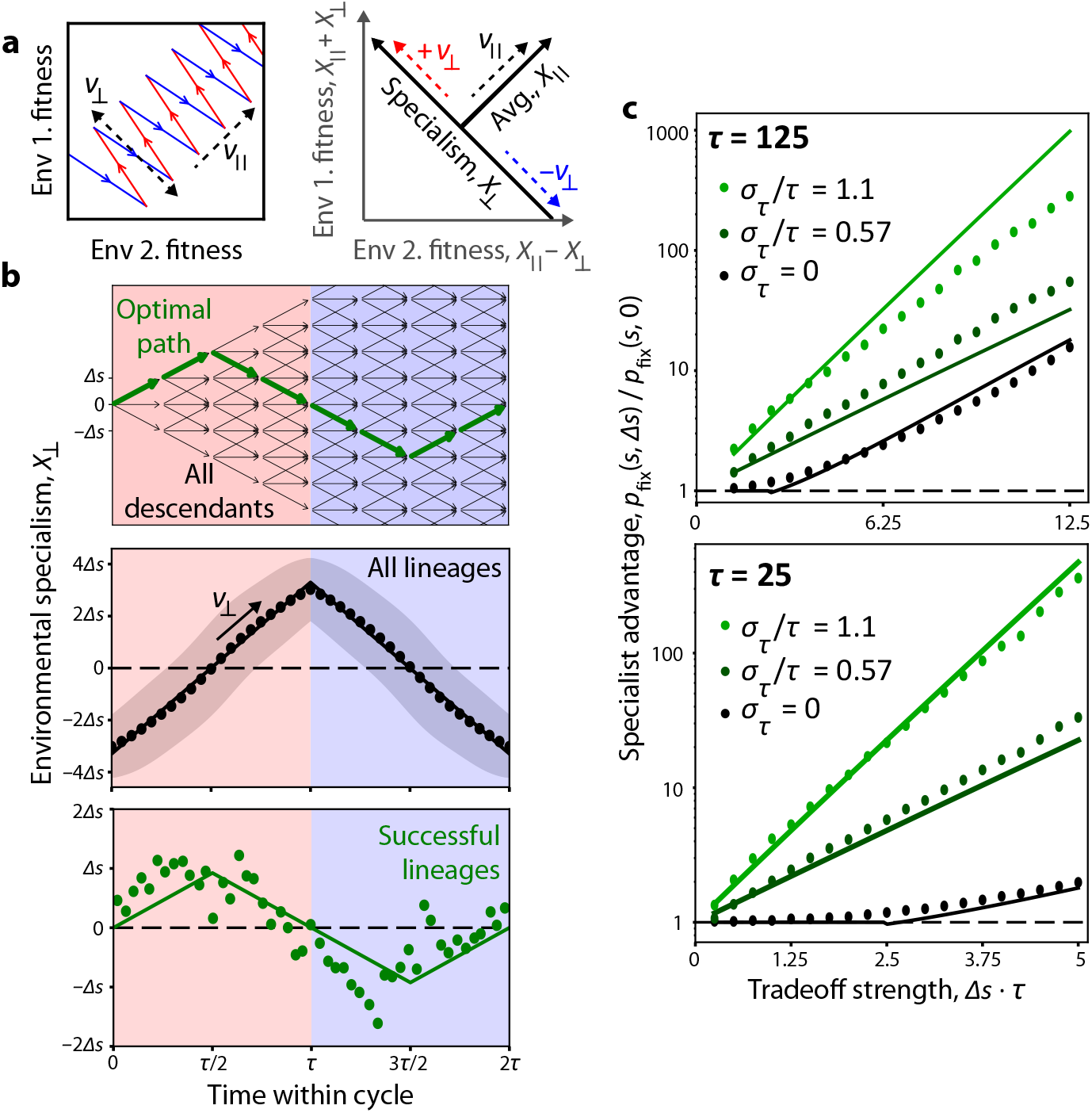
Populations with frequent specialist mutations follow an optimal mutational path. **(a)** Left: When specialist mutations are frequent, the population mean zigzags in two-dimensional fitness space, adapting at rate *v*_∥_ over the long term, but cyclically specializing to the current environment at rate *v*_⊥_. Right: We parametrize fitness as average fitness *X*_∥_ and environmental specialism *X*_⊥_, corresponding to *v*_∥_ and *v*_⊥_. **(b)** Top: Schematic of the environmental specialism of the members of a lineage, with mutations represented by arrows. The descendants with the largest long-term growth rate follow the “optimal mutational path” shown by the green arrow. Middle: The average environmental specialism 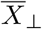 over an environmental cycle. The line is a best-fit line to simulation results (points); the shading shows a standard deviation. Bottom: The empirical environmental specialism *X*_⊥_ of new mutations (accounting for their genetic background) binned by establishment time, conditioned on those mutations later reaching fixation. The line shows a best-fit line consistent with the optimal mutational path. **(c)** Specialist advantage *p*_fix_(*s*, Δ*s*)*/p*_fix_(*s*, 0) of frequent specialist mutations, for various values of Δ*s, τ*, and *σ*_*τ*_. Theory curves show prediction from Eq. (S135) when *σ*_*τ*_ = 0, and from Eq. (S165) when *σ*_*τ*_ *>* 0, using values of *v*_∥_ fit from simulations. Simulations run with *N* = 10^20^, *U*_*g*_ = *U*_*s*_ = 3 · 10^−9^, *s* = 0.02, Δ*s* = 0.05, and *τ* = 200 unless otherwise noted.

To approach this complicated regime in the simplest possible way, we continue to employ our three-class JDFE containing generalist and specialist mutations, and assume that these mutations have the same geometric mean fitness effects 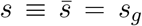 and mutation rates *U*_*g*_ = *U*_*s*_ (see SI 3.2.7 for the asymmetric generalization). We also specialize to the slower-cycling case *τ* = *τ*_1,2_ ≫ *s/v*_∥_. Finally, for the frequent-specialists regime we make a new assumption that *s >* Δ*s*, so specialist mutations are always beneficial regardless of the environment. In the alternate regime Δ*s > s*, many qualitative conclusions will hold (indeed, many of our simulation results are in this regime), but the optimal mutational path changes (SI 3.2.8). Finally, we note that the critical fitness scale *x*_*c*_ can be redefined in terms of *v*_∥_ according to Eq. (S60).

Now, we are equipped to generalize our rolling fitness recursion relations in Eq. (2) to account for acquisition of further specialist mutations:

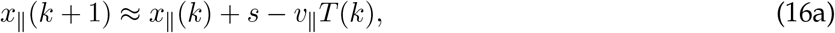

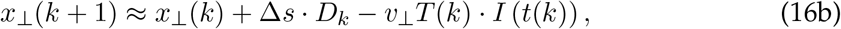

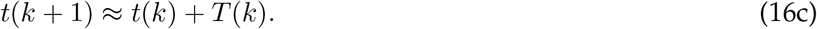

The identity of the mutations the focal lineage acquires is given by *D*_*k*_ ∈ (−1, 0, 1), with 0 representing generalists and ±1 representing each specialist type. These equations are valid when *t*(*k* + 1) and *t*(*k*) are both in the same environmental state, consistent with *τ* ≫ *s/v*_∥_. They reduce to the rare-specialists case with constant *x*_⊥_ when *v*_⊥_ = *D*_*k*_ = 0.

#### Optimal mutational path

Because the waiting time *T* (*k*) depends on *x*_∥_(*k*), *x*_⊥_(*k*) and *t*(*k*), the rolling fitness *x*_∥_(*k*) will depend on the mutational history of the lineage up to that point, i.e. the *D*_*i*_ with *i < k*. We can write a solution for *x*_∥_(*n*) and *x*_⊥_(*n*) that generalizes Eq. (5) to a specializing lineage, valid when the effects of specialism *x*_⊥_ can be treated perturbatively relative to *x*_∥_:

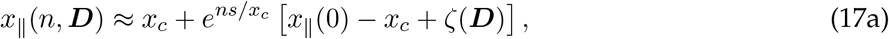

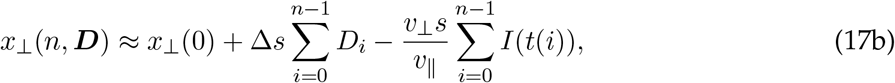

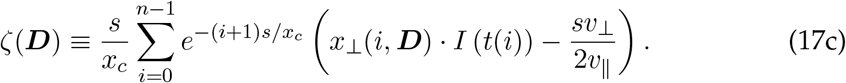

Here, *ζ* is an effective fitness shift due to the net effect of specialization in the focal lineage and the rest of the population, which depends on the mutational path ***D*** through *x*_⊥_(*i*, ***D***). The optimal mutational path ***D***^∗^ maximizes *ζ*(***D***), and can be expressed as the sign of a weighted sum of environmental states after *t*(*j*):

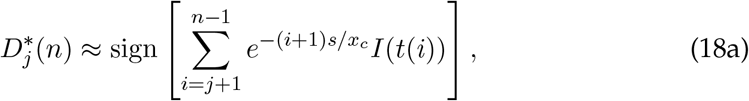

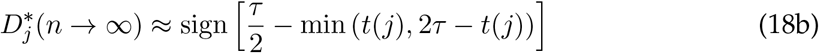

for *n* ≳ *j* + *x*_*c*_*/s*, measuring times modulo 2*τ*. The fitness boost for a lineage following this optimal path is

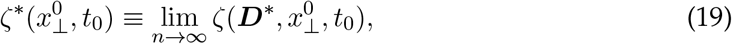

explicitly calculated in Eq. (S68) (SI 3.2.2).

The long-term optimal path in Eq. (18b) exactly matches our intuition from the rare-specialists case in Eq. (6b): specialist mutations aligned with the current environment are optimal only until it is halfway over. Afterward, a lineage should switch to the opposite type of specialist mutation to “pre-adapt” for the upcoming environment. This optimal behavior is phase-shifted by *τ/*2 generations relative to the specialization of the bulk population, which only adapts to the current environment – meaning that successful lineages are significantly more pre-adapted than typical ones (Fig. 4B; Fig. S4). Signatures of this pre-adaptation are also visible in the trajectories of individual mutations (Fig. 1D), where successful lineages often rise in frequency shortly after environmental switches – regardless of which environmental state has just started.

If the symmetry condition is relaxed so that environment-sensitive mutations have a smaller mutation rate or average fitness effect relative to generalist mutations, there will be a period roughly halfway into each environmental state where the optimal mutational path includes generalist mutations with *D* = 0 (SI 3.2.7). Extreme differences in mutation rate of order *U*_*g*_*/U*_*s*_ ~ *e*^Δ*s*·*τ/*2^ mark the transition between the rare-specialist and frequent-specialists regimes; more modest differences will largely be similar to the symmetric case.

#### Fixation probability and specialist advantage

As in Eq. (8), we can estimate the fixation probability of a focal mutation with fitness effect 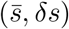 by integrating over the distribution of genetic backgrounds, which is now two-dimensional. (*δs* represents the tradeoff strength of the focal mutation: 0 for a generalist and ±Δ*s* for a typical specialist mutation.)

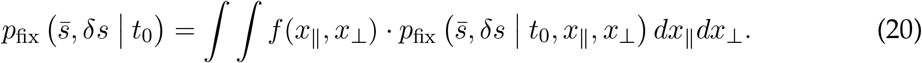

We can calculate the background distribution by estimating the effective fitness cost of deviating from the optimal path (SI 3.2.3), finding a two-dimensional Gaussian (Fig. S5):

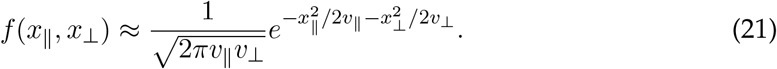

The integral in Eq. (20) will be dominated by a narrow range of genetic backgrounds where the mutation lands at a critical fitness scale, 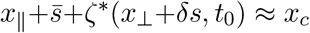 (Fig. S5, black curve); this favors values of *x*_⊥_ which are pre-adapted to the upcoming environment (Fig. S5, pink dots). If the focal mutation is a specialist with *δs* ≠ 0, it effectively shifts *x*_⊥_, allowing the lineage to pre-adapt further. Thus, well-timed specialist mutations have a fixation advantage over generalists for the same reason as before: they provide an effective fitness boost which enables them to fix even on backgrounds with slightly lower average fitness.

The overall time dependence of Eq. (20) is somewhat more complicated than the rare-specialist regime, because the focal mutation must compete against environment-sensitive mutations on other genetic backgrounds (SI 3.2.5). Because these competing mutations will have an advantage if they arise at the start of a favorable environment, generalist mutations with *δs* = 0 will be most likely to fix if they arise near the middle of an environmental state, where specialists have no advantage (Fig. S6A). While this effect attenuates the overall specialist advantage 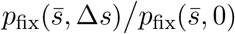 to scale linearly with Δ*s* · *τ* rather than exponentially (Fig. S7A), mutations with tradeoffs are still strongly favored overall.

When tradeoffs are strong enough that specialist mutations appear deleterious in their unfavored environment (Δ*s > s*), the optimal mutational path shifts so that successful lineages *stop* acquiring mutations in the middle of each environmental state, then acquire “bursts” of specialist mutations in rapid succession near environmental switches (SI 3.2.8; Fig. S6B). Because specialist mutations are strongly favored by natural selection at these times, this shift coincides with the specialist advantage growing to scale with exp(Δ*s* · *τ*), as in the rare-specialists regime (Fig. 1C, Fig. 4C, Fig. S7A).

#### Rates of specialization and adaptation

We can estimate the rate of specialization *v*_⊥_ by considering the “short-term” optimal mutational path for mutations shortly before the present (SI 3.2.3), yielding

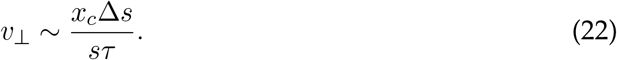

As *τ* increases, the population maintains less steady-state diversity in environmental preference, lowering the rate of specialization *v*_⊥_. This result holds qualitatively in simulations (Fig. S3B), but Eq. (22) tends to overestimate the measured *v*_⊥_ except in extremely large populations where all of our asymptotic assumptions can be strongly satisfied.

We can estimate the long-term rate of adaptation *v*_∥_ through the self-consistent relation

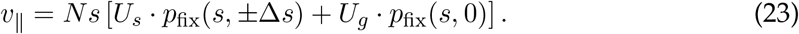

The fitness boost *ζ*^∗^ from following the optimal mutational path will increase *p*_fix_ and therefore *v*_∥_ relative to the rare-specialist regime. To leading order (SI 3.2.6), this boost is

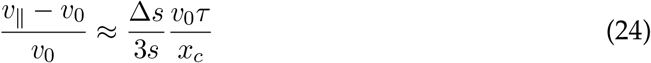

where *v*_0_ corresponds to *v* for a population with rare specialists (or *τ* → 0). This small boost stems from the fact that specializing lineages will rise in frequency more quickly by adapting to the environment, making the rate of adaptation larger than the one predicted by average fitness alone (Fig. 5A). This boost is sensitive to our symmetry assumption that the JDFE contains specialist and generalist mutations with the same average fitness effect *s*. In fact, as we will discuss in the next section, *v*_∥_ can fall below *v*_0_ if specialist mutations provide a smaller average fitness benefit than generalists.

**Figure 5.**
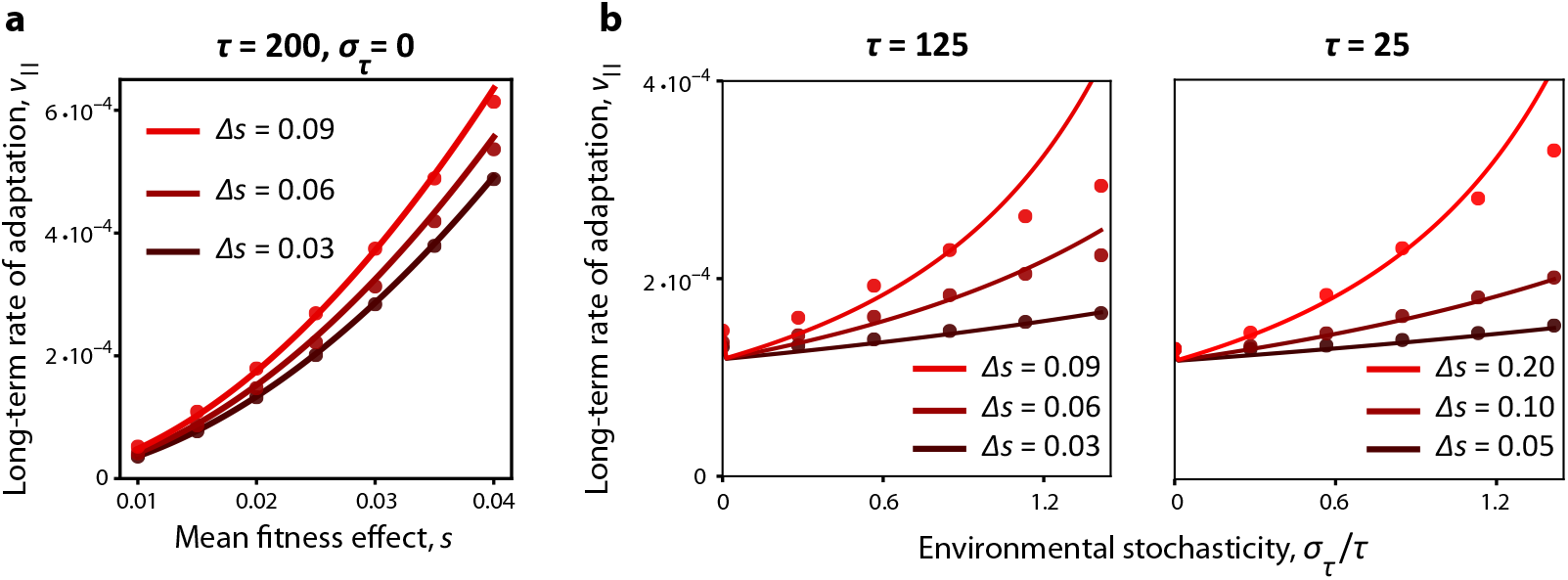
Environmental fluctuations increase the population’s overall rate of adaptation. **(a)** Rate of adaptation *v*_∥_ as a function of the mean fitness benefit *s* of mutations, for different specialist environmental tradeoffs Δ*s* in a deterministically switching environment (*σ*_*τ*_ = 0). Curves show theory prediction, numerically solved from Eq. (S106). **(b)** The rate of adaptation as a function of environmental stochasticity *σ*_*τ*_ for different values of Δ*s* and *τ*. Curves show theory prediction from solving Eq. (S167), compared to values of *v*_⊥_ measured from simulations. Simulations used *s* = 0.02, *N* = 10^20^, and *U*_*g*_ = *U*_*s*_ = 3 · 10^−9^ unless otherwise noted.

Despite the complexities of the frequent-specialists regime, we see that the key qualitative conclusion from the rare-specialists regime holds. Mutations with environmental tradeoffs have a significant fixation advantage over generalist mutations when they arise near the beginning of their favored environment, because transient fitness benefits have more time to compound if they occur soon after a lineage is founded. Because of this advantage, successful lineages will generically “pre-adapt,” acquiring a suite of opposing specialist mutations at times aligned with upcoming environmental states.

### Environmental stochasticity with multiple specializing lineages

When the environmental state varies stochastically (*σ*_*τ*_ > 0) in a population with common specialist mutations (*NU*_*s*_ ≫ 1), the entire population will respond to stochastic environmental change. To roughly describe this adaptation, we will continue to use the linear ansatz in Eq. (15) described by the rate of adaptation *v*_∥_ and the rate of specialization *v*_⊥_. However, the environmental state *I*(*t*) is now a random variable that does not perfectly balance the time spent in each environment, so the population will have more time to specialize toward an environment that lasts longer.

#### Optimal mutational path

The optimal mutational path will also depend on the realized environmental stochasticity – but unlike the bulk population, it will “anticipate” the environmental state that happens to be overrepresented in the near future. To formalize this, we take the fast-cycling *τ* ≪ *s/v*_∥_ limit, where the relevant unit of environmental stochasticity is the environmental bias over a single founding step of time *s/v*_∥_ ≫ *τ*:

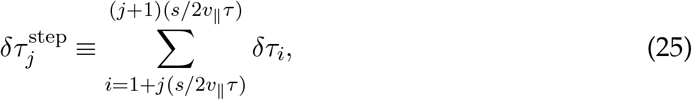

with *δτ*_*i*_ the time difference between adjacent environmental states. Solving the rolling fitness relations (SI 3.3) reveals that the optimal mutational path is given by a weighted sum of future environmental stochasticity:

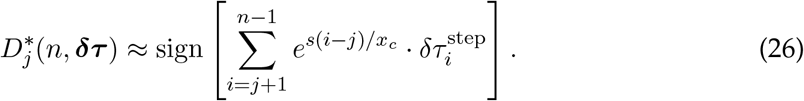

A lineage which follows this optimal mutational path will acquire an effective fitness boost of 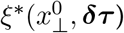, playing a similar role to *ζ*^∗^ in Eq. (19). A focal lineage will be likely to fix if its starting fitness satisfies 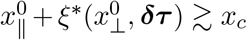. However, the fitness shift *ξ*^∗^ has a complicated dependence on the environmental stochasticity ***δτ***, since it encodes the competition between an optimally-specializing lineage and the background population. Thus, in order to average over environmental stochasticity and estimate the overall fixation probability, we employ a series of rough semianalytic approximations to the distribution of *ξ*^∗^ rather than solving it exactly (SI 3.3.1).

#### Specialist advantage

When specialist mutations are widely available, the specialist advantage 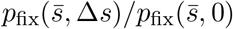 in stochastically fluctuating environments continues to be much greater than unity (Fig. 4C). This advantage arises from the contribution of a *single* specialist mutation to a lineage following the optimal mutational path, effectively shifting 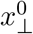 to 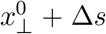. This shift widens the distribution of *ξ*^∗^ toward higher fitness values (Fig. S8), making it easier for mutations to reach fixation.

However, the specialist advantage does not attain the extreme values of ~10^6^ and larger we observed in the rare-specialists regime (Fig. 3B-C; Fig. S7B). This is because the fates of specialist and generalist mutations alike are both sensitive to environmental stochasticity, due to linkage and competition with other specialist mutations. When the entire population is able to specialize at rate *v*_⊥_, a single lineage will not be able to take sole advantage of a long stretch of beneficial environments: rather, it will compete with many other specialist mutations on other genetic backgrounds that are also rising in frequency, cutting off the benefit stochasticity can provide to the focal lineage.

#### Rate of adaptation

Because successful lineages will acquire mutations that match future environmental stochasticity, they will rise in frequency more quickly and raise the overall rate of adaptation. We can estimate this by determining how typical values of *ξ*^∗^ feed back on the rate of adaptation through Eq. (S167), finding

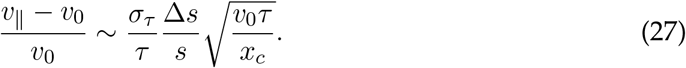

While this adjustment is nominally small, for large enough Δ*s* and *σ*_*τ*_ it can raise the population rate of adaptation by a factor of two (Fig. 5B). As in the case without seasonal drift, Eq. (27) assumes that specialist and generalist mutations provide the same average benefit *s*. If specialists have a smaller average fitness effect than generalists, this boost will decline and can become negative, when natural selection favors specialist mutations despite the fact that they contribute less to long-term fitness gain. Thus, the presence of specialist mutations can speed up or slow down the rate of adaptation depending on the shape of the underlying JDFE (Fig. S9).

#### Rate of specialization

Finally, we can estimate the rate of specialization *v*_⊥_ by tracking how the optimal mutational path depends on the realized stochasticity (SI 3.3.3):

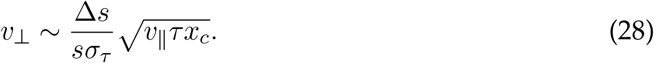

We see that *v*_⊥_ declines with the stochasticity *σ*_*τ*_, mirroring Eq. (22), which shows that *v*_⊥_ declines with *τ* in deterministic environments. Intuitively, environmental stochasticity reduces the diversity in the population by creating long stretches where one environment is favored over another, lowering the rate of specialization (Fig. S10). Thus, the rate of specialization is maximized for perfectly periodic environments with small *τ* – the same conditions where tradeoffs become effectively neutral.

Our results in this section continue to roughly apply in slower-cycling environments with *τ* ≫ *s/v*_∥_ (SI 3.3.2). In principle, for larger *τ* the optimal mutational path will depend on a combination of stochasticity and the perfectly-periodic effects discussed in the previous section. However, we find that focusing on stochasticity alone is reasonable if *σ*_*τ*_ is sufficiently large.

## Discussion

Existing theory struggles to predict how natural selection contends with varying environmental conditions, when beneficial mutations with different fitness tradeoffs compete. To address this issue, we modeled a population in a two-state environment whose members acquire linked mutations, according to a simple joint distribution of fitness effects (JDFE). We found that competition between linked mutations systematically selects for environmental tradeoffs (Fig. 6), providing environment-sensitive mutations a fixation advantage over environment-insensitive ones with larger geometric mean fitness effects – in contrast to past work where mutations accumulate one-at-a-time (16–21). This “specialist advantage” is present because a mutant lineage which arises in a favorable environment can found further beneficial mutations sooner, providing a long-term advantage even after the environment changes. The specialist advantage is most evident in the strong-tradeoff regime when the environmental fluctuation timescale is long (Δ*s* · *τ* ≫ 1) or sufficiently stochastic 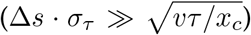, and applies whether or not environment-sensitive mutations are rare or common in the JDFE.

**Figure 6.**
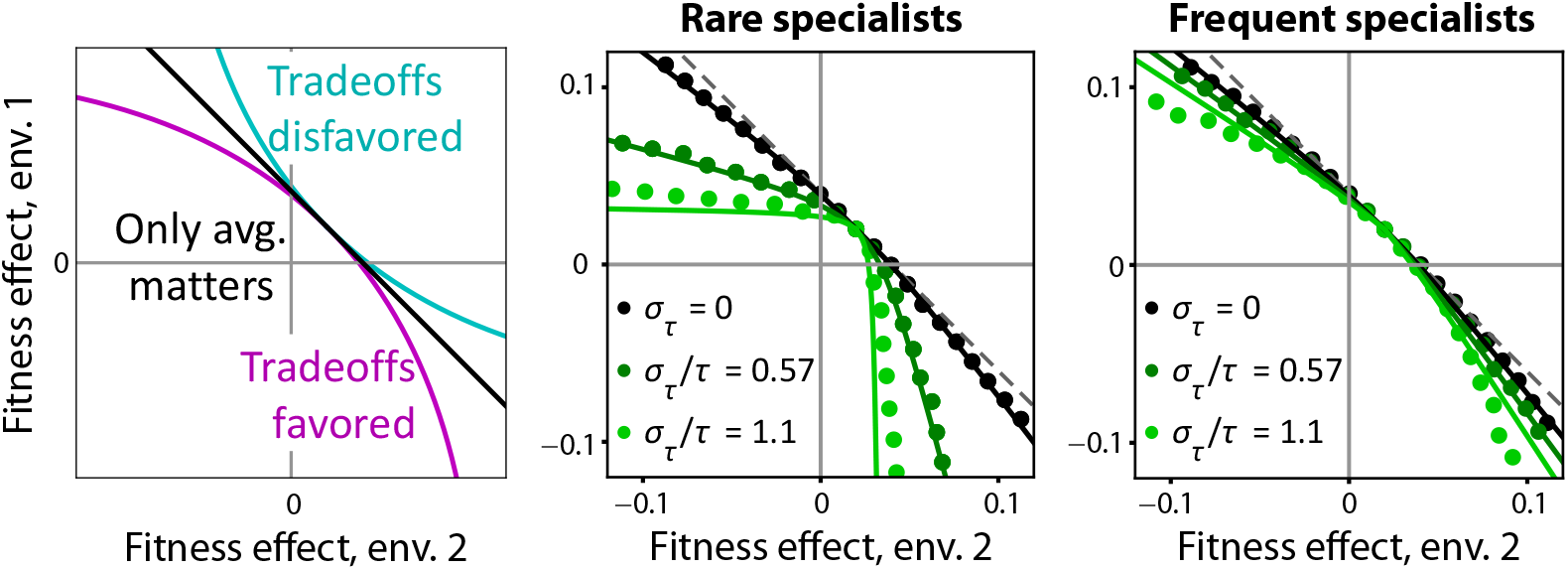
Natural selection favors environmental tradeoffs when linked mutations compete. Fixation probability isoclines (as described in SI 4.3) in two-dimensional fitness space, with fixation probability equal to *p*_fix_(*s*, 0). Plots show a schematic (left) and results for simulations with *τ* = 200 with rare and frequent specialist mutations (center and right). Dashed lines show the “average only” curve in the schematic, equivalent to *τ* 0 and *σ*_*τ*_ */τ* → 0. Solid lines show theory predictions. Parameters are the same as Fig. 4B unless otherwise noted.

Our results demonstrate how the fitness effects and tradeoffs of *successful* mutations depend on environmental fluctuations and the underlying JDFE, quantities which can be measured or manipulated in evolution experiments (9, 22, 26). Renewable barcoding systems are particularly potent experimental tools to study rapidly adapting populations, because they allow beneficial mutations to be detected while they are still at low frequencies (14, 26). Applying these methods to fluctuating environments would test our prediction that as the fluctuation timescale increases (for a fixed JDFE), natural selection will increasingly filter for strong-tradeoff mutations that arise near the start of favorable environments.

When environment-sensitive mutations are common, clonal interference between mutations with different tradeoff profiles dramatically reshapes the outcome of natural selection. By tracking how the fate of a lineage depends on the joint fitness effects of the entire sequence of mutations it acquires, we found that successful lineages generically pre-adapt to upcoming environmental transitions, by acquiring well-timed mutations with strong environmental tradeoffs. This alignment between tradeoffs and environmental change allows beneficial mutations to rise in frequency more quickly, speeding up the rate of adaptation of the population as a whole. However, environment-sensitive mutations can also slow down adaptation, by outcompeting constitutively beneficial mutations with higher geometric mean fitness. Thus, natural selection does not always optimize long-term fitness gain, and the rate of adaptation is sensitive to the shape of the two-dimensional JDFE.

In this work, we limited our analysis to a two-state environment, allowing us to focus on a single axis of environmental tradeoffs. While this approach could be considered a rough model of seasonal cycles (44) or experimental evolution protocols (9, 11), natural populations can experience a much larger number of potential environmental states. We conjecture that many of our conclusions should naturally extend to a broader set of environments: the fate of a mutation is determined by its fitness across each environment it encounters on its way to fixation, with more weight placed on early environments. On the population level, rapid adaptation constitutes a strategy to respond to environmental change, comparable to bet-hedging by stochastically switching between phenotypes (45). This evolutionary strategy is likely to be favored when environments change over many generations (46), relative to strategies like genetic regulation (47, 48) which allow for phenotypic flexibility on shorter timescales, but may carry fitness costs (49, 50).

Our work focuses on a simplified point JDFE that neglects epistatic interactions between mutations. In reality, the fitness effects of mutations depend not only on the environmental state but also on the mutation’s genetic background. While these epistatic fitness landscapes are beyond the scope of this work, recent work has begun to characterize their environmental dependence empirically (10). The structure of these landscapes critically determines whether long-term evolution tends to alleviate the effects of tradeoffs (51, 52), or sharpen them in the form of a Pareto front at the “boundary” of mutational space, where any increase in fitness requires a corresponding cost in another environment (53). Our results demonstrate that successful mutations can still carry strong tradeoffs even when unconditionally beneficial mutations are available, far from a hypothetical Pareto front. This circumstance is plausible for an organism which has recently migrated to a new environment, or for experimentally measured fitness effects in non-natural environments (9, 11).

Finally, a key limitation of this work was ignoring recombination between genomes, which has the potential to break up the linkage between alleles that underlies many of our results. Recombination has been proposed to be an important mechanism of adaptation in fluctuating environments, providing a gene pool of environment-specific alleles that allow for modularity under changing environmental conditions (54, 55). Understanding how these factors conspire to shape long-term evolutionary outcomes in fluctuating environments is a key direction for future work.

## Data and Code Availability

Code for simulations and figure generation is available on GitHub (github.com/jdmcenany/ Fluctuating_Environments). Repository of precomputed simulation results is available on Zenodo (56).

## Acknowledgments

We thank Daniel Wong and members of the Good lab for useful discussions and feedback. This work was supported in part by NIH NIGMS Grant No. R35GM146949. B.H.G. is a Biohub – San Francisco Investigator.

## Supplementary Information

**Figure S1.**
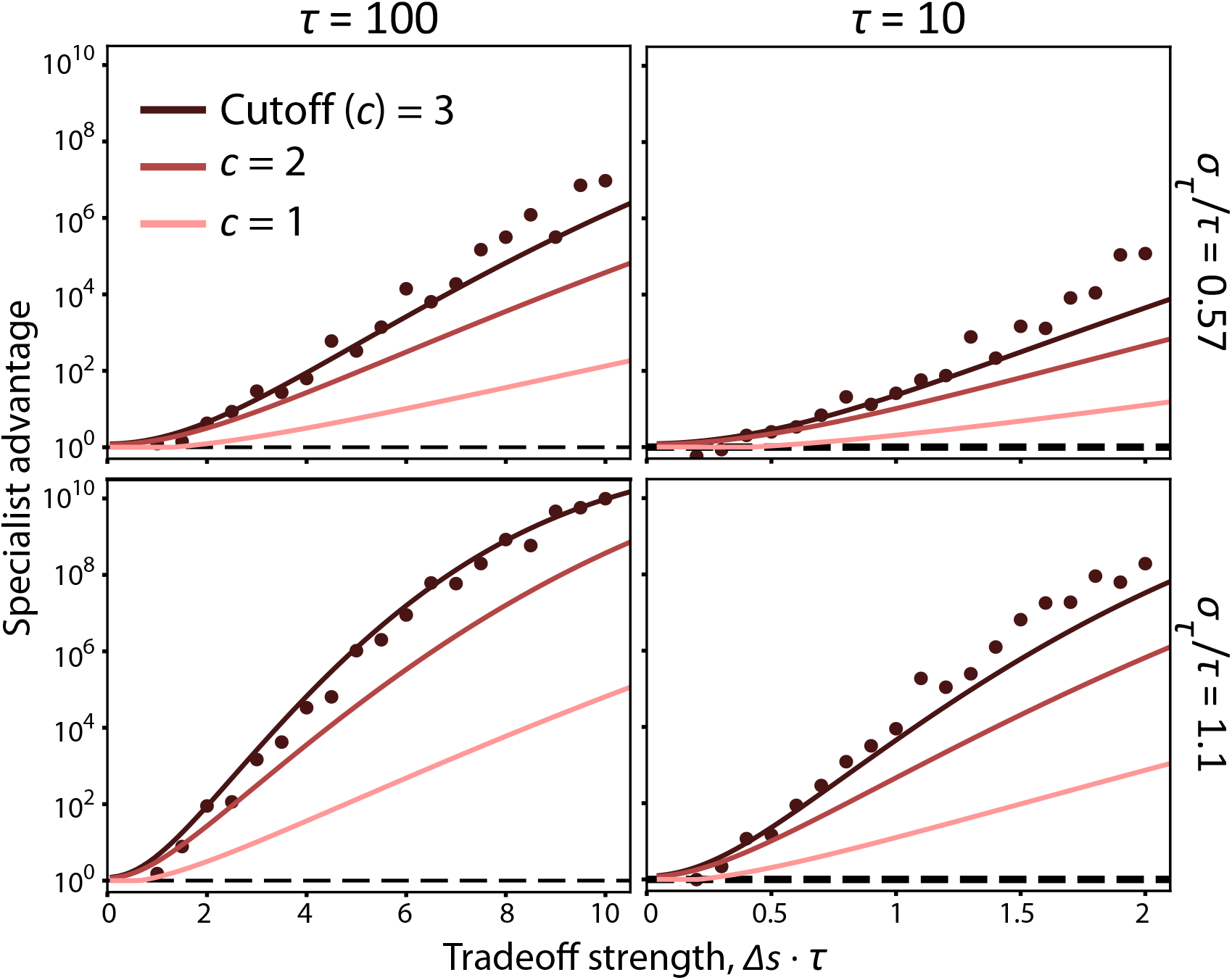
Typical stochastic fluctuations still provide a large advantage to specialists. Points show the specialist advantage 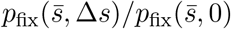 in a stochastic environment, for different values of *τ*, *σ*_*τ*_ and Δ*s*. Curves show theory prediction as in Fig. 3B-C for different values of *c*, the cutoff (measured in standard deviations) imposed on stochastic fluctuations. Typical environmental fluctuations with *c* ~ 1 still have a large specialist advantage, although reduced from the full average captured over many independent simulations. Simulations were run with *N* = 10^20^, 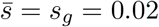, and *U*_*g*_ = 3 · 10^−9^.

**Figure S2.**
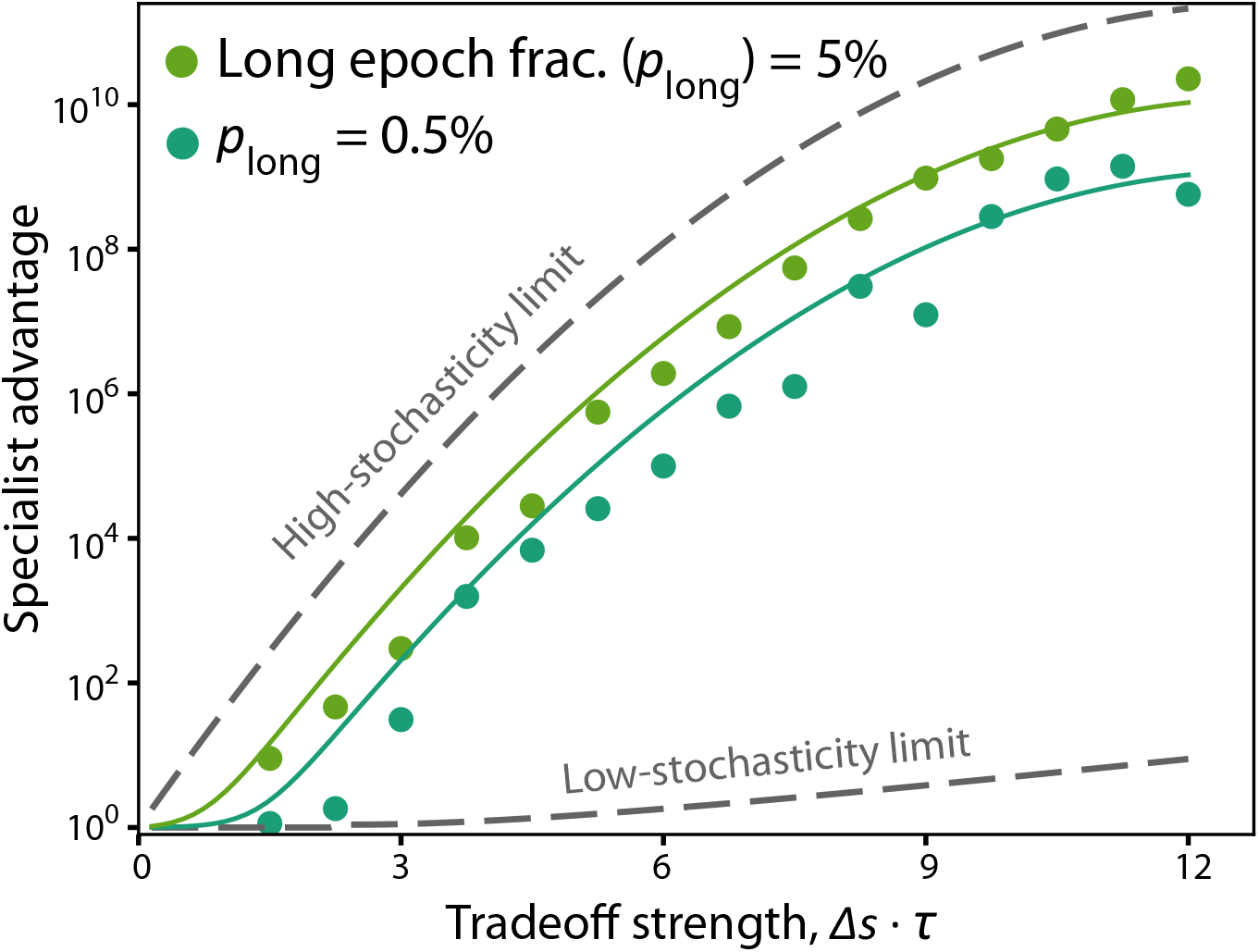
Rare environment-sensitive mutations in high-variance environments. Specialist advantage 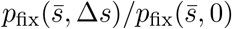 in a high-variance environment where environmental states usually last *τ* = 150 generations, but rarely last *τ*_long_ = 1000 generations with probability *p*_long_. Points show simulation results for two values of *p*_long_. Bottom dashed line shows the low-stochasticity limit (*σ*_*τ*_ = 0) from Eq. (S31) and Eq. (S32). Top dashed line shows the high-stochasticity limit, 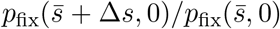. Theory curves show the low-stochasticity limit plus *p*_long_ times the high-stochasticity limit, which is dominated by the probability of a rare long epoch. Simulations were run with 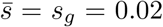, and *U*_*g*_ = 3 · 10^−9^ with rare specialist mutations.

**Figure S3.**
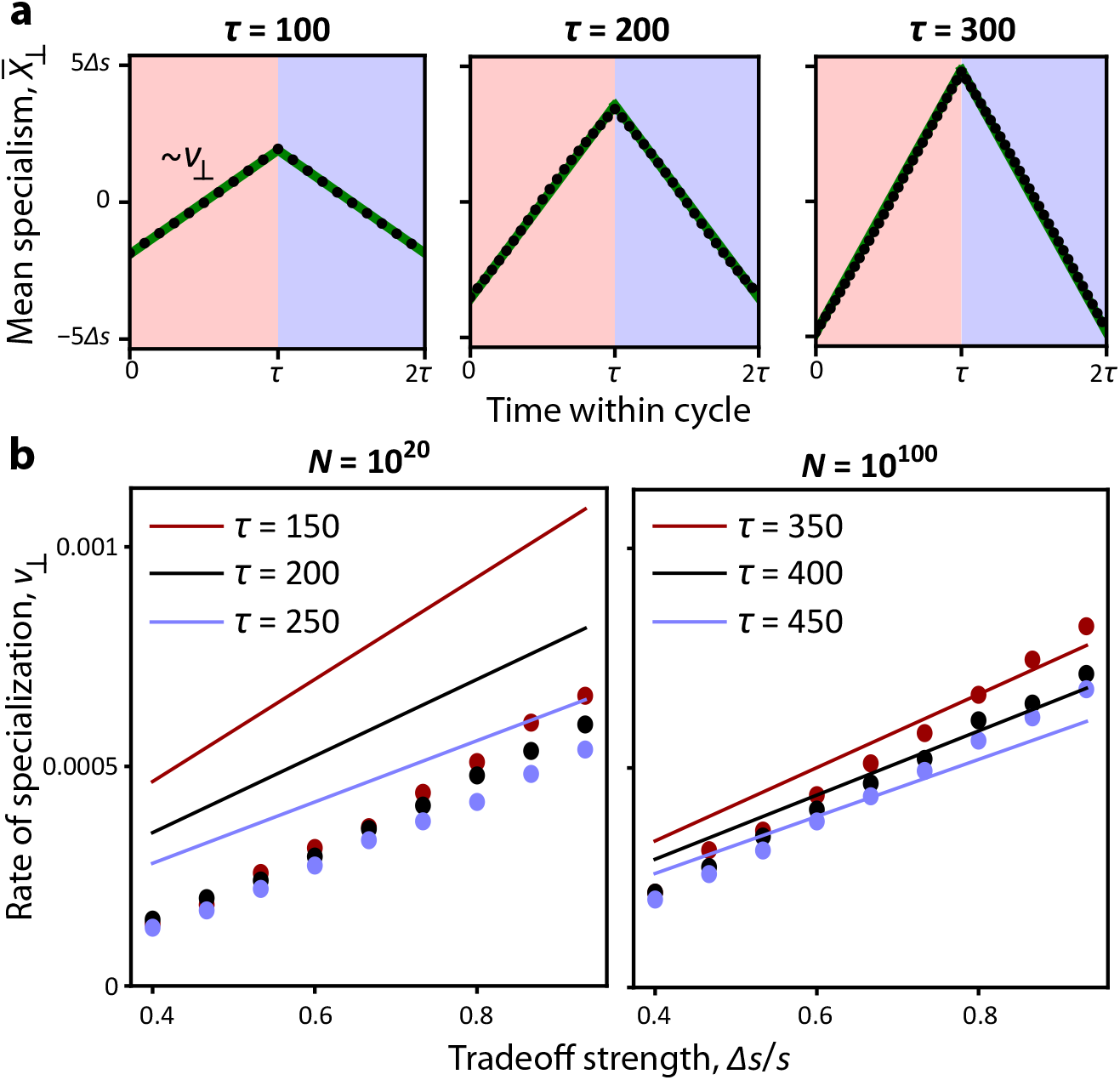
The bulk population specializes toward the current environment. **(a)** The mean specialism of the population 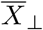 as a function of time within an environmental cycle, for different environmental cycling times *τ*. Points show simulation averages (measured relative to the start of an environmental cycle and normalized to have mean zero), while green curves are fit lines. The slope of these fit lines defines *v*_⊥_. **(b)** Rate of specialization *v*_⊥_ as a function of tradeoff strength Δ*s* and environmental cycling time *τ*. Points represent simulation results, while theory curves are from Eq. (S71) (using the leading-order approximations for *v*_∥_ and *x*_*c*_ from the rare-specialists regime). Note that the theory only agrees quantitatively with simulations when the population size is large enough to strongly satisfy all the regime assumptions. Parameters for (a) and (b), left: *N* = 10^20^, *s* = 0.015, *U*_*g*_ = *U*_*s*_ = 10^−6^. Parameters for (b), right: *N* = 10^100^, *s* = 0.015, *U*_*g*_ = *U*_*s*_ = 10^−15^.

**Figure S4.**
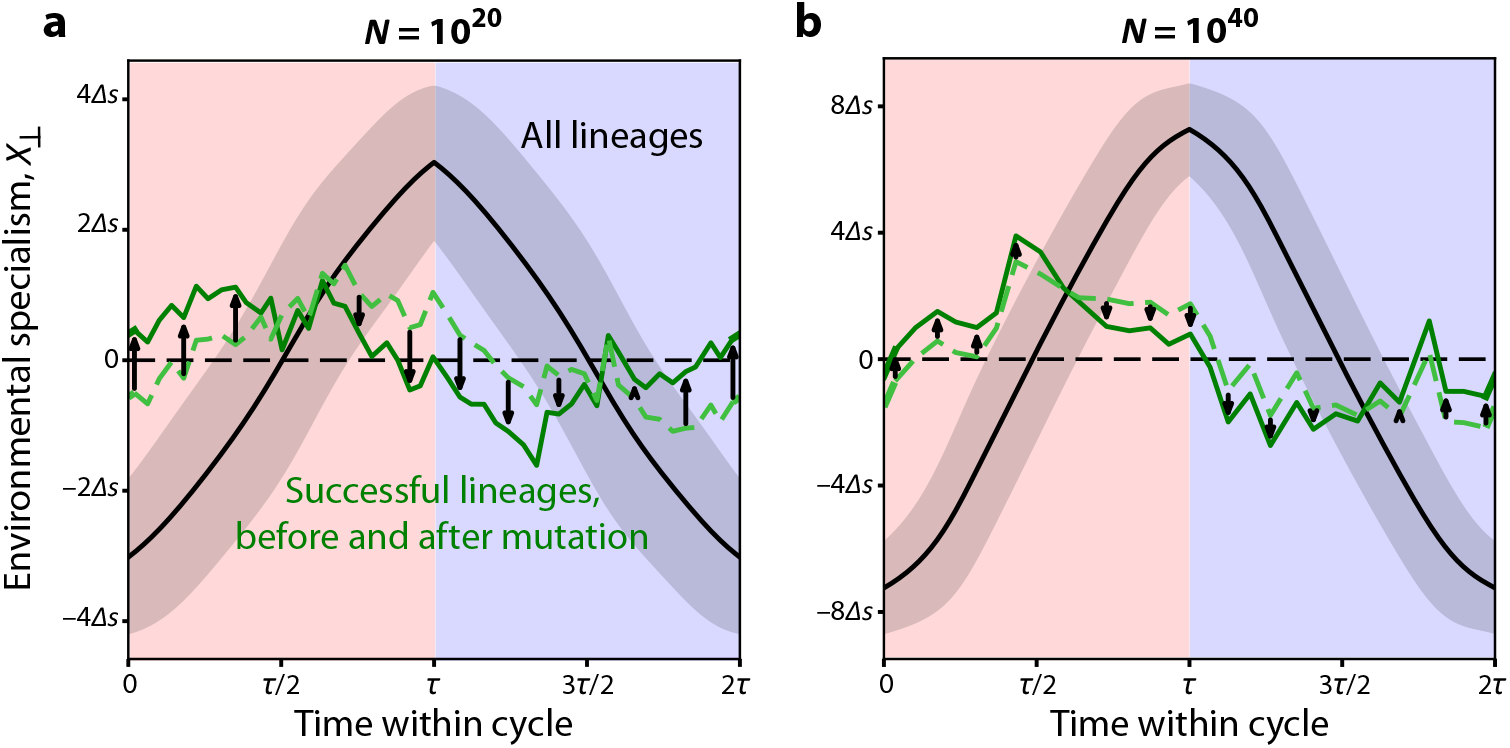
Optimal mutational path in asymptotically large populations. **(a)** The empirical specialism *X*_⊥_ of lineages over a cycle, with the same simulation results as in Fig. 4B. Black points show the population mean, with shading ± one standard deviation. Green curves show the specialism of successful mutants (i.e., those fated for fixation) immediately before (dashed) and after (solid) a new mutation, with the arrows indicating the direction of mutation. Mutations are binned by establishment time. Simulations were run with *N* = 10^20^, *U*_*g*_ = *U*_*s*_ = 3 · 10^−9^, *s* = 0.02, Δ*s* = 0.05, and *τ* = 200. **(b)** Similar plot as (a), but in a larger population with *N* = 10^40^, *U*_*g*_ = *U*_*s*_ = 5 · 10^−9^, and *s* = 0.03. Note the *y*-axis spans a wider range of specialism.

**Figure S5.**
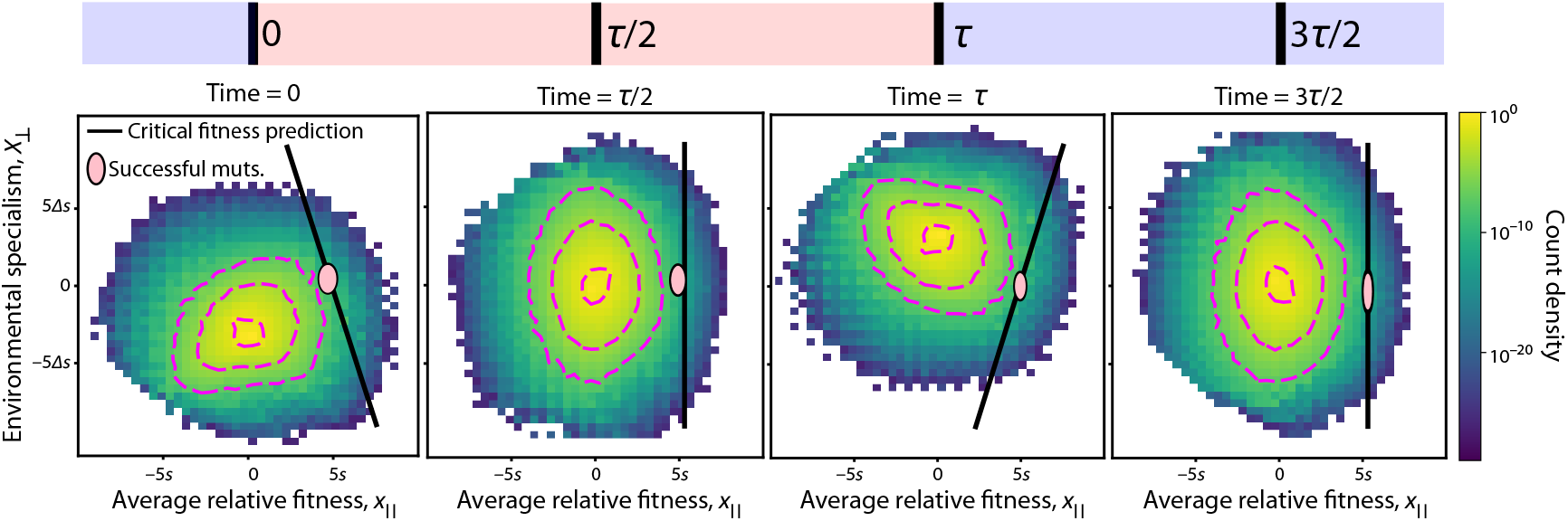
Two-dimensional fitness distribution in a population with frequent specialist mutations. Two-dimensional fitness distribution of the population at various points in the environmental cycle, parametrized by average fitness *X*_∥_ and environmental preference *X*_⊥_. Dashed magenta curves show empirical contour lines for densities 10^−1^, 10^−4^ and 10^−7^. Solid black line is the theory curve where *x*_∥_ + *s* + *ζ*^∗^(*x*_⊥_ + *δs, t*_0_) = *x*_*c*_, with *ζ* defined in Eq. (S68); fixation is likely for mutatnts which arise on or to the right of this curve. Pink ellipse shows the average values of *x*_∥_ and *X*_⊥_ for lineages which have just acquired mutations fated for fixation, with the height and width of the ellipse equal to twice the standard deviation among these successful lineages in each dimension. Note that the pink ellipse is expected to roughly correspond with the maximum-density portion of the black line. An animated GIF of this distribution over the full time range can be found on Zenodo (56), where the position of the ellipse was inferred via linear interpolation. Simulation was run with *N* = 10^20^, *U*_*g*_ = *U*_*s*_ = 3 · 10^−9^, *s* = 0.02, Δ*s* = 0.05, and *τ* = 200.

**Figure S6.**
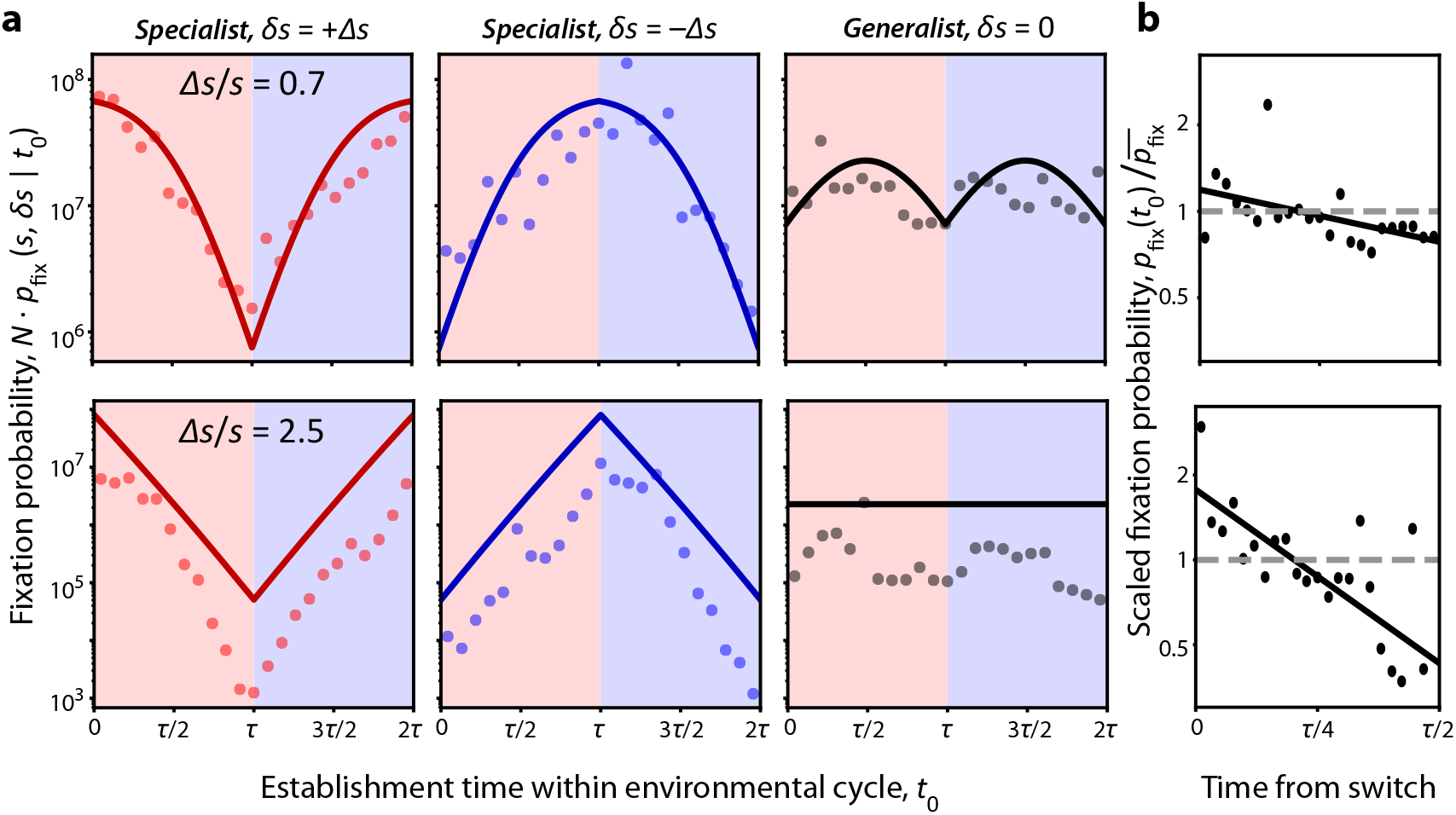
Time-dependent fixation probabilities when specialist mutations are frequent. **(a)** Fixation probabilities of each type of mutation as a function of their establishment time in a cycle, in a population where specialist mutations are common. Solid lines show theory prediction from Eq. (S102) (top row) or Eq. (S134) (bottom row). **(b)** Fixation probability averaged across all types of mutations, as a function of the time to or from the nearest environmental switch in the cycle. Dashed line shows the time-averaged fixation probability, while solid line is a best-fit line in log space, demonstrating that successful mutations tend to occur near environmental switches when tradeoffs are large. Top row: parameters are the same as Fig. 4B, with Δ*s > s*. Bottom row: *N* = 10^20^, *U*_*s*_ = *U*_*g*_ = 10^−10^, *s* = 0.022, Δ*s* = 0.015, and *τ* = 300. Note that *s >* Δ*s*, meaning tradeoffs are weaker than the generalist benefit of mutations.

**Figure S7.**
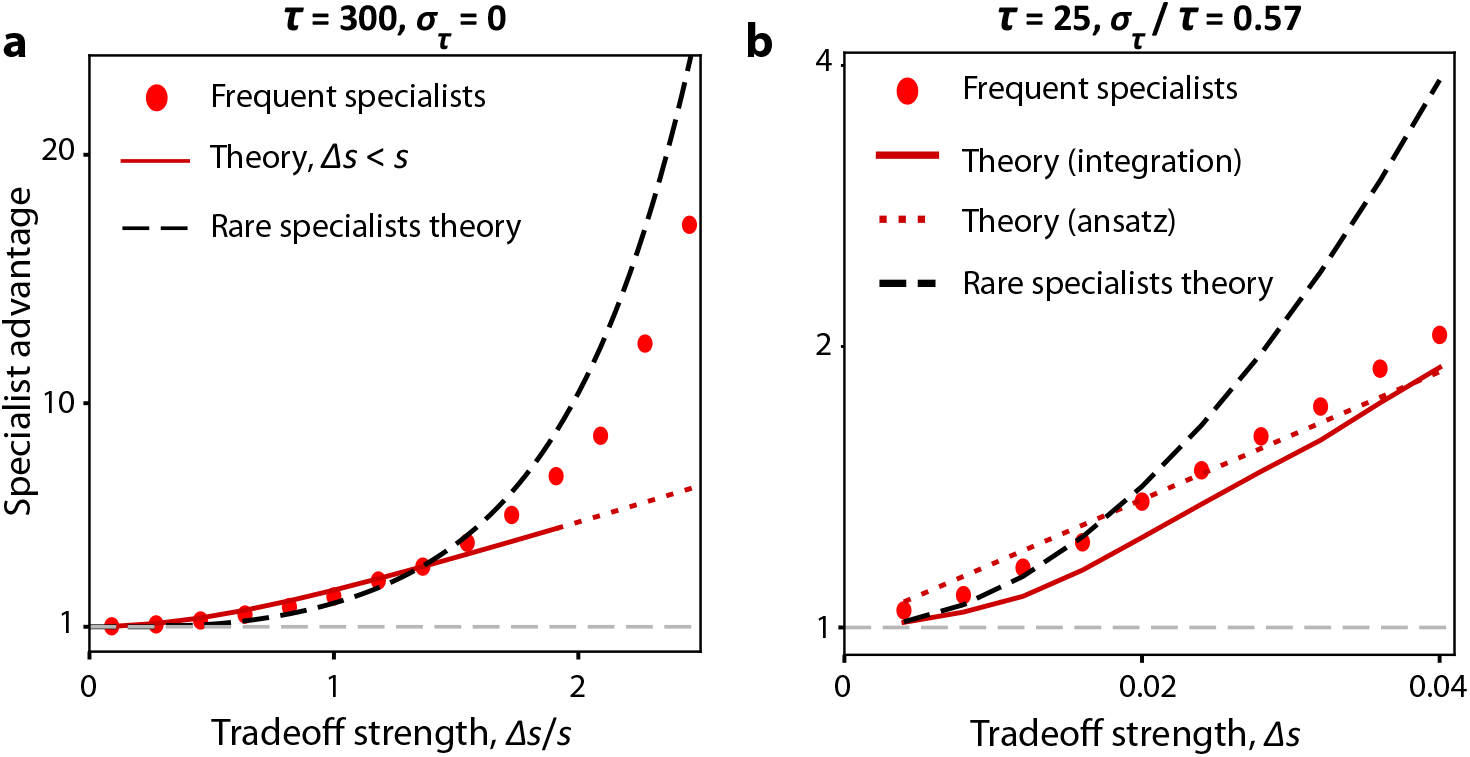
The specialist advantage depends on the mutation rate of specialist mutations. **(a)** The specialist advantage *p*_fix_(*s*, Δ*s*)*/p*_fix_(*s*, 0) in a periodic environment where specialist mutations are common. Points show simulation results, while red curve shows prediction from averaging Eq. (S102) over time, valid when Δ*s < s*. Black dashed line shows rare-specialists prediction from Eq. (S31). Note that the specialist advantage increases to become exponential when *s >* Δ*s*, similar to the rare-specialists regime. Parameters: *N* = 10^20^, *s* = 0.022, *U*_*g*_ = *U*_*s*_ = 10^−10^ (or *U*_*g*_ = 2 · 10^−10^ and *U*_*s*_ → 0 for the rare-specialists curve). **(b)** The same plot for a stochastically fluctuating environment. Red curve shows prediction from numerically integrating Eq. (S156) for fit *σ*_eff_; dotted red curve shows ansatz from Eq. (S165). Black curve shows rare-specialists prediction from Eq. (S46). Parameters: *N* = 10^20^, *s* = 0.02, *U*_*g*_ = *U*_*s*_ = 3 · 10^−9^ (or *U*_*g*_ = 6 · 10^−9^ and *U*_*s*_ → 0 for the rare-specialists curve).

**Figure S8.**
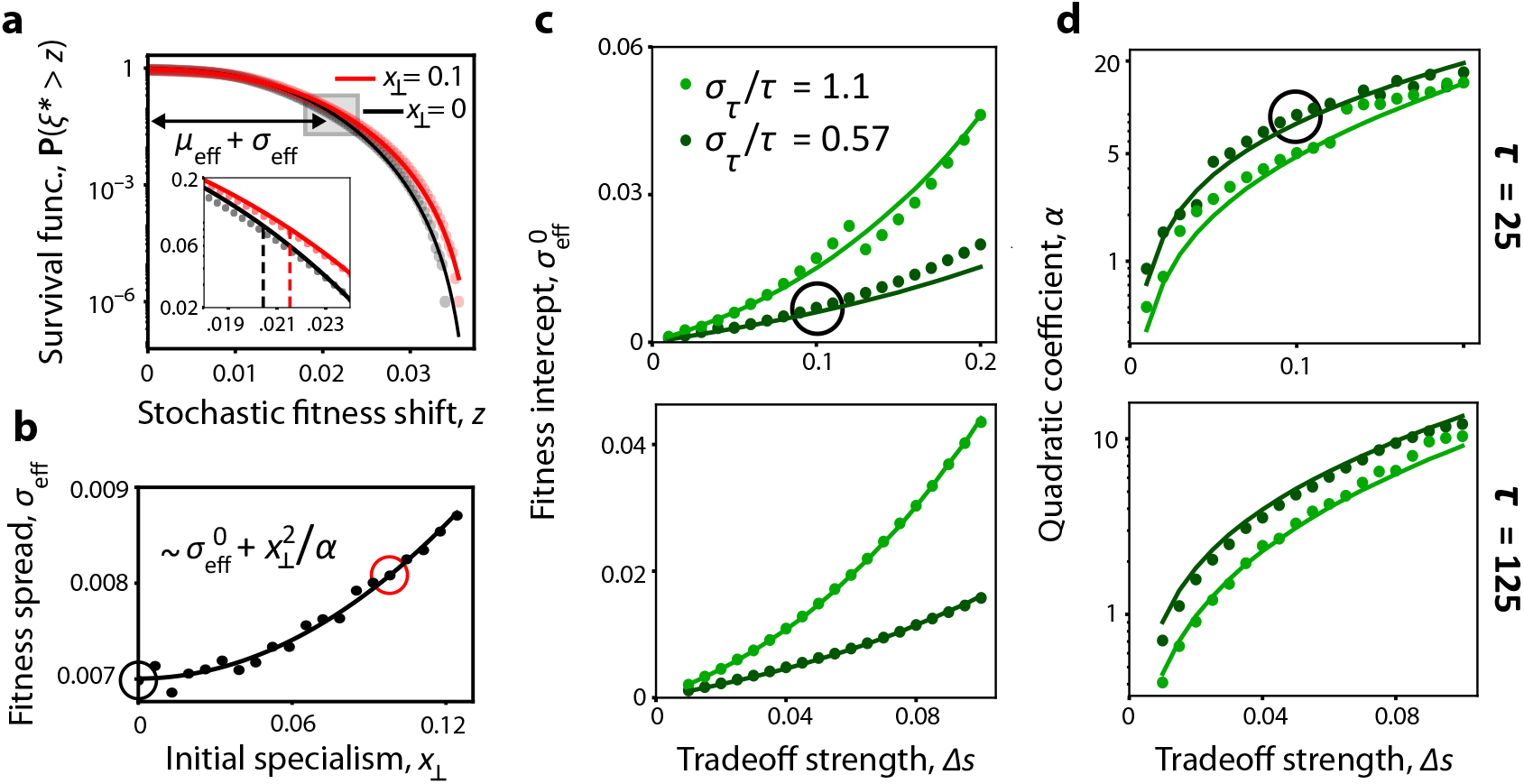
Characterizing the effect of seasonal drift on fixation in specializing populations. **(a)** Positive tail of the empirical survival function of the stochastic fitness shift *ξ*^∗^, defined in Eq. (S143), for two values of initial specialism *x*_⊥_. Curves show the best fit of *σ*_eff_ to Eq. (S149). Inset shows a zoomed-in version of the shaded region with dashed lines indicating *µ*_eff_ + *σ*_eff_ for each distribution. **(b)** Dependence of fit *σ*_eff_ on *x*_⊥_ was parametrized using a quadratic fit for *x*_⊥_ between 0 and 2Δ*s*, yielding intercept 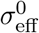 and quadratic scale *α*. Circles correspond to red and black curves in (a). **(c)** Dependence of the intercept *σ*^0^ on Δ*s* and *σ*_*τ*_ for small *τ* (top) and large *τ* (bottom). Curves show the ansatz in Eq. (S153) with fit *C*_0_ = 0.10 (top) or 0.086 (bottom). Circled point corresponds to the parameters in (a). **(d)** Dependence of the quadratic scale *α* on Δ*s* and *σ*_*τ*_ for small *τ* (top) and large *τ* (bottom). Points show fits as in (b), while curves the ansatz in Eq. (S154) with fit *C*_*α*_ = 1.2 or 3.5 (bottom). Parameters in all panels are the same as Fig. 4C, with *v*_∥_ and *v*_⊥_ measured from simulations and *x*_*c*_ calculated as a function of *v*_∥_.

**Figure S9.**
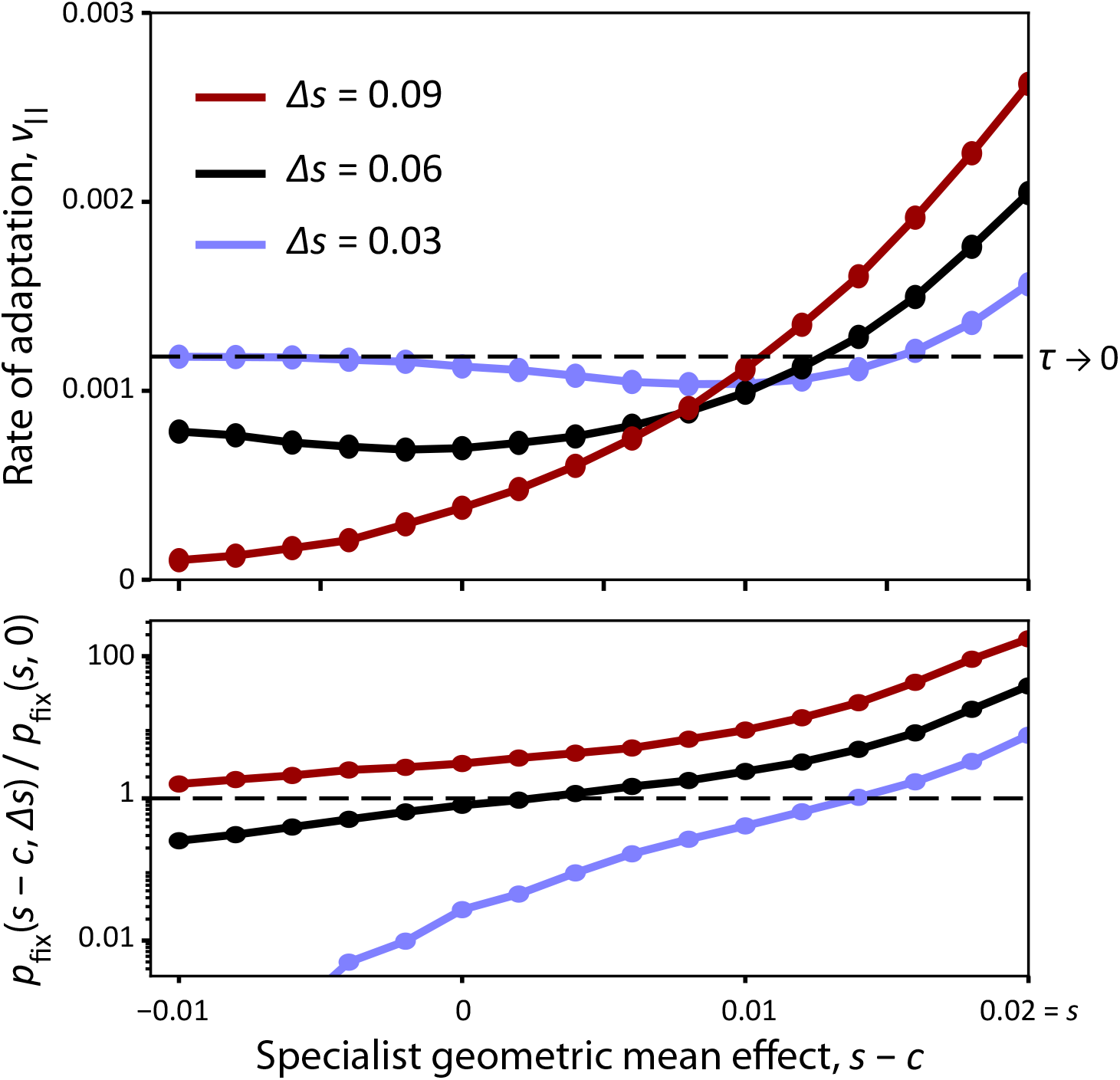
The rate of adaptation depends on the joint distribution of fitness effects. Top: Long-term rate of adaptation *v*_∥_ in a stochastically varying environment measured in simulations where generalist mutations have fitness effect (*s*, Δ*s*) and specialist mutations have fitness effect (*s* − *c*, Δ*s*) for different fitness costs *c* and tradeoff strengths Δ*s*. Dashed line shows the no-specialists rate of adaptation (*τ* → 0 or *U*_*g*_ = 0). Bottom: The specialist advantage for the same parameters as the top panel. The dashed line shows where fixation probabilities of specialist and generalist mutations are equal. Simulation parameters were *N* = 10^20^, *U*_*g*_ = *U*_*s*_ = 3 · 10^−9^, *s* = 0.02, *τ* = 125, and *σ*_*τ*_ */τ* = 1.1.

**Figure S10.**
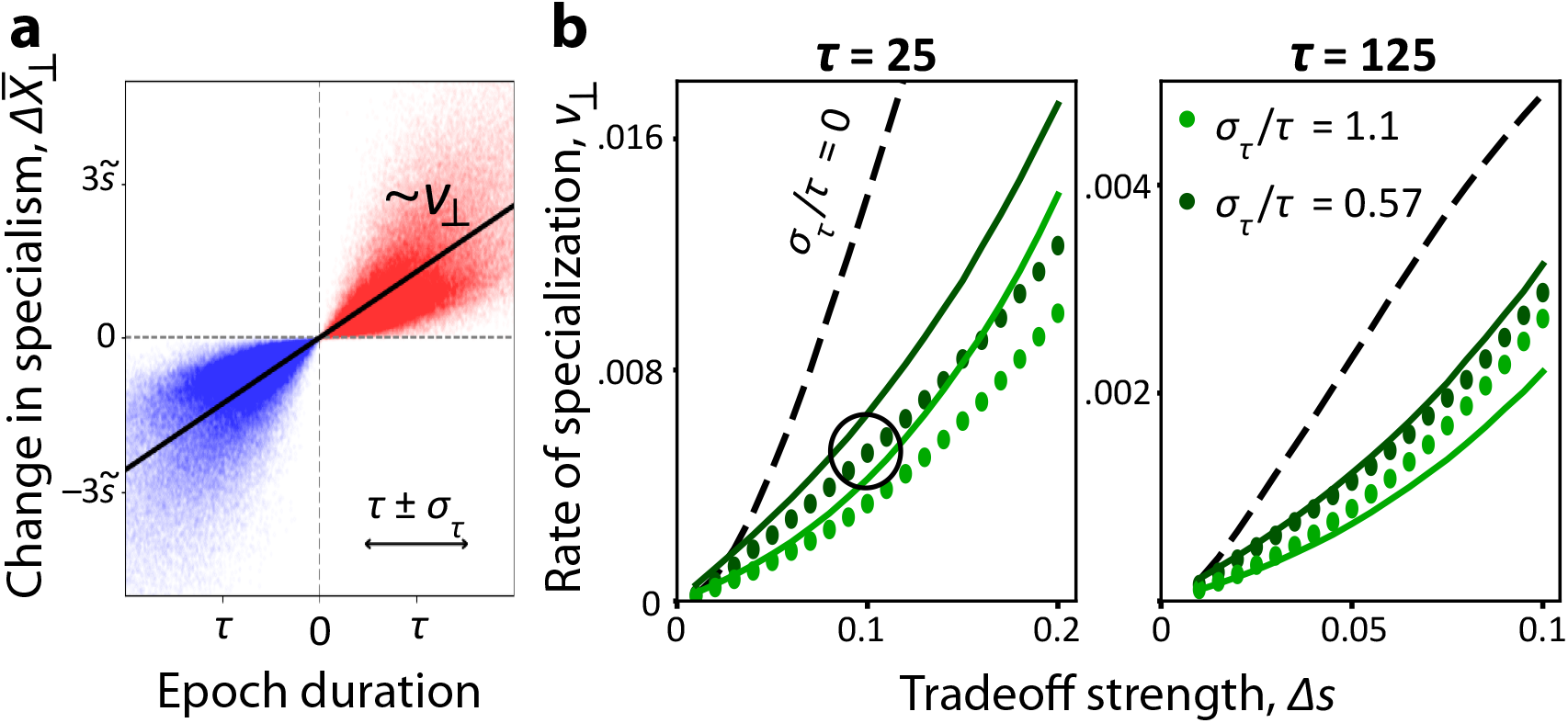
The rate of specialization declines in stochastic environments. **(a)** Scatterplot of the change in population mean specialism 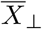 over a single environmental epoch, as a function of the stochastic epoch length (with different environmental states reflected across zero). The slope of the best-fit line to this plot is defined as the rate of specialization *v*_⊥_. **(b)** Measured rate of specialization *v*_⊥_ for stochastic environments (points) and periodic environments (dashed line), compared to theory prediction in Eq. (S178), left, or Eq. (S179), right. Parameters in all panels were the same as Fig. 4C, with (a) corresponding to the circled point in (b).

## 1 Model Overview

### 1.1 Clonal Interference Regime

This work follows a body of literature modeling the evolution of populations in the “clonal interference” regime (38–41), where each beneficial mutation must compete with many others in order to reach fixation. Thus, much of our model setup follows previous work, whose central assumptions we restate here for completeness.

We consider a single-species population of constant size *N* ≫ 1, whose members evolve over time by acquiring beneficial mutations. For simplicity, we model these mutations with a coarse-grained point distribution of fitness effects in the infinite-sites limit: beneficial mutations arise at a constant rate *U* (regardless of background), and each mutation has the same incremental log fitness effect *s*. Under these assumptions, any individuals with the same number of beneficial mutations will have the same fitness, making it convenient to define individuals by their “fitness class.” Over time, new higher-fitness classes will be founded as old ones die out, and the absolute fitness of the population will increase.

Generally speaking, the empirically relevant clonal interference regime will apply for a broad range of parameters when *Ns* ≫ *NU* ≫ 1, such that many beneficial mutations arise each generation. More precisely, the parameter range we consider satisfies

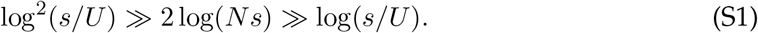

In physical terms, the left inequality requires that the fitness benefits of individual mutations are strong enough to be felt in the time it takes for a new fitness class to be founded. The right inequality states that mutations are frequent enough that at any given time, individuals with a range of fitnesses coexist, rather than a “pseudo-sweep” regime where only two fitness classes are present at once.

When these conditions are satisfied, the steady-state fitness distribution of the population can be described by a “traveling wave” with well-studied properties. This wave advances at rate *v*, the rate of adaptation of the population. While most individuals in the population have fitness close to the mean, the high-fitness “nose” of the wave has a fitness advantage *x*_*c*_ relative to the bulk. It is often convenient to work with these derived parameters, whose dependence on *N* and *U* has been calculated in previous work:

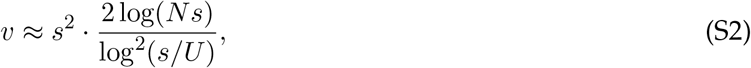

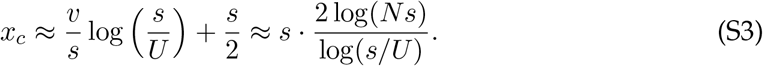

Expressed in these parameters, the clonal interference conditions are *s*^2^ ≫ *v* and *x*_*c*_ ≫ *s*. The distribution of relative fitness of the population can be approximated by

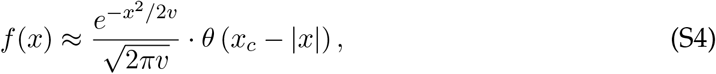

where the Heaviside function *θ*(*x*) sets a fitness cutoff.

### 1.2 Mutations in a Fluctuating Environment

We would now like to extend the framework in the previous section to account for mutations whose effects depend on a time-varying environment. We can account for this in the simplest way possible by considering only two environments, *E*_1_ and *E*_2_. Suppose that the environmental state is given by a time-dependent indicator function *I*(*t*) = ±1, which applies to all members of the population. To begin with, we consider an environment where each environmental state *E*_1,2_ lasts exactly *τ*_1,2_ generations:

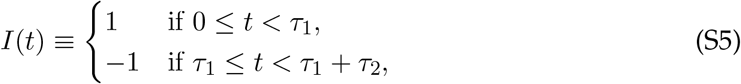

and is defined by the periodic relationship *I*(*t* + *τ*_1_ + *τ*_2_) = *I*(*t*) afterward. Through most of the main text, we focus on the symmetric case *τ* ≡ *τ*_1_ = *τ*_2_ (although see SI 2.1.5 for the more general case). In this symmetric case, it is also useful to define the function

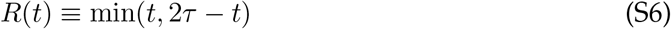

for 0 ≤ *t <* 2*τ*, and periodic with period 2*τ* otherwise. This triangle wave has the useful interpretation that *R*(*t*_1_) − *R*(*t*_0_) is equal to the difference in the number of generations spent in each environmental state between *t*_0_ and *t*_1_. Later, we will extend this framework to consider the impact of seasonal drift, where the duration of each environmental epoch is a random variable.

The (log) fitness effect of a general mutation must now be expressed as a two-dimensional quantity, (*s*_1_, *s*_2_). It is convenient to reparametrize these fitness effects into geometric mean fitness 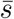 and “tradeoff strength” Δ*s*, defined as

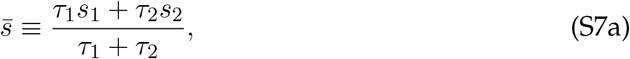

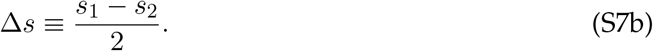

In the symmetric case *τ*_1_ = *τ*_2_, these new variables form an orthogonal basis and the original fitness effects can be simply written as 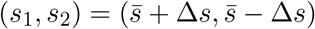.

Aggregating over the fitness effects of all possible mutations will yield a two-dimensional joint distribution of fitness effects (JDFE), which we choose to express in the 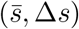 basis. In this work, we will focus on a particularly simple JDFE where mutations are grouped into three classes, which each have constant fitness effects. “Generalist mutations” have fitness effect (*s*, 0), providing the same constant fitness boost *s* in both environments (notated as *s*_*g*_ in the main text). By contrast, there are two symmetric classes of “specialist mutations” with fitness effects 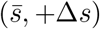 and 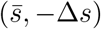. The tradeoff component Δ*s* of these mutations causes them to favor one environment over the other. We will suppose that generalist mutations arise at rate *U*_*g*_, and each type of specialist arises at rate *U*_*s*_*/*2, regardless of genetic background.

The behavior of the population will be highly sensitive to the environmental fluctuation timescale. In particular, past research suggests the existence of simple regimes where mutations in fluctuating environments behave like mutants in static ones. The first case is *τ*_1,2_ ≪ 1*/*Δ*s*, where the environment switches too quickly for the tradeoff to significantly change the frequency of a specialist mutation. In this case, selection will proceed along the direction of geometric mean fitness, so only the mean fitness effect 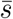 of the mutation matters. Conversely, when *τ*_1_ and *τ*_2_ are greater than the fixation time *T*_fix_, then the mutant will either reach fixation or go extinct in a single environment, so its fitness in the other environment will be irrelevant. This fixation time is known to be *T*_fix_ ≈ *x*_*c*_*/v* for a traveling wave in a constant environment. However, these limiting cases do not describe a wide range of intermediate fluctuation times,

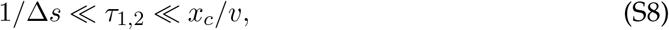

where specialist mutations with environmental tradeoff Δ*s* may behave very differently from generalist mutations with Δ*s* = 0. The rest of our analysis focuses on this intermediate regime.

We will also specialize to the regime where the tradeoff strength is not so strong that it dominates the spread in generalist fitnesses in the population, requiring

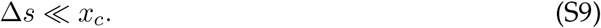

Under this condition, specialist mutations which arise on high-fitness backgrounds can have a positive initial growth rate ~*x*_*c*_ ± Δ*s* regardless of the environmental state. Importantly, this does not set a restriction on the relative sizes of 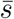 and Δ*s*, meaning that tradeoffs of individual mutations can still be strong or weak relative to their average effect.

A final important scale is the dimensionless parameter *vτ*_1,2_*/s*, which can either be much larger or smaller than unity given our other assumptions. In the fast-cycling regime defined by *vτ*_1,2_*/s* ≪ 1, a newly established lineage will not produce a new beneficial mutation until many environmental cycles have passed. In the slow-cycling regime *vτ*_1,2_*/s* ≫ 1, a lineage founded near the beginning of an environmental state is likely to have descendants with many new mutations by the time the environment switches. We will consider the effects of environmental fluctuations in both of these regimes and find that they are often surprisingly similar, although we will sometimes specialize to one regime for ease of calculation.

## 2 Fixation Probability of a Rare Specialist

To understand how specialist mutations behave when subject to clonal interference, we will first aim to calculate their fixation probability while rare. More precisely, we will assume that the specialist mutation rate *U*_*s*_ and the generalist mutation rate *U*_*g*_ satisfy *NU*_*s*_ ≪ 1 ≪ *NU*_*g*_, so that no more than one specialist lineage is typically present in the population at any given time. This assumption enables us to consider the fixation probability of a single specialist lineage competing against a purely generalist background, which is well-described by the traveling wave framework discussed above. For simplicity, we will also assume symmetry in temporal fluctuations (*τ* ≡ *τ*_1_ = *τ*_2_), relaxing this assumption in SI 2.1.5.

Crucially, the specialist lineage is able to acquire additional generalist mutations at rate *U*_*g*_ (just like the rest of the population), which increase its fitness by *s* but have no effect on its level of specialism. In biological terms, rare specialists might arise in an evolving population where a small fraction of mutations can exploit a subtly changing environmental niche, which most of the population is insensitive to. More broadly, the rare-specialists regime will provide crucial intuition for the more generic *U*_*s*_ ~ *U*_*g*_ case, making it useful to analyze first.

### 2.1 Fixation in a Deterministic Environment

We will first analyze the case of a perfectly periodic environment, which switches between states every *τ* generations according to Eq. (S5).

#### 2.1.1 Successive Establishments and Rolling Fitness

To analyze the fixation of specialist lineages, we will employ a heuristic approach, tracking the time it takes a lineage to found successively more-fit offspring. This calculation is similar to that of Ref. (42), extended to the case of fluctuating environments. The central idea is that the fate of a lineage will be determined by whether its descendants are able to increase their fitness (by acquiring additional mutations) faster than the rest of the population.

Consider a specialist mutation which arises at time *t*_0_ on an otherwise generalist genetic background. Its initial relative fitness will be *x*_0_ +*I*(*t*_0_) · Δ*s*, where *x*_0_ is the geometric mean fitness, which depends on the mutation’s original genetic background. If the specialist genotype establishes in the population (i.e., rises to high enough frequency to escape initial genetic drift), it will thereafter undergo approximately deterministic dynamics,

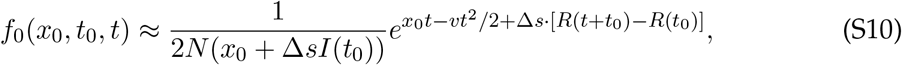

where *t* tracks the number of generations since the genotype’s establishment. We will focus on high-fitness genotypes with *x*_0_ ~ *x*_*c*_, since lower-fitness genotypes are unlikely to contribute to fixation. In this case, *x*_0_ will be the dominant fitness scale, and non-exponential correction factors such as the Δ*sI*(*t*_0_) in the denominator can be ignored.

Eventually, this genotype will begin establishing beneficial generalist mutations, founding a new fitness class with higher absolute fitness. Ref. (42) demonstrates that because the new fitness class grows more quickly than its parent genotype, it will be dominated by descendants of the *first* successful mutation, rather than future mutants that have had less time to grow at higher fitness. This result implies that the frequency trajectory of the new class can be approximated as that of a single genotype, of the same form as Eq. (S10) – but with a new mean establishment fitness *x*_1_. Accounting for environmental fluctuations also requires that we keep track of the new establishment time *t*_1_, since the trajectory depends on when in the environmental cycle a lineage established. The same pattern will hold for the offspring of the new genotype, and so on, such that the frequency of the genotype with *k* additional beneficial mutations is

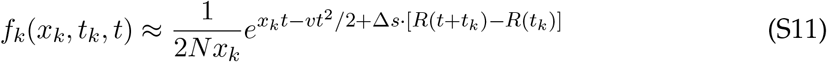

*t* generations after it establishes. As shown in Ref. (42), we can write recursive relations for *x*_*k*_ and *t*_*k*_ which depend on the random variable *T*_*k*−1_, the number of generations between *t*_*k*_ and *t*_*k*−1_:

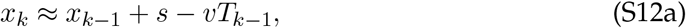

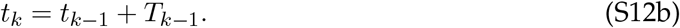

The last two terms on the RHS of Eq. (S12a) have a simple interpretation: it takes *T*_*k*−1_ generations for the focal lineage to produce a mutation with fitness effect *s*, during which time the rest of the population increases its average fitness by *vT*_*k*−1_. A successful lineage will tend to increase its “rolling fitness” *x* over many successive establishments, until the lineage reaches high frequency in the population and achieves fixation. Conversely, typical lineages will be outcompeted by the rest of the population, until *x* falls to the point that the specialist lineage goes extinct. Thus, the fixation probability of the specialist mutation is equivalent to the probability that *x* tends to increase over the long term, marginalizing over the distributions of *T*_*k*_. Most of the complexity in this process is therefore hidden in the random variables *T*_*k*_, which we discuss in the upcoming sections.

#### 2.1.2 Long-term Rolling Fitness with Environmental Tradeoffs

The rate at which a genotype described by Eq. (S11) establishes new beneficial generalist mutations is

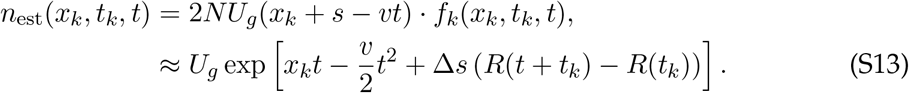

To uncover the key effects of environmental fluctuations, it is sufficient to consider only the *typical* values of *T*_*k*_. This typical waiting time (which we will also denote by *T*_*k*_ for convenience) is defined as the time when the expected number of mutations founded becomes order unity,

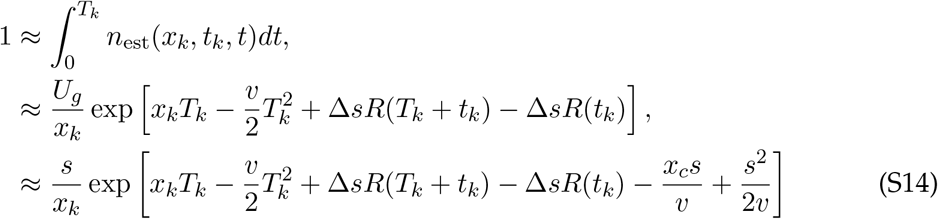

after plugging in the relationship in Eq. (S3). Neglecting the ~*s/x*_*c*_ logarithmic prefactor, the solution to this equation satisfies

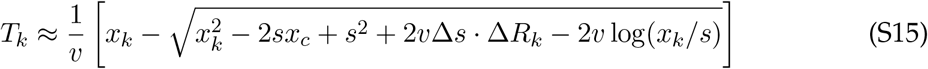

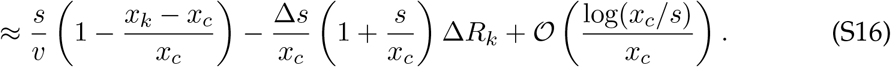

for |*x*_*k*_ − *x*_*c*_ | ≪ *x*_*c*_, where Δ*R*_*k*_ ≡ *R*(*t*_*k*+1_) − *R*(*t*_*k*_). Plugging this into Eq. (S12a) yields a solution to the recursive rolling fitness 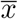 over successive foundings:

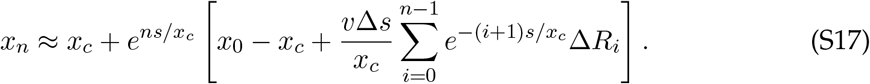

Because the Δ*R*_*i*_ depend on *t*_*i*_, this solution is still not fully explicit. In the main text, we make the additional assumption that *τ* ≫ *s/v*, so that *t*_*k*_ and *t*_*k*_ + *T*_*k*_ will usually be in the same environmental cycle. In this case, we can simplify Δ*R*_*i*_ ≈ (*s/v*) · *I*(*t*_*i*_). However, a more general approach is to expand the sum

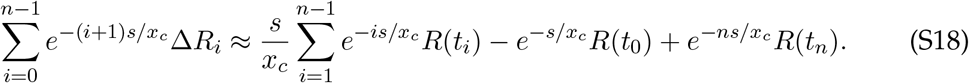

The fate of the focal lineage will be determined at least a coalescence time after it arises, *n* ≳ *x*_*c*_*/s*. In this case, we can neglect the third term and approximate the slowly decaying sum in the first term as converging to its average, 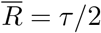. This yields

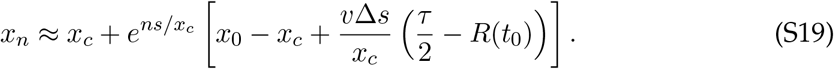

Fixation will be likely if the contents of the brackets in Eq. (S19) are positive, so that *x*_*n*_ increases with time. Thus, the tradeoff component of the specialist mutation acts like an effective shift in the initial geometric mean fitness *x*_0_, which depends on the time the lineage arises:

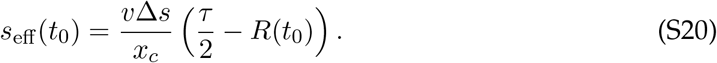

A specialist mutation with mean fitness effect 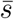 and tradeoff Δ*s* will therefore have a similar fixation probability to a generalist mutation with mean fitness effect 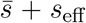 and no tradeoff. Lineages which establish at the optimal time (*R*(*t*_0_) = 0, at the start of a beneficial epoch) will be capable of fixing even if they start at a slightly lower fitness. Conversely, specialist lineages which establish at the start of a deleterious period (*R*(*t*_0_) = *τ*) must have a higher average fitness than a corresponding generalist lineage to avoid going extinct.

The source of this time-dependence is evident in Eq. (S17). Small differences in rolling fitness compound over successive foundings (even for a generalist lineage with Δ*s* = 0), because a lineage with higher starting fitness founds further beneficial mutations faster, giving less time for the rest of the population to adapt. The influence of the environment enters as a weighted sum in Eq. (S17) which places more weight on environmental states soon after the lineage arises, because they have more time to compound in the future. Specifically, a lineage that establishes at the start of a beneficial period will found its next mutations more quickly, giving it a fitness head start which increases its rate of founding future beneficial mutations, even in a deleterious period. Thus, the fate of a specialist mutation will depend on the time it arises even many environmental cycles later, rather than the environments averaging out.

#### 2.1.3 Jackpot Mutations

Our next goal is to map the fitness shift in Eq. (S20) to the fixation probability 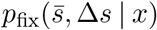 of a specialist mutation which arises on a genetic background with fitness *x*. Since we have already calculated the effective fitness shift induced by environmental tradeoffs, the remainder of the calculation closely follows previous work (42), and is consistent with past results predicting how the fixation probability depends on the fitness effect of a mutation (39). We reproduce the main features of the calculation here for clarity.

In the previous section, we employed a deterministic approximation to the mutation waiting time *T*_*k*_. This corresponds to a “step function” approximation to the fixation probability, where fixation occurs if the brackets in Eq. (S19) are positive: i.e. if *x*_0_ *>* 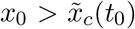 where 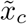 is a critical “break-even” fitness scale defined as

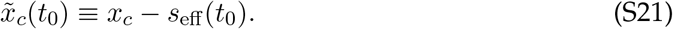

While the fixation probability indeed experiences a critical shift around this value of *x*_0_, accounting for stochasticity in *T*_*k*_ is important to predict it more accurately. This is because successful lineages can acquire atypically early “jackpot” mutations that increase their rolling fitness more than expected, allowing them to fix even if their starting fitness is slightly below 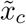.

The probability density of establishing an early jackpot mutation in only *t* generations is given by Eq. (S13). The corresponding establishment fitness is *x*_*k*+1_ = *x*_*k*_ +*s* − *vt*. Changing variables from *t* to *x*_*k*+1_, the probability of jackpotting from *x*_*k*_ to *x*_*k*+1_ is

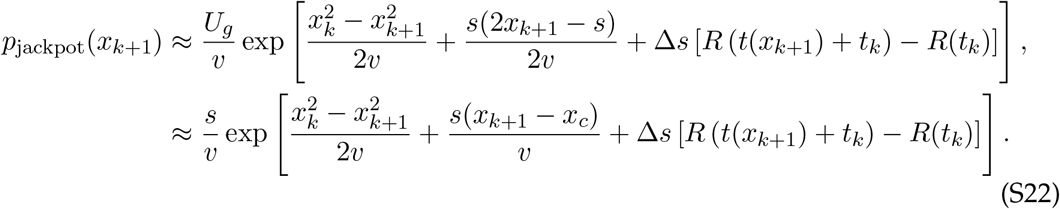

We can remove the remaining dependence on *t* by changing variables to 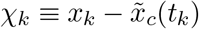, yielding

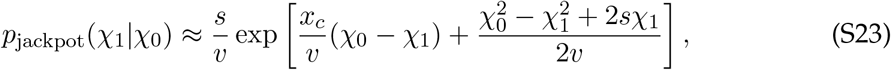

retaining only the dominant terms in the exponent. This jackpotting probability is identical to the constant-environment jackpotting probability in Ref. (42), but in terms of *χ* rather than *x*. Thus, the calculation of the fixation probability proceeds identically:

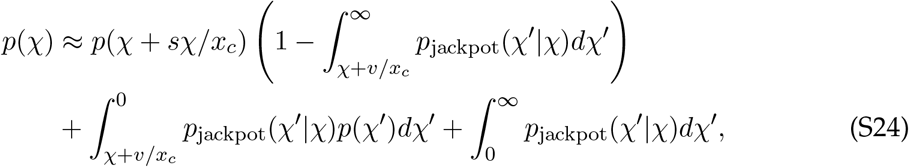

where *p*(*χ*) is the fixation probability of a lineage which establishes at shifted fitness *χ*. The first term represents the deterministic decline of a non-jackpotting lineage with *χ <* 0, the second term is the probability of jackpotting below the break-even fitness, and the third term is the probability of jackpotting above the break-even fitness (followed by deterministic fixation). The solution to this equation is

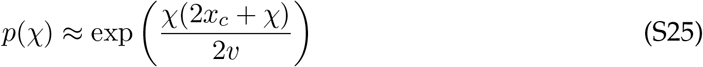

for *χ <* 0, or

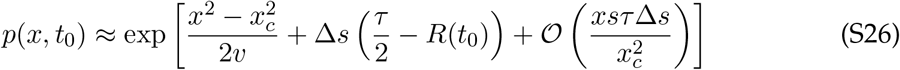

in untransformed fitness units, where *t*_0_ is the establishment time of the lineage within a cycle. Because this is the fixation probability of a lineage after establishment, the full fixation probability of a new mutation also accounts for the probability of escaping genetic drift:

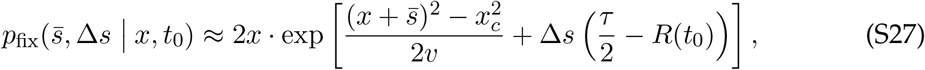

which applies when 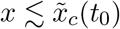.

#### 2.1.4 Overall Fixation Probability

The overall fixation probability of a specialist mutation with fitness effect 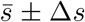 can be calculated by integrating over the starting fitness and establishment time of the mutation:

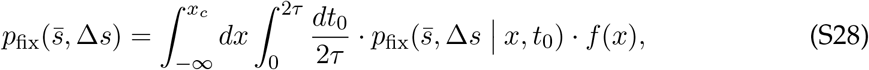

with *f* (*x*) defined in Eq. (S4) and *p*_fix_ in Eq. (S27). The dominant contribution to the integral over background fitness *x* will be 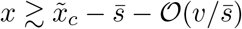, just below the break-even fitness:

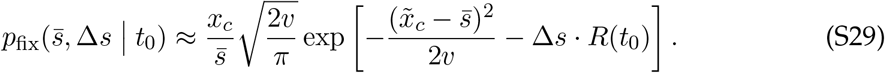

Because Eq. (S27) is only valid for 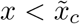, we must verify this contribution is greater than the contribution of mutations that arise above 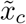 and fix deterministically after escaping genetic drift, where *p*_fix_ ≈ 2*x*. The dominant contribution in this drift-dominated region will be 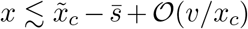, just above the break-even fitness. Ignoring smaller correction terms, we find

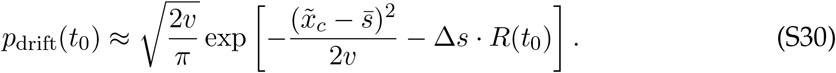

This is a factor of 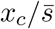 smaller than Eq. (S29), so it can be safely ignored, in which case Eq. (S29) approximates the entire integral. Finally, we can integrate over time, which will be dominated by times within 1*/*Δ*s* of *t*_0_ = 0:

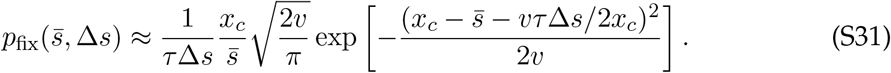

This is larger than the fixation probability of a generalist with the same geometric mean fitness effect,

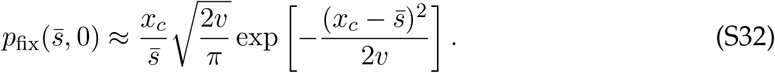

If the generalist has a different fitness effect *s*_*g*_, the log-ratio between the fixation probability of a specialist and a generalist is

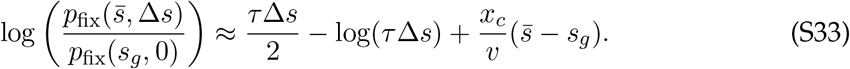

The first term reflects the increase in fixation probability for a specialist which arises at the ideal time, while the second term represents the reduced probability of the mutant appearing in this critical period. The final term simply accounts for any difference in average fitness between the mutation types, and is zero for the symmetric case 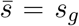. Note that while the logarithmic term is subdominant for Δ*s* · *τ* ≫ 1, it prevents the deterministic specialist advantage from becoming notable until Δ*s* · *τ* ≳ 10, broadening the transition from the weak-tradeoffs regime Δ*s* · *τ* ≪ 1.

#### 2.1.5 Asymmetric Temporal Fluctuations

Here, we briefly discuss the impact of considering a periodic environment which does not spend an equal amount of time in each environmental state (*τ*_1_ ≠ *τ*_2_). We will find that our main conclusions still hold, but the harmonic mean of the environmental durations becomes the relevant timescale.

When one environmental state lasts longer than the other, natural selection will more strongly favor mutations that increase fitness in that environment. However, this effect is already described by the geometric mean fitness defined in Eq. (S7a), which weighs each environmental state by its duration. The effect of environmental *tradeoffs* is given by the influence of Δ*s* for a fixed 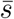. We can calculate this by generalizing the sum in Eq. (S19) to asymmetric environments, yielding

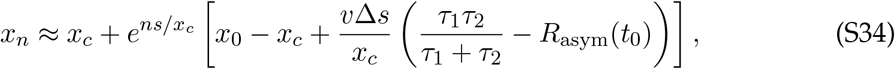

where *R*_asym_ is an asymmetric triangular wave:

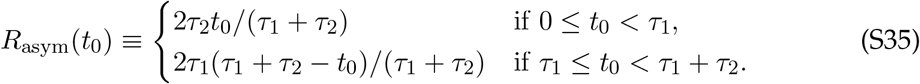

The effective fitness boost provided by the tradeoff is therefore

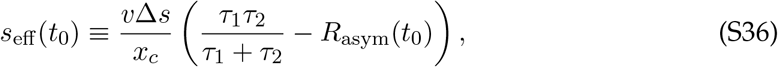

reducing to Eq. (S20) in the symmetric case.

As before, the influence of the environmental tradeoff is maximized at the beginning of each environmental state, *t*_0_ = 0 or *t*_0_ = *τ*_1_. However, the scaling of *s*_eff_ with *τ* has been generalized to the harmonic mean of the epoch durations *τ*_1_ and *τ*_2_. If the same total time *τ*_1_ + *τ*_2_ is split between environments asymmetrically, the specialist advantage will be weakened slightly. Intuitively, the limiting case is a constant environment with *τ*_2_ = 0, where geometric mean fitness 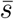 coincides with *s*_1_ only and the tradeoff Δ*s* is irrelevant at fixed 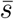. Despite these nuances, the overall takeaway is that our conclusions do not critically depend on the environmental states having identical durations; instead, environmental asymmetry simply results in a new effective timescale parameter.

### 2.2 Fixation with Seasonal Drift

In general, environmental conditions may not be perfectly periodic, and can vary stochastically over time. The fate of specialist mutations will be strongly affected by this “seasonal drift.” A simple modeling approach to capture this effect is to take the spent in each environment as an i.i.d. random variable with mean *τ* and variance 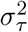. We will generally focus on the regime of modest seasonal drift (*σ*_*τ*_ ≲ *τ*) where the switching time is approximately normally distributed, although we will also discuss the impact of higher-variance distributions. (In simulations, we generally draw switching times from a gamma distribution to enforce non-negativity.)

We will find that even modest amounts of stochasticity can have enormous impacts on the specialist fixation probability, with a critical scale of *σ*_*τ*_ where the effects of stochasticity dominate over the deterministic effects discussed in the previous section. As such, we will largely neglect those deterministic effects, focusing on the seasonal drift-dominated regime. Our approach is an extension of the “rolling fitness” analysis discussed above for deterministic environments (*σ*_*τ*_ = 0).

#### 2.2.1 Rolling Fitness with Seasonal Drift

In this section, we will calculate the fixation probability of specialist mutations in a randomly fluctuating environment. To do so, it is important to consider the relative timescale of founding a new mutation compared to environmental change. If environmental epochs are the longer timescale (*τ* ≫ *s/v*), then seasonal drift over a single environmental cycle will have significant impacts on a lineage’s rolling fitness. If environmental cycles are faster (*τ* ≪ *s/v*), then the effect of seasonal drift on rolling fitness is not felt until a lineage actually produces a beneficial mutation – meaning that the net effect of seasonal drift over many cycles is the relevant “unit” of stochasticity. However, it will turn out that both regimes give identical results for the fixation probability under the normal approximation we will employ.

##### Longer environmental cycles

To start, we will assume that environmental epochs are the longer timescale, *τ* ≫ *s/v*. Consider a lineage with relative fitness *δx* ≡ *x* − *x*_*c*_ at the start of an environmental cycle. Over the cycle, seasonal drift means the time spent in the beneficial and deleterious environmental states will not perfectly balance out. We can express the impact of this imbalance on rolling fitness by assuming that the lineage takes *n*_1_ founding steps in a beneficial environment, followed by *n*_2_ founding steps in a deleterious environment. Following the recursion relations in Eq. (S12a), its rolling fitness after this process will be

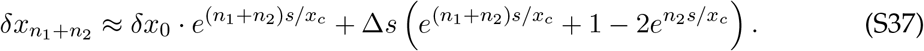

The difference between *n*_1_ and *n*_2_ comes from a random difference *δτ* between the environmental epoch times. For relatively weak stochasticity |*δτ*| ≪ *τ*, we can approximate *n*_1_ + *n*_2_ ≈ 2*vτ/s* and *n*_1_ − *n*_2_ ≈ *vδτ/s*. Plugging these in and expanding the exponentials gives us a new recursion relation in terms of *δτ*,

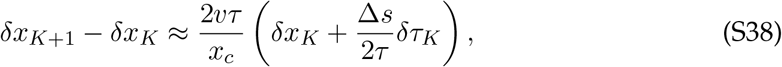

where the capital *K* indices represent environmental cycles, during which many new fitness classes are founded. We see that Eq. (S38) takes the form of a biased random walk, where the first term on the RHS deterministically pushes *δx* away from 0, and the second term introduces a random fluctuation *δτ*_*K*_ each environmental cycle. The solution to Eq. (S38) can be expressed as

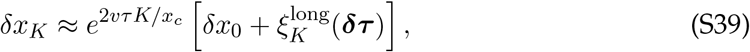

with *ξ* a random fitness scale dependent on the realized environmental stochasticity:

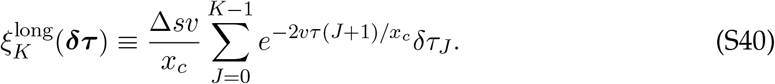

Similarly to *s*_eff_ in the deterministic case, 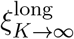 acts as an effective boost to the lineage’s initial fitness. Incorporating the effects of early jackpot mutations as discussed in SI 2.1.3, the fixation probability is

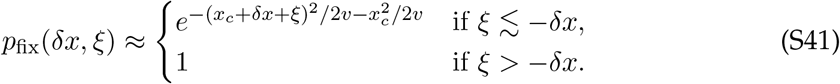

Thus, the focal lineage is likely to reach fixation if stochasticity boosts the lineage above its nominal break-even fitness. In particular, seasonal drift allows lineages with *δx*_0_ *<* 0 to nonetheless have a significant chance of fixation if *δx* is comparable to the typical scale of *ξ*. In other words, a lower-fitness lineage can be brought to fixation by a lucky sequence of longer favorable environments.

##### Shorter environmental cycles

In the *τ* ≪ *s/v* regime, we consider a similar random walk process, but where the timescale of a “step” is now *s/v* rather than *τ*. The impact of seasonal drift over this time now includes stochastic effects from many environmental epochs with varying length, yielding a rolling fitness update rule

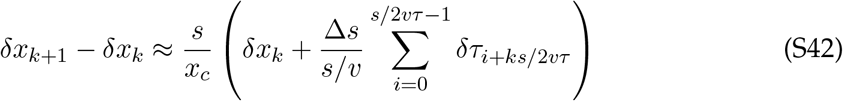

with solution

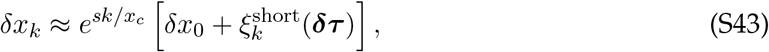

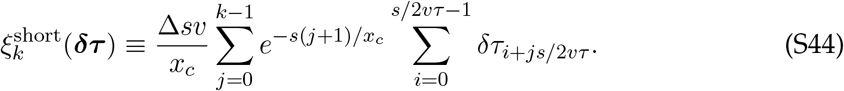

Besides the new effective value of *ξ*, the fixation probability exactly maps onto Eq. (S41). Notably, both 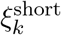 and 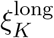 have mean zero and the same width at long times:

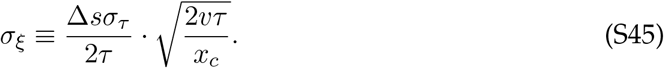

This width corresponds to the scale of the lineage’s *realized* fitness averaged over the ~ *x*_*c*_*/*2*vτ* cycles it takes to reach fixation. The first factor represents the fitness effect of stochasticity over one environmental cycle, while the second factor attenuates the stochasticity by averaging over the many independent cycles a lineage experiences before fixation.

##### Averaging over seasonal drift

In both regimes, the overall specialist fixation probability is given by averaging over the background fitness *x* and the stochasticity-induced fitness shift *ξ*. Suppose the distribution of *ξ* is *ρ*_Δ*s*_(*ξ*), which depends on Δ*s* through *σ*_*ξ*_. Then the fixation probability is

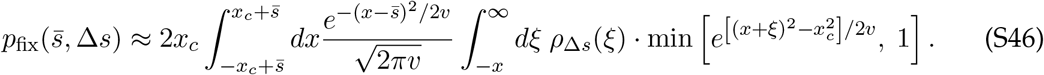

While the full distribution of *ξ* is complicated, it depends on a weighted sum of many independent random variables, so in its bulk it is reasonable to invoke the central limit theorem and approximate *ξ* ~ *σ*_*ξ*_ *N* with *N* a standard normal random variable. However, integrating out to the tail of *ξ* represents averaging over many independent environmental realizations, which may not be appropriate for realistic experimental setups where fixations are observed over a limited number of environmental epochs. Furthermore, the extreme tail of *ξ* is sensitive to the underlying distribution of *δτ*, and is not always well-described by the central limit theorem. A clear limiting case is that the effective benefit *ξ* can be no more than Δ*s*, the fitness benefit from a permanent beneficial environment. (This is evident from Eq. (S37) before expanding for |*δτ*| ≪ *τ*.) To account for these effects, it is useful to include an explicit cutoff *c* for the stochasticity,

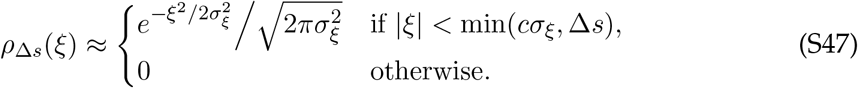

Plugging this into Eq. (S46) reveals that environmental stochasticity can provide a huge boost to the fixation probability for lineages with tradeoffs. This is because lucky environmental stochasticity (large *ξ*) can allow a lineage to reach fixation even on a lower-fitness background (smaller *x*), providing an alternate route to fixation that can increase *p*_fix_ by many orders of magnitude.

When boundary effects in the stochasticity are negligible, then the integral in Eq. (S46) will be dominated by *ξ* near 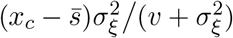, and we obtain

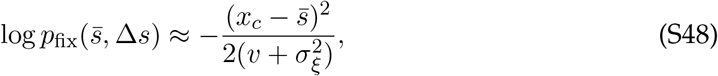

reducing to the generalist fixation probability when 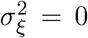. If stochasticity is relatively weak, 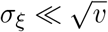 (equivalently, 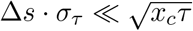) then the fixation probability simplifies to

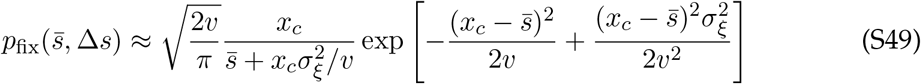

to leading order in the exponent. Thus, the specialist advantage becomes noticeable when the effect of stochasticity over a single cycle is above a critical scale, 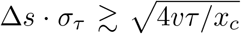. Since *vτ/x*_*c*_ ≪ 1, this scale can be much less than unity, implying that even small levels of seasonal drift strongly influence selection. This counterintuitive effect arises because the possibility of lucky environmental fluctuations expands the range of background fitnesses that have an appreciable chance of reaching fixation. Because fixation is so difficult to begin with in the clonal interference regime, an additional source of stochasticity acts like a strong benefit, even though the environmental fluctuations are not systematically biased toward beneficial environments.

The full numerical integral in Eq. (S46) with the truncated normal approximation in Eq. (S47) is in good agreement with the specialist fixation probability measured from simulations. In particular, simulation results agree with the large-*c* limit of Eq. (S47), indicating that our ~ 10^5^ simulated mutation trajectories sufficiently sample the relevant range of environmental stochasticity. However, this thorough sampling is not required for our qualitative results: even if we restricted our sampling to environments within a single standard deviation of the mean (as might be more realistic for experiments), we still predict that specialist mutations should fix 10^2^ − 10^4^ more times as often as generalists (Fig. S1). (For our isocline results in Fig. 6, we find that a cutoff *c* = 3 is most accurate, which nearly saturates the stochasticity in Fig. S1 anyway.)

The effects of seasonal drift will dominate the deterministic specialist advantage in Eq. (S33) when

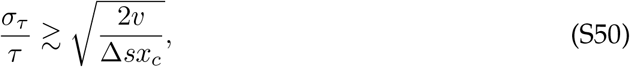

meaning that stochastic effects are likely to be relevant even when the overall scale of stochasticity is not very large. Note that our normal approximation sometimes underestimates the fixation probability slightly, likely because our simulated *δτ* possess wider exponential tails. Even wider-tailed distributions will also make the normal approximation to *ξ* suspect, as discussed in the next section.

#### 2.2.2 Higher-Variance Environments

The normal approximation will be most sensible if fixations are dominated by accumulating a fitness advantage over many independent environmental cycles, rather than a single extremely long environmental state. We will briefly discuss how this assumption depends on the tail statistics of *δτ*, and the properties of the fixation probability in the higher-variance regime.

Suppose that the probability distribution of *δτ* (the difference in duration between two adjacent environmental states) has a tail which is given by a compressed exponential,

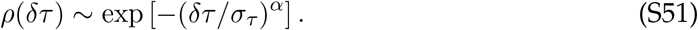

This corresponds to a Gaussian when *α* = 2 (used in the normal approximation) and an exponential when *α* = 1 (used in simulations with gamma-distributed times). The probability of drawing an extremely long time difference of at least *δτ* ^∗^ will have a similar scaling ~ *ρ*(*δτ* ^∗^), up to algebraic factors.

Now (specializing to *τ* ≫ *s/v*), we will consider the probability a lower-fitness lineage is set on a deterministic path to fixation by a *single* random walk step with an abnormally large *δτ*. For a lineage which starts at fitness *x*_*c*_ +*δx*, this roughly requires *δτ >* |*δx*|*x*_*c*_*/*Δ*sv*. Integrating over −*δx*, we write

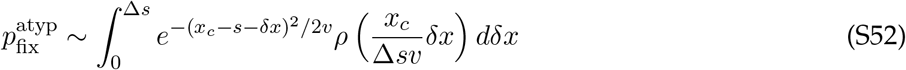

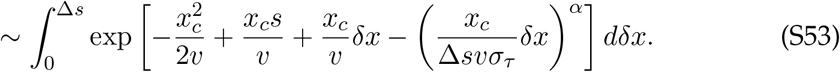

When *α >* 1, the integral will have a saddle point at *δx*^∗^ ~ (*v/x*_*c*_)(Δ*sσ*_*τ*_)^*α/*(*α*−1)^ (up to *α*-dependent prefactors), yielding

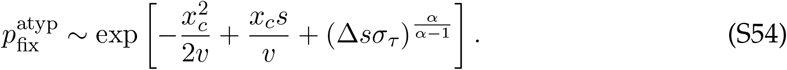

when 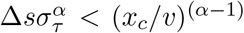. For larger stochasticity, the integral will be dominated by its upper bound, yielding

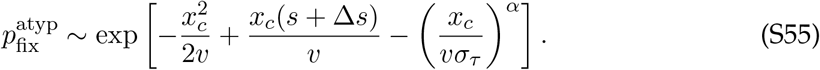

This atypical path to fixation entails an effective fitness benefit of ~ (*v/x*_*c*_)(Δ*sσ*_*τ*_)^*α/*(*α*−1)^, generically weaker than the normal approximation unless *α* is close to unity and Δ*sσ*_*τ*_ ≳ 1.

For *α* = 1 (an exponential tail), the relevance of rare environmental cycles is marginal, depending on the size of Δ*sσ*_*τ*_. When Δ*sσ*_*τ*_ *<* 1, randomness across many environmental cycles will dominate; when Δ*sσ*_*τ*_ *>* 1, the most likely path to fixation will be lineages ~ Δ*s* below the break-even fitness experiencing an abnormally long beneficial environment. This effect likely explains the fact that the normal approximation can underestimate the specialist fixation probability in simulations (Fig. 3C), where Δ*sσ*_*τ*_ ≳ 1. However, the quantitative predictions are still broadly reasonable even up to Δ*sσ*_*τ*_ ~ 10, likely because the effective fitness benefit is capped at Δ*s* in both cases, yielding similar results for higher stochasticity.

For wider-tailed distributions (e.g. *α <* 1), the normal approximation is suspect, but may still apply for sufficiently small Δ*sσ*_*τ*_. We consider a simple high-variance distribution in Fig. S2, where rare environmental states last an extremely long time. In this case, the fixation probability is dominated by the probability of these rare environmental states, where specialist mutations experience an effective benefit of Δ*s*.

We emphasize that the approach taken in this section is critically dependent on the assumption that specialists are rare, such that only one lineage at a time can take advantage of environmental fluctuations’. Outside of this regime, many lineages can simultaneously respond to environmental fluctuations, and their fixation advantage will be reduced. We explore this effect in SI 3.3.

## 3 Dynamics with Frequent Specialists

### 3.1 Introduction

Our analysis so far has focused on the case where specialist mutations with Δ*s >* 0 arise rarely, so that only one specialist lineage is present in the population at any given time. In this section, we will relax this strong assumption, using heuristic methods to capture an overall picture of how the fates of mutations are determined in a population where specialist mutations are common. Relative to our discussion of rare specialists, this analysis will focus on a more specialized regime, and neglect some quantitative details which would be compelling to further explore in future work. However, our approach will nonetheless reveal many of the most interesting ways frequent specialist mutations influence population dynamics, and allow for qualitative and quantitative agreement with key features of simulations.

When the specialist mutation rate is large, *NU*_*s*_ ≫ 1, the background population will adapt just like the focal strain does, forming a fully two-dimensional fitness distribution. We parametrize this distribution in terms of geometric mean fitness *X*_∥_ and specialism *X*_⊥_, defined such that a mutation with tradeoff strength Δ*s* shifts *X*_⊥_ to *X*_⊥_ + Δ*s*. In the time-symmetric case *τ*_1_ = *τ*_2_ (which we assume henceforth), the instantaneous fitness of a genotype is simply *X*_∥_ + *I*(*t*) · *X*_⊥_. Its relative fitness is given by 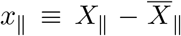 and 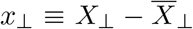, where bars represent the population mean. We make a linear ansatz for how these population averages vary with time:

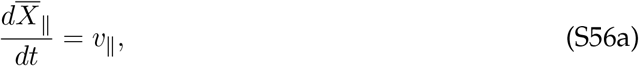

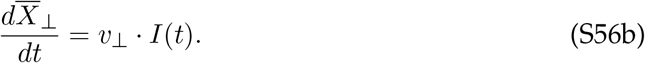

Under this ansatz, the instantaneous mean fitness of the background population grows at rate *v*_∥_ + *v*_⊥_ as it specializes to one environment, but drops discontinuously when the environment switches, such that the long-term rate of adaptation is only *v*_∥_. The “rate of specialization” *v*_⊥_ represents cyclic specialization to the current environment which is lost in the next environmental state. Both *v*_∥_ and *v*_⊥_ are now unknown parameters which must be determined self-consistently. In particular, *v*_∥_ may not take the same value as *v* in the rare-specialists case, although we will find that they agree to leading order.

The dynamics in Eq. (S56) can be thought of as a linear approximation to the true dynamics of the bulk population, which is sufficient to estimate the key behavior to leading order. One neglected feature of the dynamics is that over long timescales where multiple specialist mutations fix, the number of specialist mutations of each type will not perfectly balance, so *X*_⊥_ will drift stochastically from its original value. However, because fitness is always measured relative to the rest of the population, this drift should have minimal effects over the timescale of a single mutation reaching fixation or going extinct.

The next key effect of allowing frequent specialist mutations is that any focal lineage has access to multiple types of mutations, which change environmental specialism in addition to average fitness. This means that the fate of a mutation must be determined by aggregating over all the possible mutations its descendants could acquire. We can resolve this complexity by noting that only a small fraction of these descendants are likely to reach fixation – those that follow the “optimal mutational path,” which depends on environmental fluctuations. In our analysis of rare specialists, we found that that specialist mutations aligned with the current environment lose their advantage halfway through an environmental state, so we might expect this optimal path to switch the type of mutation it gets at roughly the same time. Our rolling fitness framework will allow us to formalize this intuition.

### 3.2 Frequent Specialists in a Periodic Environment

We will first consider specialist mutations in a perfectly periodic environment (*σ*_*τ*_ = 0). For ease of presentation, we will start with the symmetric case *U*_*s*_ = *U*_*g*_ and 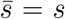, where specialist mutations arise at the same rate as generalists and have the same average fitness effect. The more general case is discussed in SI 3.2.7. We will also specialize to the *τ* ≫ *s/v*_∥_ regime, where many new fitness classes are founded each environmental cycle. (Recall that our main results in the rare-specialists regime did not critically depend on this choice, but were often more dramatic for larger *τ*, making this regime easier to explore in simulations.) Finally, we will see that the frequent-specialists regime is sensitive to the relative sizes of Δ*s* and *s*, which determines whether specialist mutations founded in the “wrong” environment have a higher initial growth rate than their parent strain. We will focus on the *s >* Δ*s* case in our analytic theory, but discuss the Δ*s > s* case in SI 3.2.8.

#### 3.2.1 Rolling Fitness of a Specializing Lineage

Consider a lineage which establishes at time *t*_0_ at relative geometric mean fitness *x*_∥_ and relative specialism *x*_⊥_. Its frequency trajectory will be

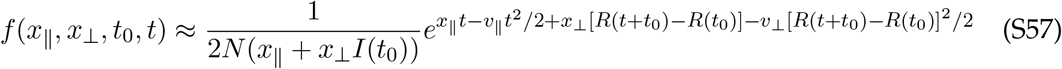

We can follow the same procedure as in SI 2.1.2 to calculate the typical time it takes this lineage to produce a mutant,

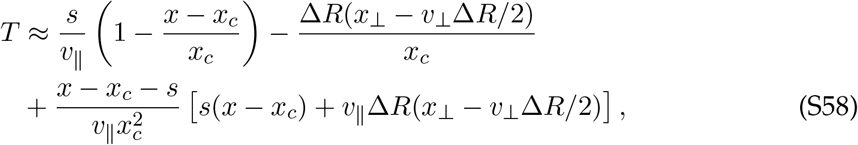

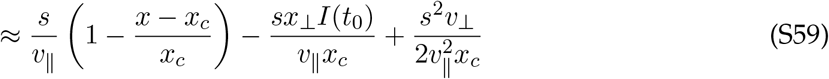

where Δ*R* ≡ *R*(*T* + *t*_0_) − *R*(*t*_0_) = *I*(*t*_0_) · *T* in the *τ* ≫ *s/v*_∥_ regime. This equation reduces to Eq. (S16) for *x*_⊥_ = Δ*s* and *v*_⊥_ = 0. Importantly, we have defined *x*_*c*_ as a large fitness scale analogous to Eq. (S3),

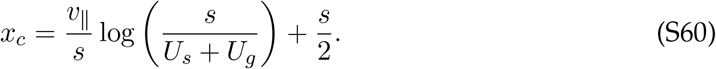

Note that our perturbative expansion of *T* assumes *x*_⊥_ ≪ *x*_*c*_, which may not be valid if Δ*s* is sufficiently large. We will compare relevant values of *x*_⊥_ to *x*_*c*_ later, and find that they are generally smaller than *x*_*c*_ if *s >* Δ*s*.

When the next fitness class is founded, the update rule for *x*_∥_ is the same as before, but now *x*_⊥_ can also change due to adaptation in the rest of the population, and new specialist mutations in the focal lineage. The resulting update rules are

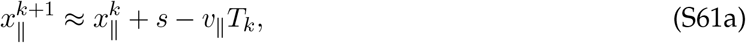

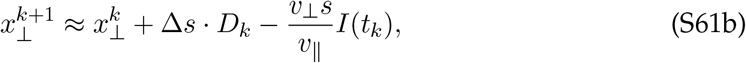

where *D*_*k*_ ∈ (− 1, 0, 1) parametrizes the type of the new mutation: 0 for a generalist, and ±1 for each type of specialist. Crucially, these update rules are only valid when *s >* Δ*s*, which guarantees that each new fitness class is dominated by descendants of the *first* mutation into that class (because daughter strains always grow faster than their parent). Under this assumption, the solution to these recursive relations is

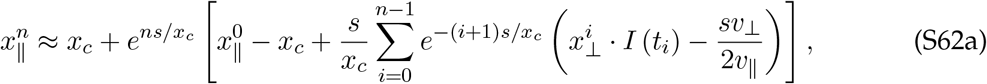

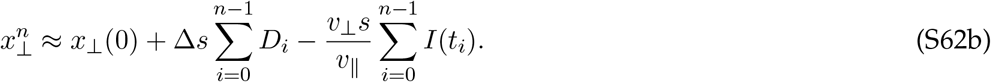

It is useful to plug in 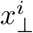 explicitly and separate out the portion of 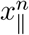 dependent on the mutational path,

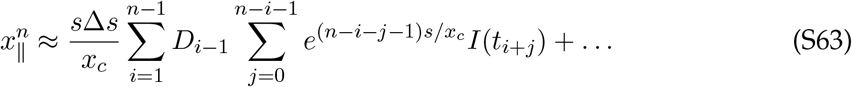

with omitted terms independent of ***D***.

#### 3.2.2 Optimal Mutational Path

Each combination of *D*_*i*_ represents a subset of the focal lineage that has taken a particular mutational path (i.e., a specific order of types of mutations). However, we expect the fixation probability of the lineage to be dominated by a particular mutational path which maximizes long-term fitness. Since Eq. (S63) depends on each *D*_*i*_ linearly, we can write this optimal path directly:

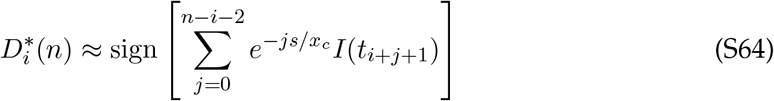

We see that 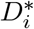 is based on a weighted sum of the environmental states between *t*_*i*_ and *t*_*n*−1_, with more importance placed on the times soon after the mutation established. Crucially, Eq. (S64) depends on *n*, the “observation time” of the fitness 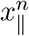. This dependence indicates that what appears to be optimal at one time may not be optimal over the long term. For example, a mutation acquired near the end of a favorable environment may appear optimal at first, but quickly be outcompeted by other mutational paths when the environment changes.

By plugging in the periodic structure of *I*(*t*), we can express the sum in Eq. (S64) as a term going through *n*_cyc_ ≡ floor (*n* − *i* − 1)*/*(2*v*_∥_*τ/s*) complete cycles, followed by an “overhang” term based on the phase difference between *t*_*i*_ and *t*_*k*_:

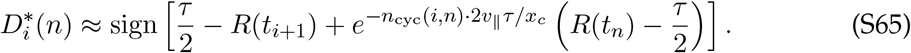

We see that the optimal mutation depends on the time the mutation establishes (*t*_*i*_) as well as the observation time (*t*_*n*_), with the importance of the latter term decaying if the mutation happened many cycles in the past. When this term is negligible (*n*_cyc_ ≳ *x*_*c*_*/*2*v*_∥_*τ*), the optimal mutational path reduces to the *long-term optimal* mutational path:

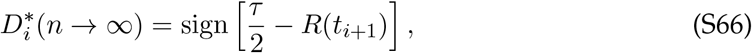

reflecting the impact of a mutation many environmental cycles into the future. This exactly mirrors our intuition from the rare-specialists case: it is best to acquire mutations favoring the current environment only until it is halfway over. Afterward, it is best to switch to mutations favoring the upcoming environment.

##### Effective fitness shift

The long-term fixation probability of a lineage following the optimal mutational path will be determined by its initial mean fitness 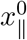 plus a fitness shift *ζ* given by the last term in the brackets of Eq. (S62a):

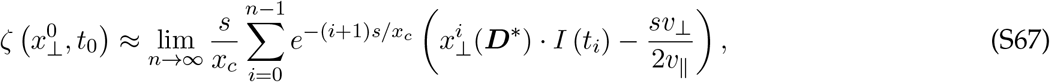

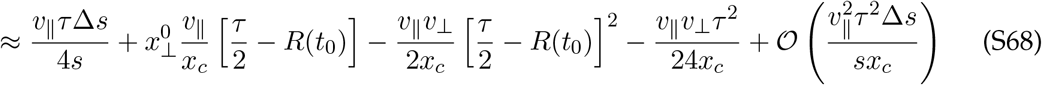

This choice of neglected terms will be valid if *v*_⊥_ ≫ *v*Δ*s/s*, which we will verify later. The first constant correction represents the effective benefit that a lineage acquires by following the optimal mutational path, providing a benefit proportional to Δ*s* (the tradeoff strength of available specialist mutations). The 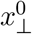-dependent term has exactly the same form as the rare-specialists regime, representing the time-dependent benefit (or cost) a lineage has from having specialized already. Finally, the *v*_⊥_-dependent terms represent the cost of competing against a specializing background population. Note that this cost is minimized in the middle of an environmental state, where the optimal path begins to diverge from the direction the background population is specializing.

#### 3.2.3 Two-Dimensional Traveling Wave

In order to estimate the fixation probability of a mutation, we must characterize the two-dimensional fitness distribution of the population *f* (*x*_∥_, *x*_⊥_). We will find that *f* is a two-dimensional traveling wave which moves according to Eq. (S56). Our central approach is to realize that all extant organisms in the population must be descended from a common ancestor in the distant past, whose dynamics when small were given by the rolling fitness calculation we just performed. Thus, our update rules for a focal lineage can be used to self-consistently determine the entire two-dimensional fitness distribution of the population. In the process of performing this calculation, we will also be able to estimate the rate of specialization *v*_⊥_, which we address first.

##### Rate of specialization

The rate of specialization is encoded in the optimal mutational path we found earlier, Eq. (S65). This path corresponds to the organisms at the highest frequency in the high-fitness nose of the population at time *t*_*n*_, which will eventually rise in frequency until they form the bulk of the wave. The specialism *x*_⊥_ of this genotype can be inferred by aggregating over all the mutations it has gained:

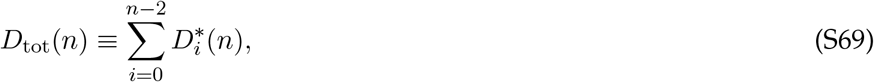

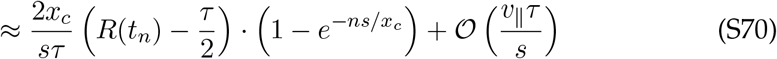

since *n* ~ *x*_*c*_*/s* ≫ *x*_*c*_*/*2*v*_∥_*τ* is large enough to span many environmental cycles. Crucially, this sum is highly sensitive to the observation time *t*_*n*_: unlike the long-term optimal mutational path, the short-term optimal mutational path entails strongly specializing toward the *current* environment. We see that the average specialism in the high-fitness nose changes at rate Δ*s* · 2*x*_*c*_*/sτ*. We will show that this rate of change in the nose feeds back to set the rate of specialization in the bulk,

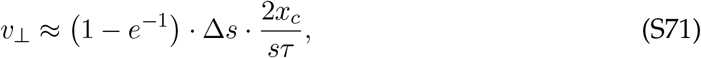

taking *n* = *x*_*c*_*/s* to cut off the sum at a coalescence time, which improves agreement with simulations (Fig. S3B). Thus, longer environmental cycles tend to limit the diversity in environmental preferences present in the population, lowering *v*_⊥_ according to Fisher’s Fundamental Theorem. Because *v*_∥_*τ/x*_*c*_ ≪ 1, the asymptotic limit of *v*_⊥_ is large enough to satisfy our assumption that *v*_⊥_ ≫ Δ*sv*_∥_*/s*.

Simulations verify this rough dependence of *v*_⊥_ on Δ*s* and *τ*, although its value quantitatively approaches Eq. (S71) only for extremely large population sizes when both *v*_∥_*τ/x*_*c*_ and *s/v*_∥_*τ* are very small (Fig. S3). Simulated values of *v*_⊥_ are generally less than the asymptotic limit, meaning that it acts as an upper bound. In practice, we often measure *v*_⊥_ from simulations and treat it as an input rather than relying on this leading-order analytic result.

##### Frequency distribution

We will now describe the frequency distribution of the population as a whole, showing that it is consistent with the rate of specialism we found in Eq. (S71). The peak of this frequency distribution in the nose is given by the optimal path in Eq. (S70), which predicts that the peak relative specialism 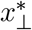 should obey

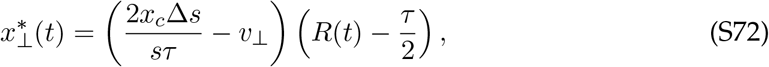

where we are once again treating *v*_⊥_ as an unknown parameter. We would like to characterize not only the location of this peak, but also the shape of the frequency profile in the nose near 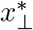. To do so, we must extend Eq. (S63) to consider deviations from the optimal mutational path, which will form the frequency profile around the peak.

The coefficient of *D*_*i*_ in Eq. (S63) can be written as

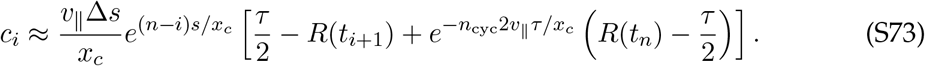

The rolling fitness cost of sign-flipping *D*_*i*_ from its optimal value is 2*c*_*i*_. To find the least costly way of flipping a fixed number *M* of mutations, we define a fitness threshold *z* and flip all mutations with a cost of less than *z* per mutation. Over a single complete cycle *n*_cyc_ cycles before the present, this will result in flipping

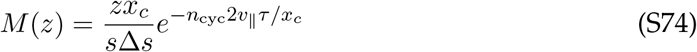

mutations. The total fitness cost of flipping these mutations will be

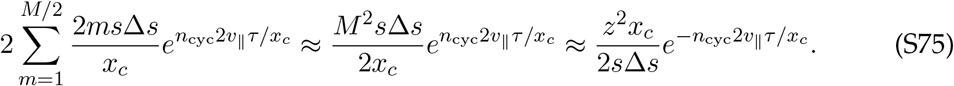

Aggregating over all cycles (i.e., summing over *n*_cyc_) yields a total number of flips

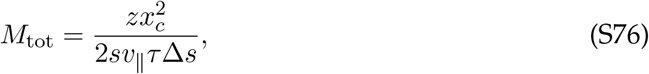

resulting in a total change in specialism

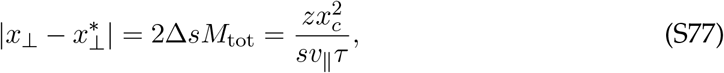

at a rolling fitness cost

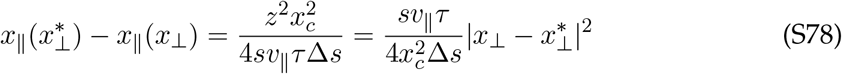

relative to the optimal path, where we have rewritten dependence on *z* in terms of 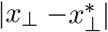. These equations describe how the rolling fitness of organisms in the nose depends on their environmental specialism *x*_⊥_.

Next, we will change variables in Eq. (S78) from *x*_∥_ to frequency in the population. A lineage whose rolling fitness is *δx*_∥_ larger than another’s must have established (*δx*_∥_)*/v*_∥_ generations earlier. Lineages at the nose of the wave will have relative fitness near *x*_*c*_, rising in frequency as 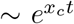 after they establish in the population. So, rolling fitness can be converted to frequency via the relation

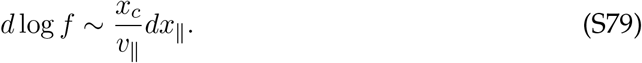

The frequency profile of organisms in the high-fitness nose is therefore

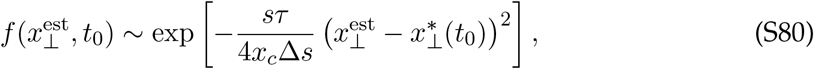

where 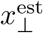 is the relative specialism of a the lineage at the time it establishes. After establishing in the nose of the wave at average fitness 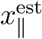 and specialism 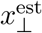, a lineage will grow in frequency over time. Its frequency *t* generations after establishing will be

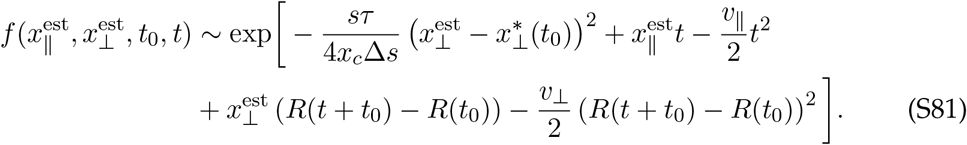

Finally, we change variables from establishment fitness to instantaneous relative fitness, substituting 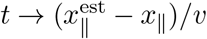 and 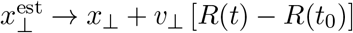. This yields

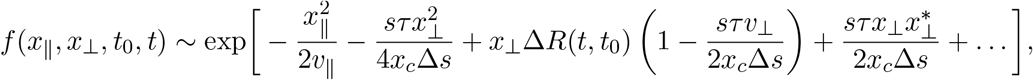

where the omitted terms are independent of *x*_∥_ and *x*_⊥_ but may depend on 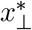 and *t*_0_. We observe that if *v*_⊥_ follows Eq. (S70), the explicit time dependence will cancel out, and 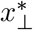 will be fixed at zero. This simplification leaves only

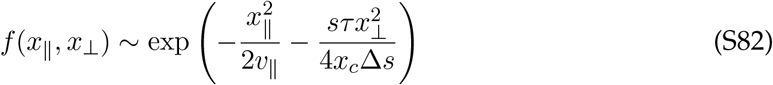

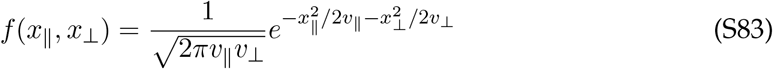

Thus, the two-dimensional traveling wave has a simple stationary Gaussian profile consistent with our original ansatz in Eq. (S56) and our expression for *v*_⊥_ in Eq. (S71). Note that because *x*_⊥_ is measured relative to the population mean, it is stationary in time even though the absolute specialism of the population fluctuates as the environment changes.

#### 3.2.4 Jackpot Mutations in the Frequent-Specialist Regime

Now that we have characterized the fitness distribution present in the population, we would like to estimate the fixation probability of a mutant. This requires accounting for the probability that a lineage obtains an abnormally early “jackpot” mutation. This analysis will closely follow the rare-specialist analysis in SI 2.1.3 and leads to essentially the same results, where a mutation’s fixation probability sharply decreases as *x*_∥_ falls below *x*_*c*_ − *ζ*.

To incorporate stochasticity in founding time, we write the rate of establishing beneficial mutations *t* generations after initial establishment as

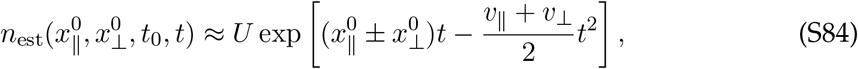

assuming that *t*_0_ and *t* are in the same environmental state, which determines the sign of 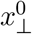 chosen. Note that we assume *U* = *U*_*s*_ = *U*_*g*_ (see SI 3.2.7 for the extension to differing mutation rates).

We first change variables from time to establishment fitness 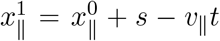, and then to 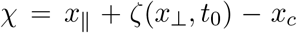 *x*_*c*_. This gives the probability an optimally-specializing lineage jackpots from *χ*_0_ to *χ*_1_:

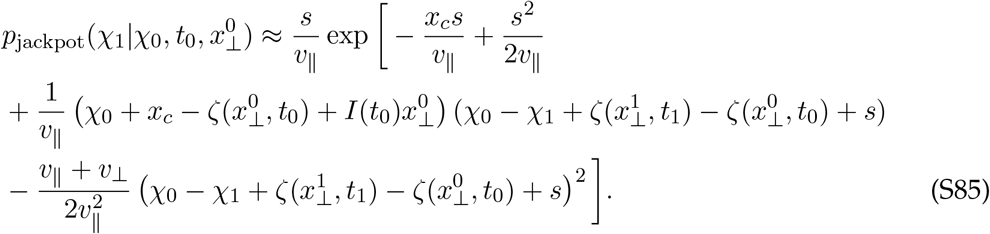

A lineage starting at the break-even fitness, *χ*_0_ = 0, will tend to land at the break-even fitness if it does not jackpot. Thus, the argument of the exponential should roughly cancel when *χ*_0_ = *χ*_1_ = 0, so we can ignore terms independent of *χ*_0_ and *χ*_1_. Next, we observe that 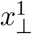 and *t*_1_ have implicit dependence on Δ*χ* ≡ *χ*_1_ − *χ*_0_ through

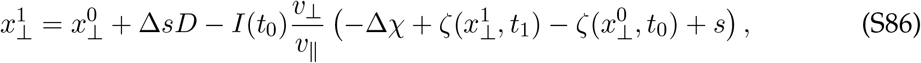

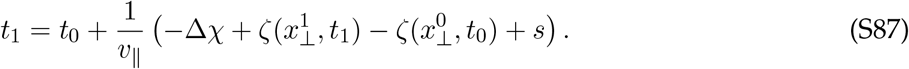

We can account for this dependence using a Taylor approximation, assuming that the jackpot is not too far from the typical waiting time for a mutation:

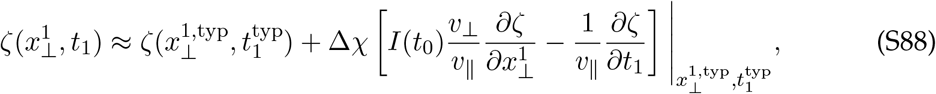

where 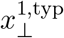 and 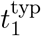 are the non-jackpotting typical values of 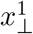 and *t*_1_, independent of *χ*_0_ and *χ*_1_. With these simplifications, we can write the leading terms in the jackpotting probability as

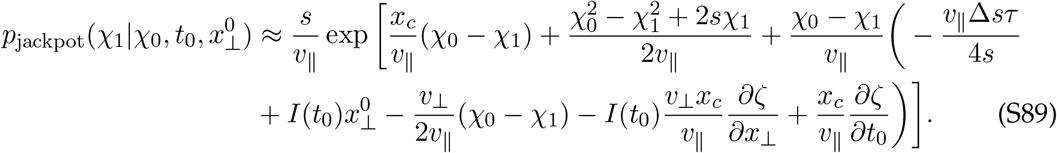

The derivatives of *ζ* are, to leading order,

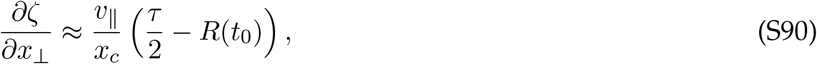

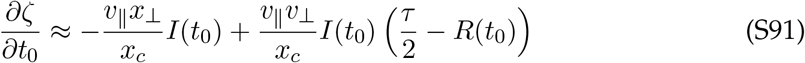

resulting in an expression identical to the rare-specialists case in Eq. (S23):

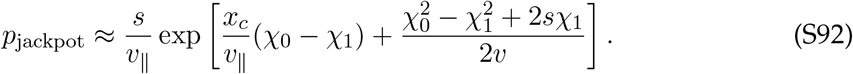

As discussed in SI 2.1.3, this results in lineages with *χ <* 0 surviving with probability

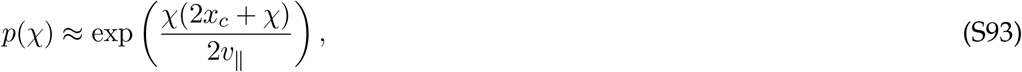

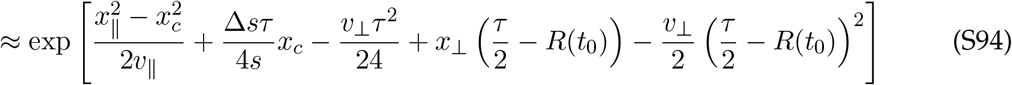

after returning to untransformed fitness units, with neglected terms of Δ*sv*_∥_*τ* ^2^*/s* in the exponent.

#### 3.2.5 Overall Fixation Probability

We can now calculate the fixation probability of a general mutation 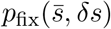 based on our estimate for the fixation probability of an established lineage on starting fitness and time, Eq. (S94), and the background fitness distribution, Eq. (S83). Note that we use 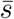 and *δs* to describe the geometric mean fitness effect and tradeoff strength of the focal mutation, which may differ from the corresponding parameters (*s* and Δ*s*) in the JDFE describing the other mutations available in the population. The fixation probability is given by an integral over backgrounds and time:

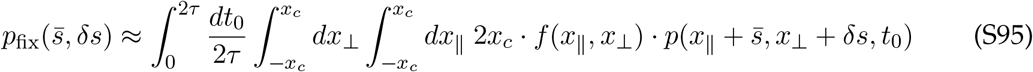

The integral over *x*_∥_ is dominated by *x*_∥_ within ~ *v*_∥_ */s* of 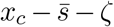, while the integral over *x*_⊥_ is dominated by *x*_⊥_ within 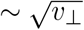 of

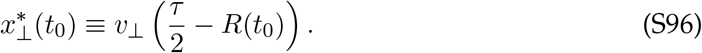

The overall time-dependent expression after performing these integrals is

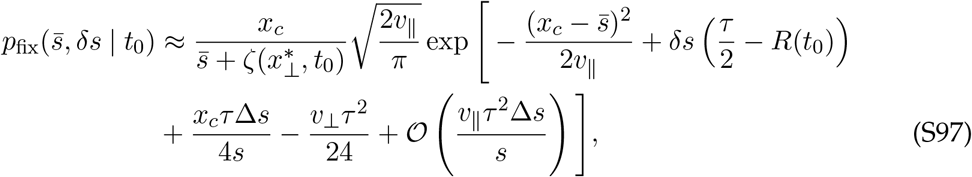

where the size of omitted terms in the exponent is set by our truncation of *ζ* in Eq. (S68), and may depend on *t*_0_ but not 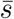 and *δs* to leading order.

Two conclusions immediately follow. First, 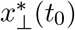 effectively cancels out the bulk population’s environmental specialization over a cycle. This means that near environmental switches, the genetic backgrounds of typical successful mutations will be “pre-adapted” to the upcoming environment relative to typical organisms. Second, the *δs*-dependence of *p*_fix_ has exactly the same form as the rare-specialists regime,

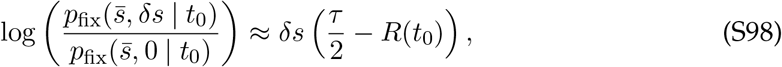

implying that the time-dependent specialist advantage has the same two-sided exponential form we obtained in the rare specialists regime.

However, calculating the time-averaged specialist advantage

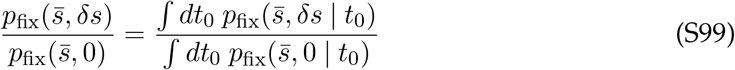

is more difficult, because of the additional time-dependence introduced by the omitted terms in Eq. (S102). These time-dependent terms are not necessarily small and can enter at similar order to the *δs* term, influencing the value of *t*_0_ that dominates each integral. Nonetheless, we can make an ansatz for the structure of these terms based on the observation that under the optimal mutational path, a successful lineage acquires specialist mutations that arise roughly uniformly over time across each environmental cycle. By enforcing this uniformity as a self-consistent constraint, we can impute the form of the omitted time-dependent terms.

Writing 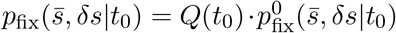, with 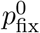 given by Eq. (S102) and *Q* the unknown correction term, we require that successful mutations arise uniformly across time:

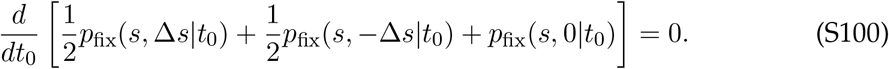

Solving this equation yields

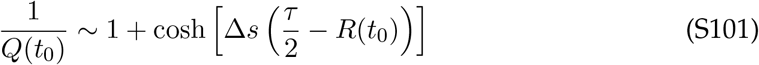

up to a time-independent constant. The overall time-dependent fixation probability is then

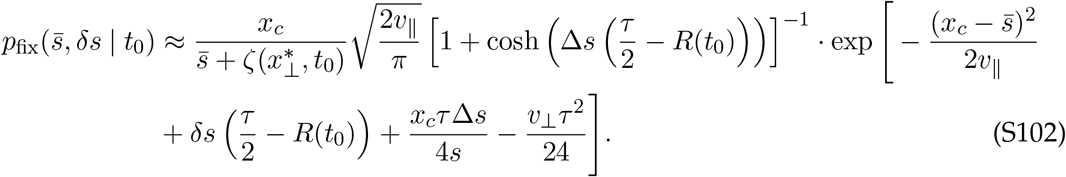

The effect of the new term is to globally reduce the fixation probability for mutations that arise near environmental switches. Intuitively, this is because mutations that arise at these times face heightened competition with specialist mutations on other genetic backgrounds. This adjustment can be observed directly in simulations (most obviously for generalist mutations, where it is the dominant time-dependent effect; Fig. S6).

The overall specialist advantage can be found by averaging Eq. (S102) over time. The time-dependence is sensitive to the relative size of *δs* and Δ*s*, determining how the tradeoff strength of the focal mutation compares to that of the JDFE. If |*δs*| = Δ*s* (a typical mutation), the fixation probability is nearly constant in the first ~ *τ/*2 generations of its favored environmental state – then rapidly shifts to favor the other type of specialist mutation, with a brief crossover period where generalist mutations are equally likely to fix as specialists. Mutations with weak tradeoffs, |*δs*| *<* Δ*s*, will act similarly to generalists with |*δs*| = 0, and are most likely to fix near this crossover region. Conversely, mutations with stronger-than-typical tradeoffs (*δs >* Δ*s*) are most likely to fix near environmental switches, and have an exponential fixation advantage because they systematically outcompete other mutations at those times:

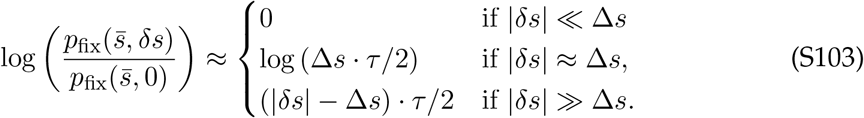

For typical mutations with *δs* = Δ*s*, the specialist advantage is now *linear* in Δ*sτ*, rather than exponential. This is because there are approximately 4*/*Δ*s* generations in an environmental cycle where specialist mutations are about as likely to fix as generalists, out of 2*τ* total. This is in contrast to the rare-specialists regime, where successful specialist mutations mostly occurred at the start of environmental epochs, where their fixation advantage is strongest. Note that this conclusion critically changes in the Δ*s > s* regime discussed in SI 3.2.8, where successful mutations are no longer evenly distributed across an environmental cycle.

In addition to the new form of the time dependence, the constant terms in the exponent of Eq. (S102) (coming from the constant term in *ζ*^∗^) are also distinct from the rare-specialists regime. The leading-order adjustment to the fixation probability is independent of *δs*,

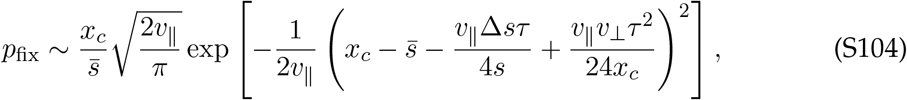

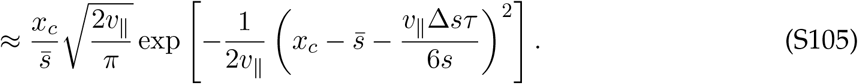

in the asymptotic limit where Eq. (S71) is valid. The new terms in the parentheses will affect the population-wide rate of adaptation *v*_∥_, discussed in the next section.

#### 3.2.6 Self-Consistent Rate of Adaptation

The presence of frequent specialist mutations will adjust the overall rate of adaptation of the population *v*, and thereby the large fitness scale *x*_*c*_, which we have defined in terms of *v* through Eq. (S60). We can calculate *v* through the self-consistent relation

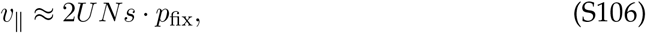

with *p*_fix_ defined in Eq. (S105). This equation is solved numerically to obtain the theory curves in Fig. S6, but we can also analytically calculate the leading-order correction to *v* by balancing the exponential terms in Eq. (S105) with *NU*_*s*_, yielding:

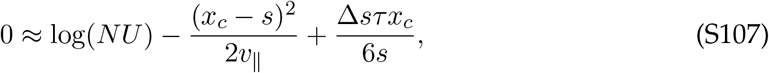

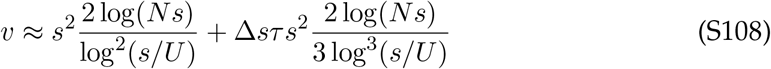

after plugging in the definition of *x*_*c*_ from Eq. (S60). We can write the frequent-specialist *v*_∥_ and *x*_*c*_ in terms of their original values in the rare-specialist regime (denoted as *v*^0^ and 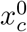):

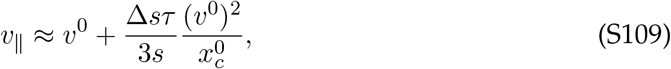

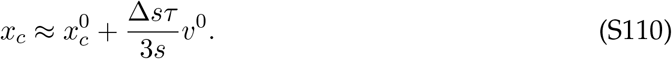

(Note that the prefactor of this adjustment term may be up to a factor of 3*/*2 larger if *v*_⊥_ is smaller than its asymptotic limit.) Thus, having access to environment-sensitive mutations allows the population to slightly increase its average rate of adaptation. This is consistent with the fact that the benefit of specialist mutations derives from their ability to speed up the rate at which a lineage founds future mutations. If all successful lineages have well-timed specialism, the bulk *v*_∥_ should be larger as well.

#### 3.2.7 Asymmetric Fitness Effects and Mutation Rates

Our analysis of the frequent-specialists regime above assumed that the average fitness effects and mutation rates of specialist and generalist mutations were equal. We will briefly discuss extending our calculation to the more general asymmetric case, focusing on specialist mutations with a lower average fitness than generalists. (Because specialist mutations are already favored over generalist mutations in the symmetric case, an additional benefit to specialist mutations can be treated as simply a higher effective value of *s* or *U*.)

Suppose specialist mutations have fitness effects (*s* − *c*, ± Δ*s*), with an average fitness cost *c* relative to generalists. Effectively, each specialist mutation a lineage acquires in lieu of a generalist mutation will impose a rolling fitness cost of *c*. By accounting for this fitness cost, we can characterize the new population dynamics.

**Long-term optimal path**. To determine the long-term optimal mutational path, we can compare the time-dependent benefit of specialist mutations to their relative cost of *c*. We will focus on the crossover regime

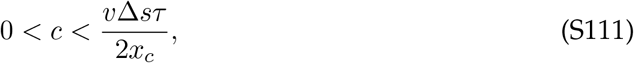

where the average fitness cost of specialist mutations is competitive with their effective benefit. (For larger *c*, generalist mutations will tend to outcompete specialist mutations over the long term.) In the crossover regime, generalist mutations will be favored near the middle of each environmental state, such that the optimal mutational path includes each type of mutation:

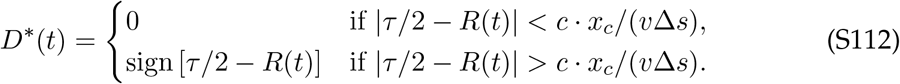

The fixation probability ratio is given by the fraction of time each mutation is optimal:

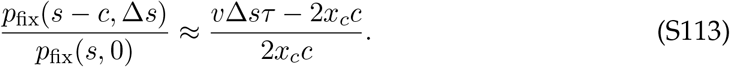

Following a similar procedure as in Eq. (S68), the effective fitness boost from following the optimal mutational path will be reduced,

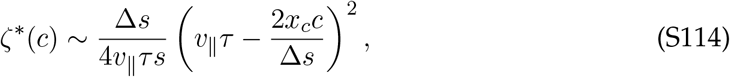

neglecting the *v*_⊥_-dependent term (which we expect to adjust numerical factors but not the overall scaling).

##### Population rate of adaptation

Asymmetric fitness effects have interesting implications for the population-wide rate of adaptation, because they modulate both the relative fixation probability of specialist and generalist mutations as well as their direct fitness effects. The rate of adaptation is defined by the equation

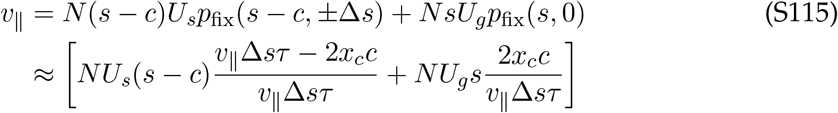

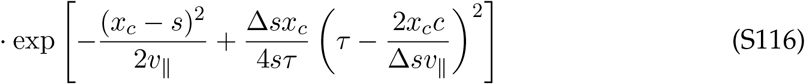

after plugging in the leading correction to the break-even fitness from Eq. (S114). When *U*_*s*_ and *U*_*g*_ are similar, the solution to this equation can be expressed as

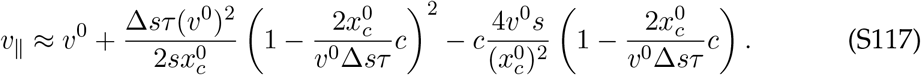

Interestingly, this result is non-monotonic in *c*. Between *c* = 0 and 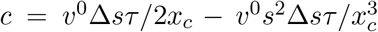, the first correction term dominates, and *v*_∥_ decreases with larger *c* as the fitness benefit of successful mutations is reduced, eventually dropping below the rare-specialists *v*^0^. But as *c* increases further toward *v*^0^Δ*sτ/*2*x*_*c*_, costless generalist mutations start to be favored by natural selection, and *v*_∥_ increases back to *v*^0^. While the non-monotonic range is small in the deterministic regime (*σ*_*τ*_ = 0), this basic behavior will also apply in the stochastic *σ*_*τ*_ *>* 0 case, where the non-monotonicity of *v*_∥_ can be easily observed in simulations.

##### Differing mutation rates

We can now easily extend our results to the *U*_*g*_ ≠ *U*_*s*_ case by observing that if *U*_*g*_ *> U*_*s*_, a lineage will typically acquire its first generalist mutation before its first specialist mutation. This extra waiting time acts as an effective rolling fitness cost for specialist mutations of

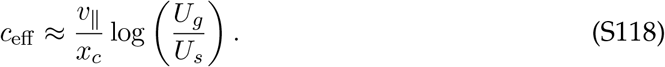

Because the effective fitness cost depends only logarithmically on the mutation rate ratio, modest differences in mutation rate (e.g., factors of 2) will have minimal impact on the population dynamics. Generalist mutations will only comprise the majority of the long-term optimal mutational path if

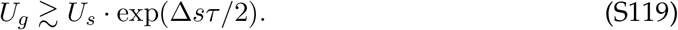

Note that this exponential factor corresponds to the specialist advantage in the rare-specialist regime. Thus, this transition represents the crossover between the frequent-specialist and the rare-specialist regime: as the specialist mutation rate increases, specialist mutations will begin to become dominant in the population when *U*_*s*_*p*_fix_(*s*, Δ*s*) ≳ *U*_*g*_*p*_fix_(*s*, 0).

#### 3.2.8 Stronger Tradeoffs and Specialist Bursting

When calculating the optimal mutational path using the update rules in Eq. (S61a) and Eq. (S61b), we assumed that *s >* Δ*s*, so that a mutant strain always grows faster than its parent strain regardless of the environmental state. In this regime, the members of a fitness class will mostly be composed of descendants of the first mutant into that class, so the optimal mutational path always entails acquiring a new mutation as soon as possible.

When Δ*s > s*, a new type of step in the optimal mutational path becomes relevant: a lineage may acquire *no* mutations for ~ *τ* ≫ *s/v*_∥_ generations, but rise to high enough frequencies that it can produce a “burst” of specialist mutations in rapid succession when the environmental state switches. While we will not provide a full summary of this regime in this work, we will derive the new optimal mutational path it implies, and its implications for some of the observables we have discussed in this section.

##### Rolling fitness of bursting lineages

Consider the process of a lineage acquiring specialist mutations that favor the upcoming environment, which (when Δ*s > s*) appear deleterious in the current environment state. The key effect is that these fitness classes will no longer be dominated by the *first* mutation into each class – instead, most of their members will be new mutations from the faster-growing parent class that occur right before the environment switches.

To describe this process, we can take advantage of the fact that stochasticity in the time each individual mutation arises becomes largely negligible, because the composition of each class will be determined by many independent mutational events. Consider a founding lineage at relative fitness (*x*_∥_, *x*_⊥_) in a constant environment *I*(*t*) = +1, with frequency

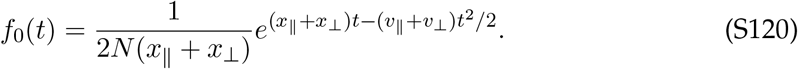

It founds successive classes via mutations that appear deleterious in the current environment, with fitness effect *α* ≡ Δ*s* − *s*. The dynamics of these classes are given by

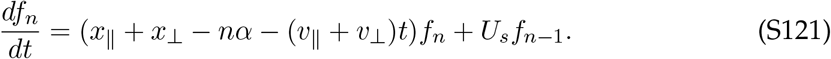

We will assume that *x*_∥_ + − *x*_⊥_ − *nα* (*v*_∥_ + *v*_⊥_)*t* remains positive (so new classes can still establish), and that enough time has passed for the focal class to contain several members (so stochasticity in founding mutations can be neglected). The solution to this equation can be written as an integral expression over all the possible times of each mutation,

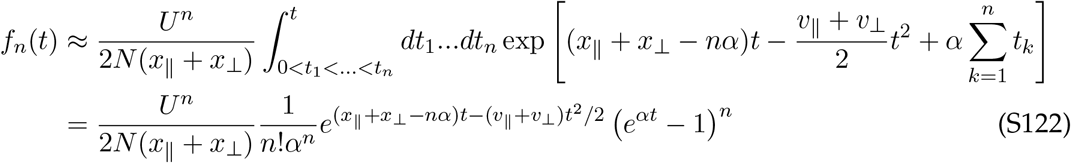

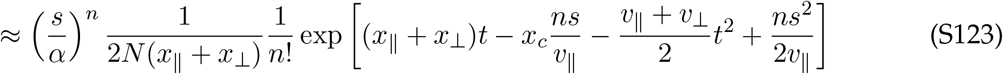

for *t* ≫ log(*n*)*/α*. (Loosely, we expect this to be valid when Δ*s* and *s* are not fine-tuned to be extremely close to each other.) Because the optimal times for each mutation to arise are shortly before the present, the frequency profile of the fitness class with *n* additional “deleterious” specialist mutations depends only algebraically on *α*.

Now, suppose the *n* + 1st mutation of this type lands in a beneficial environment, meaning that once again only the first mutation founded matters. This mutation will be founded near time *T*, defined via the integral relation

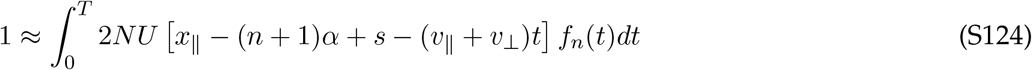

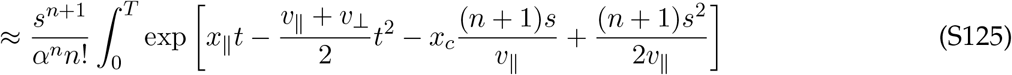

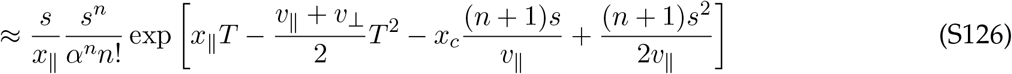

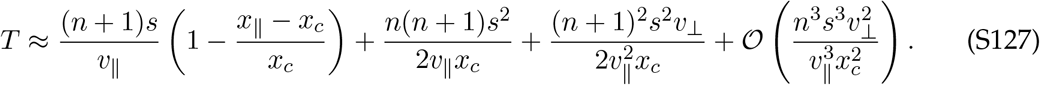

These dynamics mean that a lineage which acquires a burst of *n* mutations right when the environment switches will land at rolling fitness

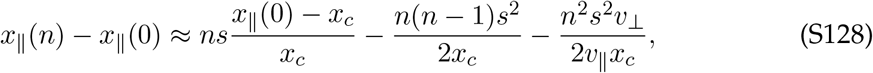

approximately independent of Δ*s*. Instead, the rolling fitness contains a cost of ~ *n*^2^*s*^2^*/x*_*c*_, due to the lineage failing to acquire *any* beneficial mutations during the waiting period of ~ *ns/v*_∥_ generations. However, this cost can be compensated by the benefit of acquiring *n* perfectly-timed specialist mutations, as we will discuss next.

##### Optimal mutational path

To derive the optimal mutational path, we will compare the effective fitness benefit of acquiring specialist mutations throughout an environmental cycle to the effective benefit of acquiring a burst of specialist mutations near the boundaries. Earlier, we found that an optimal specialist mutation which arises at time *t* in a cycle provides the same long-term fitness benefit as a generalist mutation with effect

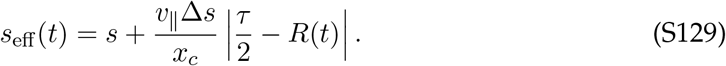

The *total* effect of acquiring successive specialist mutations between time *t*_0_ and *τ* (for 0 *< t*_0_ *< τ*) is

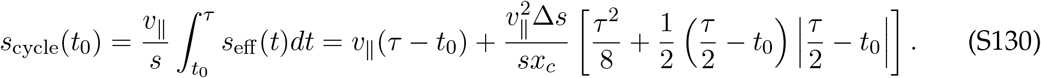

We must compare *s*_cycle_ to the total fitness effect of acquiring a burst of *n* = *v*_∥_(*τ* − *t*_0_)*/s* specialist mutations at time *τ*, after not acquiring any mutations for *τ* − *t*_0_ generations. This will include both the direct fitness effects of the mutations, their effective fitness boost from being perfectly-timed specialist mutations, and the cost of waiting to acquire them:

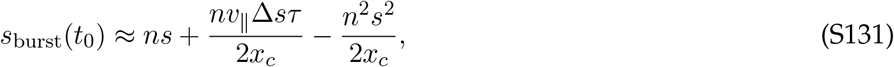

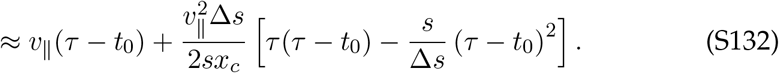

If *s >* Δ*s*, then *s*_cycle_ *> s*_burst_ for all *t*_0_, consistent with the non-bursting optimal path we analyzed earlier. When Δ*s >* 2*s*, then *s*_burst_ *> s*_cycle_, so the optimal path will *only* involve acquiring bursts of specialist mutations when the environment changes. In the crossover region 2*s >* Δ*s > s, s*_burst_ becomes larger than *s*_cycle_ at an intermediate value of *t*_0_ between 0 and *τ/*2, indicating that the optimal path entails acquiring some number of sequential specialist mutations aligned with the current environment, then waiting to acquire further mutations (of the opposite specialist type) in a burst when the environment switches.

Focusing on the burst-dominated Δ*s >* 2*s* regime, the effective fitness boost caused by following the optimal bursting path is roughly

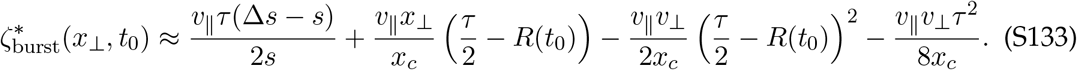

When Δ*s >* 2*s*, the leading constant term in Eq. (S133) is larger than the corresponding term in the *s >* Δ*s* regime, Eq. (S68), increasing the overall benefit of following the optimal mutational path. However, the overall scaling of this term is similar (changing by at most a factor of 2), so the correction it induces on the population rate of adaptation *v* will be qualitatively similar. The *x*_⊥_-dependent term is also identical, meaning the time-dependent advantage of specialist mutations remains the same.

##### Specialist advantage

An important qualitative difference between the Δ*s > s* bursting regime and the *s >* Δ*s* non-bursting regime is that when bursts dominate, successful specialist mutations become concentrated near the boundaries of environments, where their fitness advantage over generalists is largest. This means that after averaging over time, the overall specialist advantage 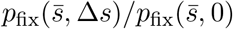 will be exponential in Δ*sτ*, as in the rare-specialists regime – but unlike the non-bursting regime in Eq. (S103). While we will not calculate the full time-dependent fixation probability, we roughly expect it to be between the rare-specialists and *s >* Δ*s* predictions. For comparison to simulations, we find that a reasonable approximation is to write a fixation probability identical to the rare-specialists case, with a leading correction coming from the constant shift in *ζ*^∗^ from Eq. (S105):

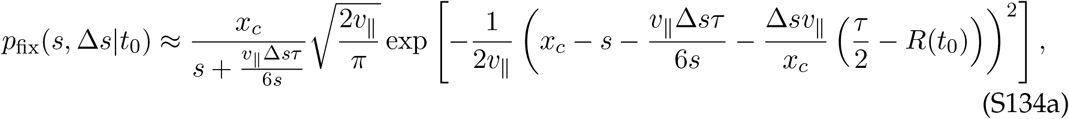

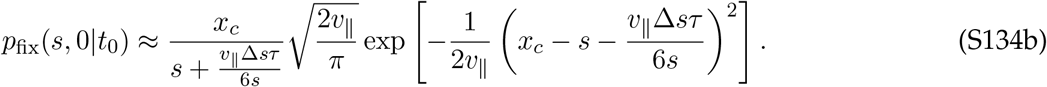

Averaging over time yields

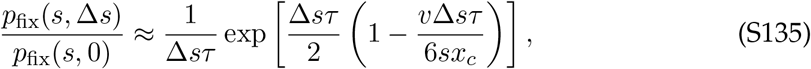

which we use to generate our theory curves for simulations where Δ*s > s*. We emphasize that the time-dependent effective fitness advantage of specialists, Eq. (S129), is the same between the two regimes – so a specialist mutation that arises near the boundaries of an environmental state is exponentially more likely to fix than a generalist, even when *s >* Δ*s*. The difference in overall fixation probability stems from the times that successful mutations typically arise: uniformly when *s >* Δ*s*, and concentrated near environmental switches when Δ*s > s*.

### 3.3 Frequent Specialists with Seasonal Drift

Finally, we will discuss the case of stochastic environmental fluctuations (*σ*_*τ*_ *>* 0) with frequent specialist mutations (*NU*_*s*_ ≫ 1). While the complexity of accounting for stochastic switching in a fully two-dimensional fitness landscape precludes us from finding a full solution in this work, we can nonetheless roughly estimate the influence of stochasticity on the population dynamics. Following our previous approach, we will track how stochastic environmental switches change the rolling fitness of a lineage, determining the population rate of adaptation and the specialist advantage.

The overall takeaway we will find is that environmental stochasticity reshapes the optimal mutational path to become aligned with upcoming environmental change. This magnifies the advantage of specialist mutations relative to the case without stochasticity. However, the scale of this additional boost is much less dramatic than we observed for rare specialist mutations (SI 2.2.1). This is because when specialist mutations are widely available, the dynamics of all lineages will be subject to random environmental fluctuations, and the relative benefit provided by a single specialist mutation decreases. Nonetheless, environmental stochasticity will still have significant impacts on the fixation probability of mutations and the population rate of adaptation.

We will continue to employ our ansatz in Eq. (S56) expressing the adaptation of the bulk population in terms of *v*_∥_ and *v*_⊥_. This is a strong approximation, because the bulk rate of specialization *v*_⊥_ over a given environmental epoch may depend on the realized stochasticity, rather than being described by a single constant (Fig. S10A). Nonetheless, this is an initial linear approximation which reflects the fact that the bulk population will adapt more toward environments that happen to last longer.

#### 3.3.1 Faster Environmental Cycles

In a stochastic environment, it is convenient to focus on the case of faster environmental cycling, *τ* ≪ *s/v*_∥_, so that many environmental cycles occur in the time it takes for a lineage to found a new mutation. First, we will calculate the impact that environmental stochasticity has over *s/v*_∥_ generations, the time it takes to found a single mutation. We can then extend this result over many foundings to determine how seasonal drift influences the fate of a mutation. (We will ignore the deterministic effects discussed in the previous section, assuming their contribution is small when stochastic effects dominate – this is more reasonable in the smaller-*τ* regime.)

Consider a lineage at initial frequency *f*_0_, average fitness *x*_∥_, and environmental preference *x*_⊥_ in an environment aligned with the +*x*_⊥_ direction. After *t*_1_ generations in a single environment, its frequency will be

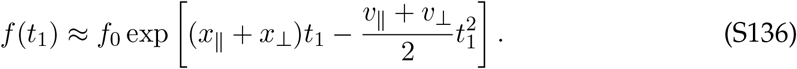

Then, suppose the environment switches states for *t*_2_ generations. As before, we will neglect stochasticity in the total environmental cycle time and focus on the time difference between environments, assuming that *t*_1_ + *t*_2_ = 2*τ* and defining *δτ* ≡ *t*_1_ − *t*_2_. The final frequency of the lineage after a full cycle will be

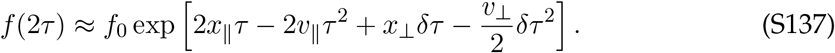

Each environmental cycle will have a new stochastic value of *δτ*, perturbing the frequency of the lineage. After *n* such cycles, the frequency of the lineage will be

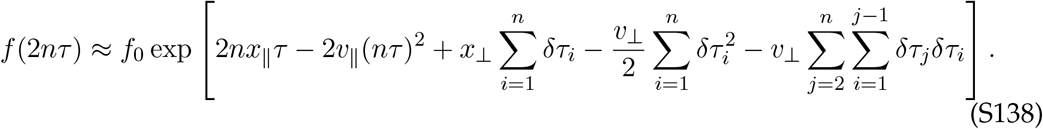

A newly established lineage must undergo *n* ≈ *s/*2*v*_∥_*τ* such cycles in order to found a new beneficial mutation. The net effect of the stochasticity over this time will determine the lineage’s rolling fitness after the *n* cycles are complete. We can use this to define new recursive relations where each step represents the founding of a new mutation, in terms of *δx* ≡ *x*_∥_ − *x*_*c*_:

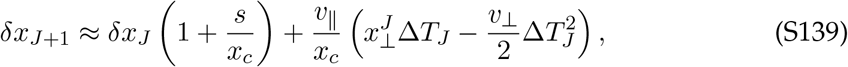

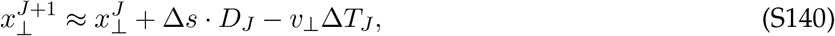

where *D*_*J*_ is the environmental preference of the *J* th mutation, and

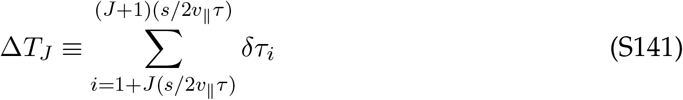

is the net environmental stochasticity over the time period during which the focal lineage founds its *J* + 1st mutation. The solution to these recursion relations is

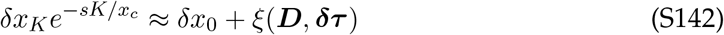

with

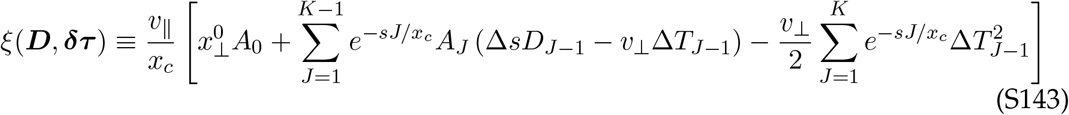

an effective shift in the starting fitness, and

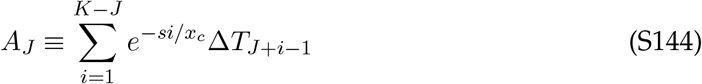

a weighted sum of the environmental stochasticity after the time indexed by *J*. We observe that the optimal mutational path is

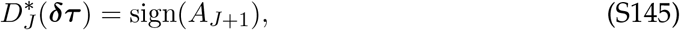

such that optimally specializing lineages should “anticipate” the upcoming environmental bias. The survival of the focal lineage depends on whether

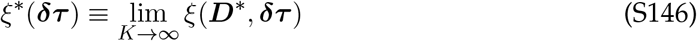

is greater than −*δx*_0_, meaning that the probability of survival will strongly depend on the distribution of *ξ*^∗^, and in particular its positive tail under the optimal mutational path.

##### Characterizing the stochasticity

The distribution of *ξ*^∗^ is complicated due to the strong correlations between the *A*_*J*_, so we will generally characterize it through semi-analytic approximations based on fitting to empirical samples. Note that *ξ*^∗^ only depends on environmental stochasticity and not the dynamics of mutations, so its distribution can be sampled straightforwardly by drawing a set of epoch times.

To leading order in *s/x*_*c*_, the mean and variance of *ξ*^∗^ are

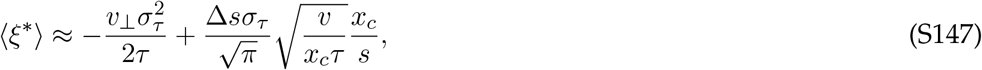

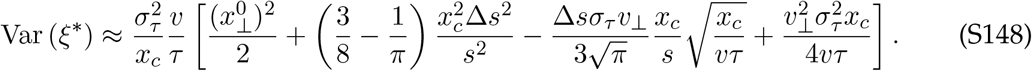

However, the most relevant quantity for calculating the influence of *ξ*^∗^ on fixation is its survival function *S*(*z*) in the positive tail, indicating the effective fitness benefit it provides given lucky environmental stochasticity. We find that a reasonable phenomenological fit to this survival function over a range of parameters is a Gaussian which approaches a cutoff at some scale *ξ*_max_ (Fig. S8A):

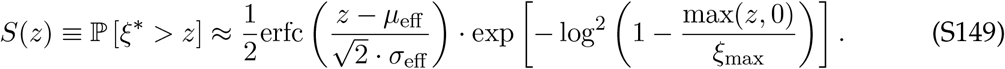

To reduce the number of free fitting parameters, we approximate *µ*_eff_ = ⟨*ξ*^∗^⟩ from the analytic expression in Eq. (S147). Furthermore, Eq. (S143) shows that *ξ*^∗^ is a quadratic form in the Δ*T*, suggesting that the cutoff can be expressed as the maximum over the Δ*T*:

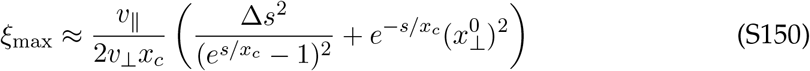

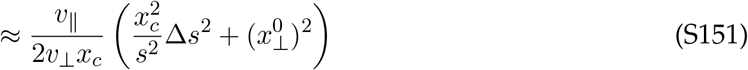

This leaves the Gaussian spread *σ*_eff_ as the only free fitting parameter for the survival function at a particular combination of variables. The analytic dependence of *σ*_eff_ on the evolutionary parameters and initial specialism 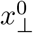 does not admit an obvious form, because *σ*_eff_ represents not the total variance in *ξ*^∗^, but the variance in the positive tail (before the cutoff becomes relevant). Thus, we often use the fit values of *σ*_eff_ directly. Nonetheless, when a semianalytic form os *σ*_eff_ is necessary, we find that for relatively small 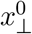 it can roughly be described by a quadratic ansatz (Fig. S8B):

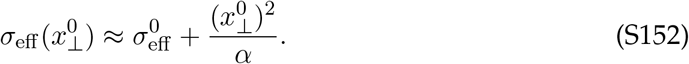

The intercept 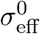 appears approximately linear in Δ*s*, suggesting it is dominated by the second term in the brackets in Eq. (S148):

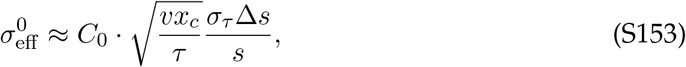

with *C*_0_ a numerical fitting constant accounting for the effective scaling of the variance in the positive tail (Fig. S8C). A quadratic dependence on 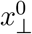 is consistent with a Taylor expansion of Std(*ξ*^∗^), suggesting a scaling of

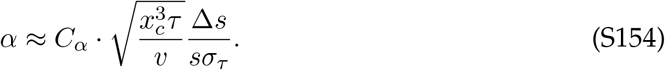

We emphasize that these are rough phenomenological fits for the parameter range used in simulations, meant to broadly characterize the distribution of *ξ*^∗^. However, they are sufficient to illustrate the key qualitative upshot, which is how the tail of *ξ*^∗^ depends on the initial specialism 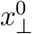 through *σ*_eff_ and *ξ*_max_.

##### Fixation probabilities

Following our previous approach in Eq. (S46), we approximate the fixation probability as a double integral over the background mean fitness *x*_∥_ and specialism *x*_⊥_:

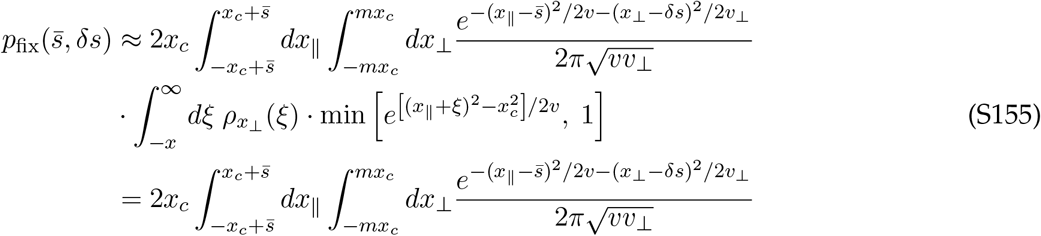

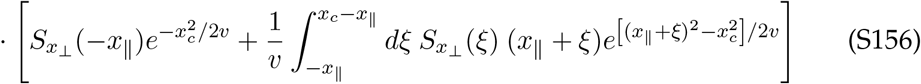

with 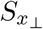 the survival function of *ξ*^∗^ at a given *x*_⊥_. Here, *m* is a constant setting the integration bounds of *x*_⊥_. Our theory assumes that |*x*_⊥_| *< x*_*c*_ so that a new mutation can escape genetic drift regardless of the environmental state, but allowing the integral to extend past this bound by setting *m* = 3 allows for better agreement with simulations.

Our approximations are not sufficient to recover the precise form of *p*_fix_, but they do accurately predict the specialist advantage *p*_fix_(*s*, Δ*s*)*/p*_fix_(*s*, 0) at small stochasticity (Fig. S7B). This advantage comes from the fact that if the focal mutation has a tradeoff Δ*s >* 0, the initial lineage can attain a slightly higher value of specialism |*x*_⊥_|, which widens the survival function of *ξ* (Fig. S8A). However, this shift in the survival function is relatively modest compared to the rare-specialists case, where the focal specialist was the *only* mutation making a lineage sensitive to environmental change.

To produce the theory curves in Fig. S7B, we numerically integrate Eq. (S156) using values of *σ*_eff_ (*x*_⊥_) inferred from the survival function of *ξ*^∗^. This approximation fails at the larger values of Δ*s* we study in Fig. 4C, likely due to to |*x*_⊥_| becoming large enough relative to *x*_*c*_. To get more intuition for the scaling at small Δ*s*, we can assume cutoff effects are negligible. In this case, the calculation proceeds identically to the rare-specialists case until Eq. (S49), except *σ*_*ξ*_ is now replaced by *σ*_eff_ (*x*_⊥_):

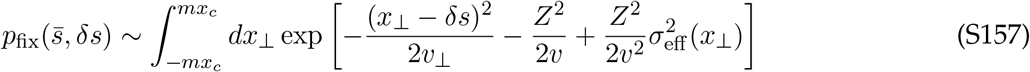

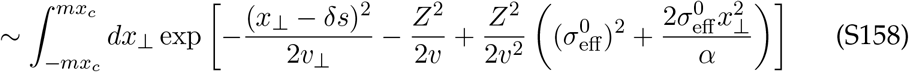

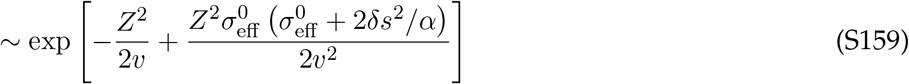

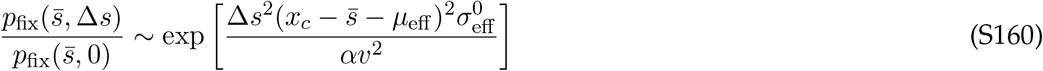

with 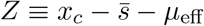. Plugging in our rough scaling forms for 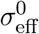 and *α* yields

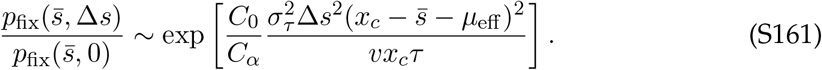

This scaling is similar to the rare-stochasticity scaling in Eq. (S49), up to an exponential scaling prefactor of *C*_0_*/C*_*α*_ ≈ 0.1 and the *µ*_eff_ shift in the exponent. However, these approximations assume the cutoff is negligible and that stochasticity is weak enough to be treated as a small perturbation, 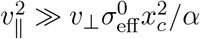. These assumptions fail above Δ*s* ≳ 0.015 for our parameters, suppressing the specialist advantage (Fig. S7B).

To estimate the specialist advantage over a wider range of Δ*s* and produce the theory curves in Fig. 4C, we use a simple ansatz for the fixation probability representing how typical values of environmental stochasticity *ξ*^∗^ produce an effective fitness shift. To approximate this shift, we assume the term in Var(*ξ*^∗^) corresponding to Eq. (S153) dominates the variance:

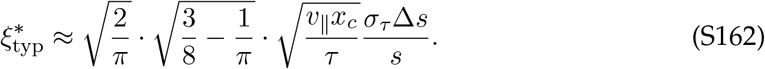

This shift comes from each of the specialist mutations a lineage acquires on its way to fixation. A single specialist mutation contributes a fraction ~*s/x*_*c*_ of this shift. So, the fixation probability of each type of mutation will be roughly

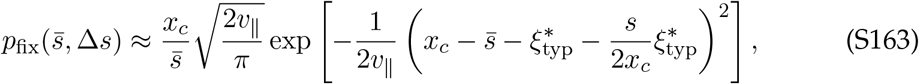

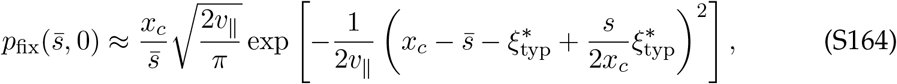

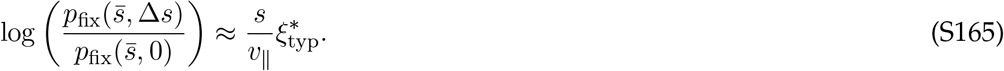

This result broadly agrees with the low-Δ*s* results for our parameter values (Fig. S7B) and has better agreement with simulations for larger Δ*s*, so we use it as a rough comparison to simulations.

##### Rate of adaptation

To estimate the influence of stochasticity on the rate of adaptation *v*_∥_, we employ the simple ansatz above and solve for *v*_∥_ self-consistently:

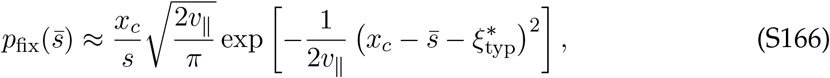

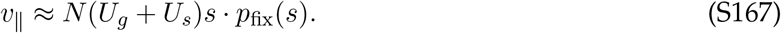

Numerically solving Eq. (S167) yields a reasonably accurate solution for *v*_∥_ over the entire range of Δ*s* we study. To leading order, the correction is

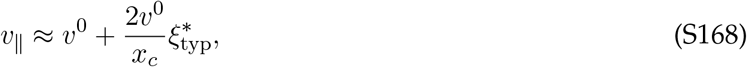

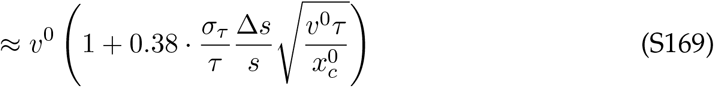

While this adjustment is nominally small, it can be large enough to increase *v*^0^ by about a factor of two for the values of Δ*s* and *σ*_*τ*_ we use in simulations.

#### 3.3.2 Slower Environmental Cycles

When *τ* ≫ *s/v*_∥_, we can take essentially the same approximate approach to characterize the population dynamics as in the faster-cycling *τ* ≪ *s/v*_∥_ regime, but with rescaled effective parameters. In the slower-cycling regime, a lineage can acquire many specialist mutations in a single environmental cycle. However, we expect the optimal mutational path to include long sequences of the same type of specialist mutation, since when *σ*_*τ*_ is reasonably large, the optimal mutations are mostly set by future stochasticity. Under this approximation, we assume that our focal lineage acquires only one type of mutation during each environmental cycle, for a total of 2*vτ/s* specialist mutations.

The rolling fitness update rule corresponding to Eq. (S139) is then

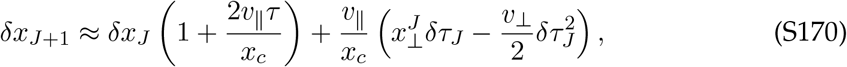

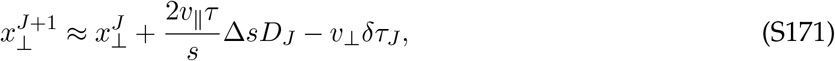

where *J* indices represent complete environmental cycles. These update rules can be roughly mapped to Eq. (S139), but with *s* replaced by 2*v*_∥_*τ*, and Δ*s* rescaled by an additional factor of 2*v*_∥_*τ/s*. This transformation preserves the ratio Δ*s/s*, which allows us to apply the results from the faster-cycling regime directly.

Our results are somewhat less accurate than in the smaller-*τ* regime (Fig. 4C, top). One possible reason for this discrepancy is that when multiple new fitness classes arise each environmental cycle, the optimal mutational path will be a mix of the periodic-environment and stochastic-environment contributions. Here, we have neglected these details and focused on the stochastic contribution only.

Regardless of the precise specialist advantage, the key takeaway from this section is that all the specialist mutations a lineage acquires on its way to fixation or extinction make its trajectory highly sensitive to environmental stochasticity. Successful lineages will acquire mutations which benefit from environmental stochasticity, acting like a deterministic fitness benefit that speeds up the overall rate of adaptation of the population. However, because all lineages can acquire specialist mutations, the advantage of a single specialist mutation is attenuated relative to the rare-specialists regime, where only one lineage experiences environmental stochasticity. A more precise analysis of this stochastic regime is a compelling avenue for future work.

#### 3.3.3 Rate of Specialization

The last unknown parameter we have not calculated is the rate of specialization *v*_⊥_, which enters in the specialist fixation probability. We can estimate this quantity using a procedure similar to the deterministic case in SI 3.2.3, by tracking how the net specialism a lineage accrues depends on environmental stochasticity (Fig. S10A).

Once again, we define *D*_tot_(*n*) as the sum of the mutations an optimally-specializing lineage has accrued just before founding its *k*th mutation. These mutations will depend on the stochasticity according to Eq. (S145):

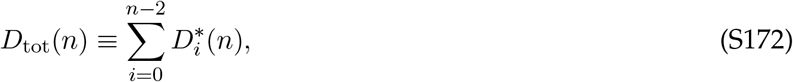

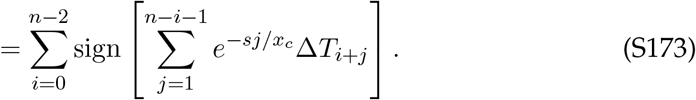

The rate of specialization *v*_⊥_ encodes how sensitive *D*_tot_(*n*) is to the environmental bias just before observation, Δ*T*_*n*−1_. This environmental bias may have an influence on the optimal 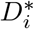 up to a coalescence time ago, *n* ≈ *x*_*c*_*/s*. Under our approximation of a constant *v*_⊥_, we therefore expect *D*_tot_(*n*) to be an approximately linear function of Δ*T*_*n*−1_ for typical values of Δ*T*. We will find this linear slope by first writing out the dependence explicitly:

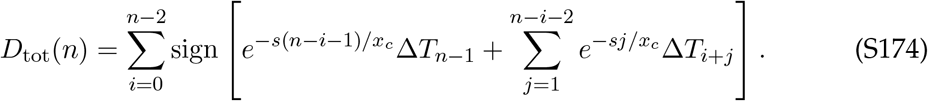

We will treat Δ*T*_*n*−1_ as a fixed bias parameter, while marginalizing over the distributions of all the other Δ*T*_*i*_. Approximating them as i.i.d. Gaussians with width 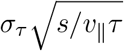, we can write the expectation of *D*_tot_ in terms of the error function:

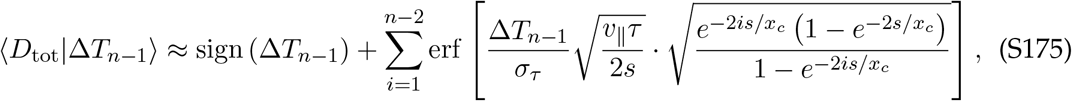

re-indexing *i* so smaller *i* are closer to the present at *k*. Smaller *i* closer to the present are more likely to be influenced by the direction of Δ*T*_*k*−1_, while Δ*T*_*k*−1_ has a relatively small influence on terms with large *i* further in the past.

To obtain the linearized behavior for small Δ*T*_*n*−1_ and *i* ≲ *x*_*c*_*/s*, we can expand the exponents for *s/x*_*c*_ ≪ 1 and the error function for small arguments. Dropping the trivial sign term, we find

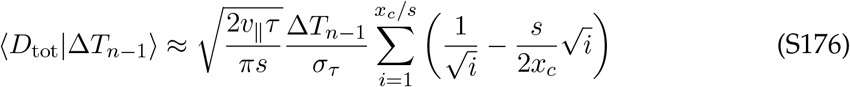

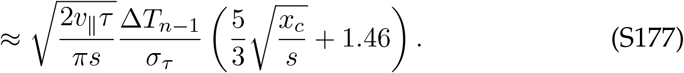

The rate of specialization is simply the linear coefficient of Δ*T*_*n*−1_ times Δ*s*:

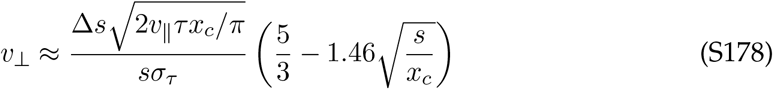

Notably, the rate of specialization *declines* with increasing stochasticity, and is markedly less than the periodic case (Fig. S10B). Intuitively, this is because more periodic environments allow for more standing diversity in specialization levels, while stochastic environments will lose some diversity when the environment becomes biased to one state for a period of time.

The same argument made in SI 3.3.2 holds when calculating *v*_⊥_ in the slower-cycling *τ* ≫ *s/v*_∥_ regime. Replacing *s* with 2*v*_∥_*τ* and Δ*s* with Δ*s* · 2*v*_∥_*τ/s* leaves Eq. (S178) unchanged to leading order, but adjusts the correction term:

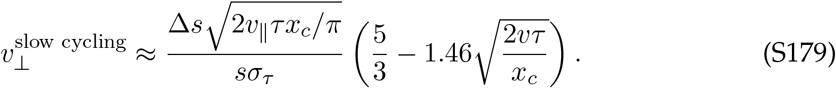

As in the periodic-cycling regime, these expressions for *v*_⊥_ are not fully accurate in the regime studied in simulations, but are largely correct for small Δ*s* and predict the correct qualitative dependence (Fig. S10B).

## 4 Simulated Population Dynamics

### 4.1 Overview

Throughout this work, we compare our theoretical predictions to results inferred from simulated Wright-Fisher population dynamics. It is convenient to represent the population as a three-dimensional matrix (one dimension for each type of mutation), where each entry represents the number of individuals with that specific combination of mutations. The basic procedure is as follows:

1. Initialize a monoclonal population of size *N* with no mutations in an arbitrary environmental state. Draw the time before the next environmental switch from the distribution Gamma(*τ* ^2^*/*2Δ*τ* ^2^, Δ*τ* ^2^*/τ*).
2. Calculate the absolute fitness *λ*_*i*_ of each mutation class, based on the current environmental state.
3. Perform a selection step by setting 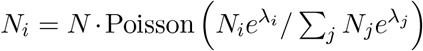. This roughly corresponds to a generation of time.
4. Sample the number of mutations of each type within each mutation class as Binomial(*N*_*i*_, *U*) with *U* = *U*_*g*_ (for generalists) or *U*_*s*_*/*2 (for each type of generalist). Update *N*_*i*_ accordingly, extending the population matrix to include a new mutation class if necessary.
5. Increment the time by 1 and check if the new time is after the next environmental switching time. If it is, switch the environmental state and sample the next waiting time as in Step 1.
6. Record the simulation state and remove empty slices of the population matrix. Then, return to Step 2 and repeat until the total simulation time is complete.

Because we often simulate large population sizes of *O*(10^20^), the size of sufficiently large mutation classes are stored as floating-point numbers rather than integers. To ensure numerical stability, we use approximations to the above probability distributions when their arguments are sufficiently large. In particular, we approximate the binomial distribution as Poisson when *N*_*i*_ *>* 10^5^ and *N*_*i*_*U* (1 − *U*) *<* 10, and as a normal distribution when *N*_*i*_ *>* 10^5^ and *N*_*i*_*U* (1 − *U*) *>* 10. We approximate the Poisson distribution as a normal distribution when its mean is greater than 10^5^.

Fixation probabilities can be inferred from this distribution by tracking the average number of each mutation type in the population over time and performing a linear fit. The total number of mutations in the population scales with *t* · *NUp*_fix_ for long times, so dividing the slope of the fit by *N* and the mutation rate yields the fixation probability.

### 4.2 Test Lineages

While the above procedure accurately simulates the model of interest, keeping track of mutation classes rather than individual lineages obscures how fixation probabilities depend on other stochastic factors, such as the time a mutation establishes or the background it appears on in the population. Furthermore, when we consider rare specialist mutations, we would like to consider situations when only one specialist mutation is present in the popoulation at a time. We can remedy both of these issues by introducing “test lineages” in the simulation.

Test lineages are parallel population matrices descended from the main population, which interact with each other and the main population just like any other individual, but are kept track of separately in numerics. If a test lineage dies or reaches fixation, a new test lineage is introduced, so that there is at most one test lineage in the population at any given time (in the case that the previous test lineage fixed, it becomes the new background population). When specialist mutations are rare, we can set *U*_*s*_ = 0 for the above procedure and treat the test lineages as their own “specialist” class, with the introduction of a test lineage effectively being a rare specialist mutation event. When specialist mutations are common, both the background population and the test lineage are full three-dimensional matrices free to explore mutational space.

In order to accurately predict fixation probabilities, test lineages should be introduced into the population in a manner which is statistically indistinguishable from how specialist mutations already arise. Specifically, they should start as single individuals, with a mutational background selected proportionally to its frequency in the population. However, most random mutations in the population will not reach fixation, meaning that almost all test lineages introduced this way will soon die. In order to more efficiently estimate fixation probabilities from test lineages, we implement a method of importance sampling which makes test lineages more likely to arise on rare backgrounds, and less likely to die due to genetic drift immediately after being introduced. We can then correct for this importance sampling when estimating the overall fixation probability.

The procedure for introducing test lineages is summarized as follows, which occurs directly after the mutation step above (Step 4):

1. If there are no test lineages in the population, set a time for the next one to appear.
  a. If Δ*τ* = 0, choose this time uniformly over the next complete environmental cycle (starting from the next environmental switch time).
  b. If Δ*τ >* 0, first calculate the probability the test lineage arises during the next cycle as *P*[Expon(*τ*) *< T*_next_], where *T*_next_ is the duration of the next environmental cycle. If it is slated to arise in the next cycle, sample the time uniformly as in (a). (This decreases the probability the test lineage arises during abnormally short cycles.)
2. If it is time for a test lineage to arise, initialize it.
  a. Choose the mutational background it arises on *uniformly* from all possible backgrounds (i.e. nonzero entries of the population matrix). This makes all backgrounds equally likely, regardless of their frequencies in the population. Save *w* ≡ *n*_bg_*f*_*i*_, where *n*_bg_ is the number of possible backgrounds and *f*_*i*_ is the frequency of the chosen background in the population. This is an importance sampling reweighting factor which we will need later.
  b. The test lineage will have mutations corresponding to its chosen background, plus an additional (random]) specialist mutation. Set its initial number as *N*_start_ ≡ Max [1, (2(*s* + Δ*s*)*n*_muts_)^−1^], where *n*_muts_ is the total number of mutations the new test lineage has (across all types). This allows the test lineage to start at a frequency above 1*/N*, but still in the regime dominated by genetic drift.
  c. dSet *w* = *w/N*_start_ to account for the probability a newly introduced lineage will drift to *N*_start_ rather than going extinct.
  d. Record the time the lineage was initialized, its background, its reweighting factor *w*, and any other desired information.
3. If a test lineage is present, it follows the same selection and mutation steps as any other individual. Descendants of the test lineage remain in the test lineage, so the frequency of the test lineage represents the frequency of the new specialist mutant it introduced in the population.
4. If the test lineage reaches fixation or dies, record its fate. If it reached fixation, the background population will be at zero frequency, so set the test lineage as the new background population and start from Step 1.

The true fixation probability can be estimated from a collection of *M* test lineages:

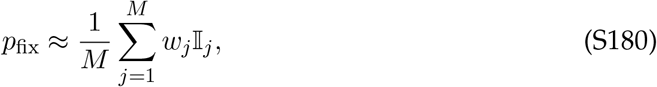

where I_*j*_ = 1 if the *j*th test lineage reached fixation, and 0 if it went extinct. The uncertainty of this estimate is roughly the standard error of the mean among the *w*_*j*_I_*j*_. In practice, this estimate of *p*_fix_ is noisier than a linear fit across frequent mutations, but allows for reasonably accurate estimations of extremely low values of *p*_fix_ ~ 10^−15^ from only ~ 10^5^ test lineages.

### 4.3 Fixation Probability Isoclines

In Fig. 6, we plot fixation probability isoclines, curves of constant fixation probability in two-dimensional fitness effect space. To obtain the simulation points corresponding to these curves, we empirically measured the fixation probability of “test charge” mutations with fitness effects (*s* − *c*, Δ*s*) for a range of values of *c* and Δ*s*. We then compared this probability to the fixation probability of generalist mutations with fitness effect (*s*, 0), fitting a line to their log ratio to estimate the critical value *c*^∗^ at which the fixation probabilities are equal. Repeating this procedure over a range of Δ*s* produced the empirical fixation probability isoclines.

The theoretical predictions were derived from our previous results for *p*_fix_(*s, δs*), with minor modifications. For rare specialists in a stochastic environment, cutting off the stochasticity at 3 standard deviations as in Fig. S1 produced the most accurate results, and the isoclines were extended linearly past the region where our theoretical prediction for *p*_fix_ broke down. For frequent specialists in a stochastic environment, the deterministic contribution to the effective fitness shift was comparable to the stochastic contribution, so we added them together to produce the stochastic-environment isoclines.

